# Climate-driven shifts in distributions and habitat suitability of amphipod species under future ocean change

**DOI:** 10.64898/2026.04.13.718119

**Authors:** Farzaneh Momtazi, Hanieh Saeedi

## Abstract

Concern about how climate change affects marine ecosystems is growing, despite international commitments to reduce CO2 emissions. Predicting amphipod species responses to ocean warming is critical due to their high abundance and key ecological role in marine ecosystems. We selected 35 widespread benthic amphipod species with at least 30 unique occurrence records after thinning from two or more biogeographical regions and classified them according to depth and feeding strategy. Following spatial thinning, 17 species retained sufficient occurrence records for Maximum Entropy (MaxEnt) modelling, of which 15 met the model evaluation criteria by having a Partial ROC value below 1 or a 5% omission rate exceeding 0.2. We projected species distributions under the low emission RCP 2.6 and high emission RCP 8.5 scenarios for 2050 and 2100. To compare species responses among feeding groups, we used linear mixed effects models with feeding type, scenario_time combination, and their interaction as fixed effects and species identity as a random effect. Species were also classified by depth, but statistical comparisons among depth groups were not performed because of limited species representation. Projected distributions showed substantial species-specific distribution shifts, including both gains and losses of suitable habitat and changes in areas of high species richness. Linear mixed effects models showed that potential future changes in suitable habitat area did not differ significantly among feeding groups, whereas centroid shifts were significantly influenced by the interaction between feeding type and scenario–time combinations. This indicates that trophic strategy influences the spatial response of amphipods to future climate change. These findings highlight that climate change may dramatically alter the functional composition of benthic communities and their ecological roles, beyond simple changes in species distributions. Incorporating trophic identity and functional roles into climate impact assessments will be essential for predicting ecosystem responses and informing conservation strategies that safeguard marine ecosystems functioning under future climate change. This approach will improve predictions of ecosystem responses and strengthen conservation and management strategies aimed at maintaining ecosystem functioning in a rapidly changing ocean.

## 1. Introduction

Climate change’s impact on biodiversity loss is recognized as one of five major direct drivers, along with land/sea use change, overexploitation, invasive species, and pollution in the global assessment report from the Intergovernmental Science-Policy Platform on Biodiversity and Ecosystem Services (IPBES) (Díaz et al., 2019). The impact of climate change on marine ecosystems is proven in distribution patterns and biodiversity, driven mainly by ocean warming, acidification, deoxygenation, and sea-level rise (Wabnitz et al., 2018). Recent studies reported that uneven species responses to climate change are highly variable across taxa and regions, including geographic range shifts, phenological shifts, physiological and morphological responses, behavioral plasticity, genetic responses, and finally extinction (Crowther & Schwanz, 2025; Radchuk et al., 2019). Such changes extend beyond individual species and may lead to the reorganization of ecological communities and their functional roles. Within the planetary boundaries framework, climate-driven biodiversity change therefore represents an important link between the climate change and biosphere integrity boundaries, highlighting how continued environmental change can affect not only the distribution and persistence of species but also the structure, functioning, and resilience of ecosystems (Richardson et al., 2023; Rockström et al., 2009).

Climate change affects marine trophic levels unevenly, while lower trophic levels, such as producers and herbivores, often experience stronger declines. Their losses can then spread through the food web, becoming stronger at higher levels such as carnivores and top predators (Guibourd de Luzinais et al., 2023). Negative climate effects will be about 1.5–2 times stronger from producers to predators, as each level loses more energy than it receives. It should be considered that warming accelerates this by raising metabolic demands faster than productivity rises (Ullah et al., 2018). Higher-level organisms tend to be more tolerant overall, likely because they are larger and more flexible in their diet and behavior (Hu et al., 2024).

Amphipods have key roles in marine ecosystems, serving as key links in food webs, nutrient cycling, and habitat engineering (Jażdżewska & Siciński, 2017; Minutoli et al., 2025). Their direct development and high abundance enable detection of phenological shifts, range changes, and adaptation in real time (Martinez-Alarcón et al., 2024). In another aspect, as ectotherms with narrow thermal tolerances, amphipods show rapid, measurable responses to warming, acidification, deoxygenation, salinity shifts, and induced physiological changes, such as reduced growth, reproduction, or survival, which often precede community-level effects (Kürzel et al., 2025; Lörz et al., 2022). Endemic and cold-water amphipods are considered early warning bioindicators for vulnerable systems (Huang et al., 2024). Amphipods, as subtropical herbivores and micro-predators that bridge primary producers (phytoplankton, algae) and higher carnivores in marine food webs, along with their limited dispersal ability, are expected to show different responses to climate change.

To predict potential species responses to climate change, Species Distribution Models (SDMs) are powerful statistical tools that correlate current distributions with environmental variables and project these onto future climate scenarios (Karp et al., 2025; Liu et al., 2025). The surveying response of amphipod distribution to climate change is rare and regional. Recently, in the North Atlantic (Kürzel et al., 2025) and the North Pacific (Kaiser et al., 2026),have provided initial evidence that amphipod and other benthic peracarid species may respond differently to future climate change. In the North Atlantic, Kürzel et al. (2025) showed species-specific responses, with some deep-sea amphipods projected to lose suitable habitat while others expanded their potential distributions or shifted northwards and towards the Greenlandic coast. Similarly, Kaiser et al. (2026) found that most benthic peracarid species in the northern North Pacific were projected to shift northwards, although the direction and magnitude of these shifts varied among species and were influenced by ecological traits such as diet and bathymetric distribution. Together, these studies suggest that climate change may lead to complex and species-specific redistributions rather than uniform responses among benthic taxa. Nevertheless, it remains unclear how these responses vary across a broader range of amphipod species representing different feeding strategies and depth preferences at a global scale. Addressing this knowledge gap is important for improving our understanding of how climate change may reshape the distribution and functional composition of benthic communities.

The previous investigation of the global distribution pattern of amphipods and the main environmental drivers revealed the importance of temperature in shaping shallow-water distribution patterns and the combination of environmental variables in deep-sea communities (Momtazi & Saeedi, 2024; Saeedi, 2026). Therefore, in the present study, we have applied species distribution modeling techniques to reveal how shallow-water and deep-sea amphipod species respond to future climate change scenarios under different Shared Socioeconomic Pathways (RCPs). The study has been restricted to benthic species that are more tightly coupled to relatively stable environmental gradients (e. g., temperature, substrate, and depth), whereas pelagic species experience highly dynamic and advective environments that are more difficult to represent with static or climatological predictors (Kaiser et al., 2026). We aim to answer: 1) How will amphipod species respond to different climate scenarios, RCPs, including RCP 1–2. 2.6 (sustainability-focused future with rapid reductions in greenhouse gas emissions, leading to a radiative forcing of ∼2. 2.6 W m□² by 2100 and limiting global warming to around 1. 1.5–2 ° C), and RCP 5–8. 8.5 (fossil-fuel-intensive future with continued increases in greenhouse gas emissions, resulting in radiative forcing of ∼8. 8.5 W m□² by 2100 and high levels of global warming) in 2050 and 2100? 2) Are trophic traits (feeding groups) predictive of species-specific vulnerability or resilience to climate-driven habitat shifts in benthic amphipods? We hypothesize that amphipod species will exhibit species-specific contrasting responses to climate change. We also hypothesize that the future habitat suitability of amphipod species will vary depending on their trophic group.

## 2. Materials and Methods

### 2.1 Species selection criteria

An initial set of 35 widespread benthic amphipod species representing major feeding groups was selected to evaluate potential distributional shifts under future climate change scenarios (Table S1). Species selection was based on the following criteria:

1. high abundance according to the most recent global assessment of amphipod distribution patterns, with more than 640 occurrence records based on Momtazi & Saeedi (2024);
2. a strictly benthic lifestyle, to avoid uncertainties associated with modeling the three-dimensional structure of the pelagic environment (Kaiser et al., 2026); and
3. Choosing widespread and well-sampled benthic amphipod species by means of occurrence records in at least two major biogeographic regions (temperate, tropical, or polar). Highly endemic or geographically restricted species were not included because the objective of this study was to assess climate-driven changes in the distribution patterns of widespread amphipod species at a global scale. For narrowly distributed species, projecting habitat suitability across the entire global ocean may identify environmentally suitable areas that are geographically distant from their known ranges and may not be accessible because of dispersal limitations or other biogeographic constraints. Thus, such projections could overestimate the potential future distribution of range-restricted species. Focusing on widespread species allowed us to evaluate potential distributional shifts within a spatial framework that was more consistent with their broad geographic ranges and the global scope of the study.
4. Species were assigned to their main feeding group. Species with at least 80% of their occurrence records at depths greater than 200 m were classified as deep-sea species.

Following spatial thinning of occurrence records, only species with more than 30 unique occurrence records were retained for modeling to ensure reliable MaxEnt model performance (Simões et al., 2021). As a result, 17 species were selected for depicting their distribution modeling by the MaxEnt model (Table S1).

### 2.2 Distribution records

Occurrence records were compiled from our previous global assessment of amphipod distribution patterns (Momtazi & Saeedi, 2024). All occurrence records were subjected to quality control following the protocol described in Saeedi et al. (2019). Taxonomic names were standardized against the World Register of Marine Species (WoRMS), and synonyms were resolved using the R package “obistools” (Provoost & Bosch, 2020). Occurrence records lacking geographic coordinates, duplicates, fossil occurrences, coordinate uncertainty >100 km, and without depth information were removed. Additionally, occurrences collected outside the period 2000–2020 were excluded to ensure temporal consistency with the Bio-ORACLE environmental predictors. In total, 3,353 occurrence records were retained after initial cleaning. The geographical position of amphipod records (after data cleaning) is shown in Figure S1. Following the approach of Simões et al. (2021), occurrence records were spatially rarefied (thinned). After thinning, only species with at least 30 occurrence records and a spatial distribution representative of their known range were retained (Table S1, Figs S1-17). The complete occurrence dataset for each species was randomly partitioned into training (75%) and testing (25%) subsets. A random partitioning strategy was adopted because no independent datasets were available for model evaluation. Data cleaning and processing were conducted in R version 3.5.1 (R Core Team, 2018), using the packages “spThin” (Aiello-Lammens et al., 2015), “raster” (Hijmans, 2015), and “rgdal” (Bivand et al., 2015).

### 2.3. Environmental data

Environmental data at 5 arcminutes were extracted from Bio-ORACLE (Assis et al., 2024; Tyberghein et al., 2012) for the benthic layer and average depths. Current environmental data covered the period 2010–2020, while future projections utilized the periods 2040–2050 and 2090–2100 to reveal the response in both medium- and long-term (In the Bio-ORACLE database, the end-of-century layers are labeled as 2090, representing climatic conditions for the late twenty-first century). Future environmental data from Bio-ORACLE included modelled variables based on RCPs, representing greenhouse gas emission scenarios under different policy decisions. This study used two scenarios: RCP 2.6 (low emissions, sustainable future) and RCP 8.5 (high emissions, worst case) (Lee et al., 2023). The environmental dataset for both current and future scenarios comprised average measures of nitrate concentration (mmol m□³), pH, bottom water temperature (°C), bathymetric depth (m), dissolved oxygen concentration (mmol m□³), chlorophyll-*a* concentration (mg m□³), seawater density (kg m□³), salinity (PSU), and current direction (degrees).

### 2.4 Ecological niche modeling

Species distribution models were developed using MaxEnt via the kuenm R package (Cobos et al., 2019). Model calibration involved testing multiple combinations of feature classes and regularization multipliers to identify optimal model complexity (Cobos et al., 2019). Candidate models were evaluated using partial ROC tests, omission rates, and Akaike Information Criterion corrected for small sample sizes (AICc). For each species, 10,000 background points were randomly sampled from the species-specific accessible area (M), which was defined using a 200-km buffer around the available occurrence records (Simões et al., 2021). Restricting background sampling to this accessible area was intended to represent the environmental conditions potentially available to each species while avoiding sampling background environments that were unlikely to have been accessible to the species. The best-performing models were selected based on a multi-criteria evaluation approach, prioritizing low omission rates and minimum AICc values. Final models were generated using bootstrap replication to account for sampling uncertainty. Model outputs were produced in logistic format, representing continuous environmentally suitable area values ranging from 0 (unsuitable) to 1 (highly suitable) (Simões et al., 2021). Final models were projected onto current environmental conditions and future climate scenarios (RCP 2.6 and RCP 8.5 in 2050 and 2100). Continuous habitat suitability maps (logistic outputs) were transformed into binary presence–absence maps using a fixed suitability threshold of 0.5, with raster cells having suitability values ≥ 0.5 classified as suitable habitat and values < 0.5 classified as unsuitable habitat. This fixed threshold was used as a standardized criterion across species, scenarios, and projection periods to facilitate consistent estimation and comparison of suitable habitat area and centroid displacement. The threshold was used as an analytical classification criterion rather than as a species-specific ecological tolerance threshold. This fixed threshold was used as a standardized criterion across species, scenarios, and projection periods to facilitate consistent estimation and comparison of suitable habitat area and centroid displacement. The threshold was used as an analytical classification criterion rather than as a species-specific ecological tolerance threshold

For each species and scenario, replicated model outputs were summarized by calculating the median suitability across replicates, which served as the primary representation of predicted species distributions. The range of suitability values across replicates was calculated to assess model uncertainty and was examined in supplementary analyses.

For each species and scenario, the following metrics were calculated following Simões et al., (2021)

1. Suitable habitat area (km²) under current and future conditions
2. Net change in suitable area, expressed as both absolute (km²) and relative (%) differences
3. Stable, lost, and gained habitat, representing areas predicted to remain suitable, become unsuitable, or become newly suitable under future conditions
4. Geographic centroid shift, calculated as the great-circle distance (km) between the centroids of the current and future suitable habitats
5. Niche similarity, quantified using Schoener’s D to assess changes in suitability patterns between current and future projections

Table S2 revealed variable importance values extracted from the final MaxEnt model for species that show a statistically significant response. To ensure robust statistical inference, feeding groups represented by only a single species were excluded from the comparative analyses. This avoided confounding feeding-group effects with species-specific responses. Principal Component Analysis (PCA) was used to examine patterns in environmental variable importance among species based on permutation importance values obtained from the species distribution models.

### 2.5 Comparative analysis among feeding strategy groups

To evaluate differences in species responses among feeding groups under different climate scenarios and projection periods, we applied linear mixed-effects models (LMMs) using the lme4 (Bates et al., 2015) and lmerTest (Kuznetsova et al., 2017) R packages. LMMs are appropriate for hierarchical and repeated-measures data because they allow fixed effects to be evaluated while accounting for species-level variation through random effects.

Two response variables derived from the species distribution model projections were analyzed: 1) percentage change in suitable area (%) and 2) geographic centroid displacement (km). Similar measures of percentage change in climate-suitable areas have been widely used to quantify projected range contraction or expansion under climate change scenarios (Adde et al., 2024). Similarly, Centroid-based measures are commonly used to quantify the magnitude and direction of projected species range shifts in ecological niche modelling studies (Abbasi et al., 2024).

Feeding type and a combined scenario–period factor were included as fixed effects. The scenario–period factor represented the four projection combinations including RCP 2.6–2050, RCP 2.6–2100, RCP 8.5–2050, and RCP 8.5–2100. This specification was used because the effect of the projection period is expected to be interpreted in conjunction with the climate scenario rather than as an independent temporal effect. The model was therefore specified as:

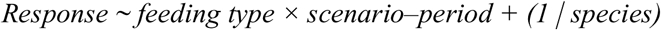

Species identity was included as a random intercept to account for repeated observations of the same species across different climate scenarios–period combinations and to control for species-specific differences in the magnitude of projected responses. The use of species-level random effects is consistent with the hierarchical structure of the data and the formulation of LMMs (Bates et al., 2015).

Significance of fixed effects was assessed using Type III analysis of variance with Satterthwaite’s approximation for denominator degrees of freedom, as implemented in lmerTest (Kuznetsova et al., 2017). When significant interactions were detected, pairwise comparisons among feeding groups within each scenario–period combination were performed using estimated marginal means with Tukey adjustment, implemented in the emmeans R package (Lenth & Piaskowski, 2026).

Schoener’s D was initially calculated to quantify the similarity between current and projected suitability distributions. Schoener’s D ranges from 0 (no overlap) to 1 (identical distributions) and has been widely used as a measure of niche or distributional similarity in ecological niche modelling (Rossi et al., 2026; Tong et al., 2023; Warren et al., 2008). All spatial calculations, including raster processing and calculation of geographic distances between current and future distribution centroids, were conducted in R using the raster and geosphere packages

## 3. Results

### 3.1 Model statistics

The present and future distribution model outputs for 17 amphipod species, representing different feeding groups, are summarized in Table 1. To balance model reliability with the retention of as many species as possible, we applied threshold criteria based on standard metrics for species distribution modeling. Specifically, species were flagged if their Partial ROC value was below 1 or their 5% omission rate exceeded 0.2. These criteria are commonly used to identify models with poor predictive performance, while still allowing the inclusion of species with limited occurrence records or broad geographic ranges, which are often challenging to model accurately at a global scale (Cobos et al., 2019; Peterson et al., 2008). Therefore, for further analyses, we removed *Corophium volutator* (Pallas, 1766) and *Cymadusa filosa* Savigny, 1816.

**Table 1.**
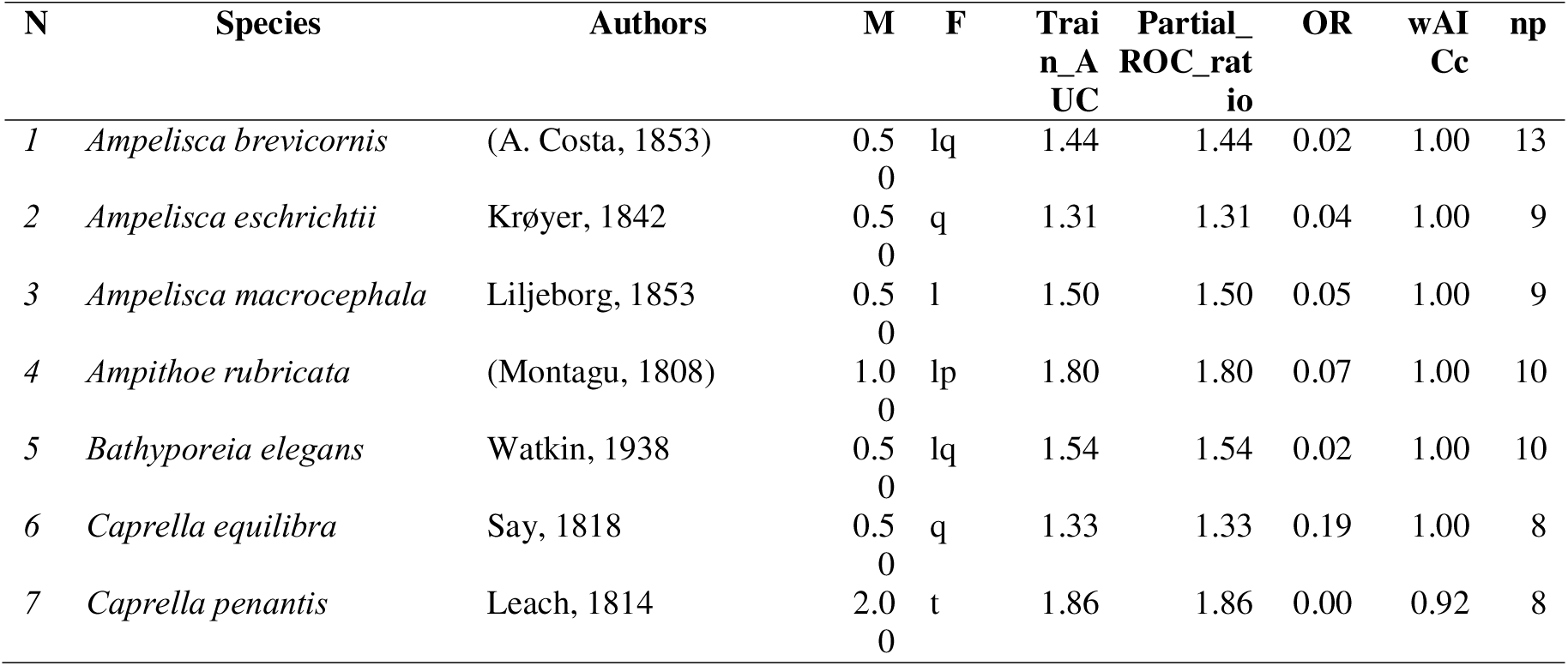
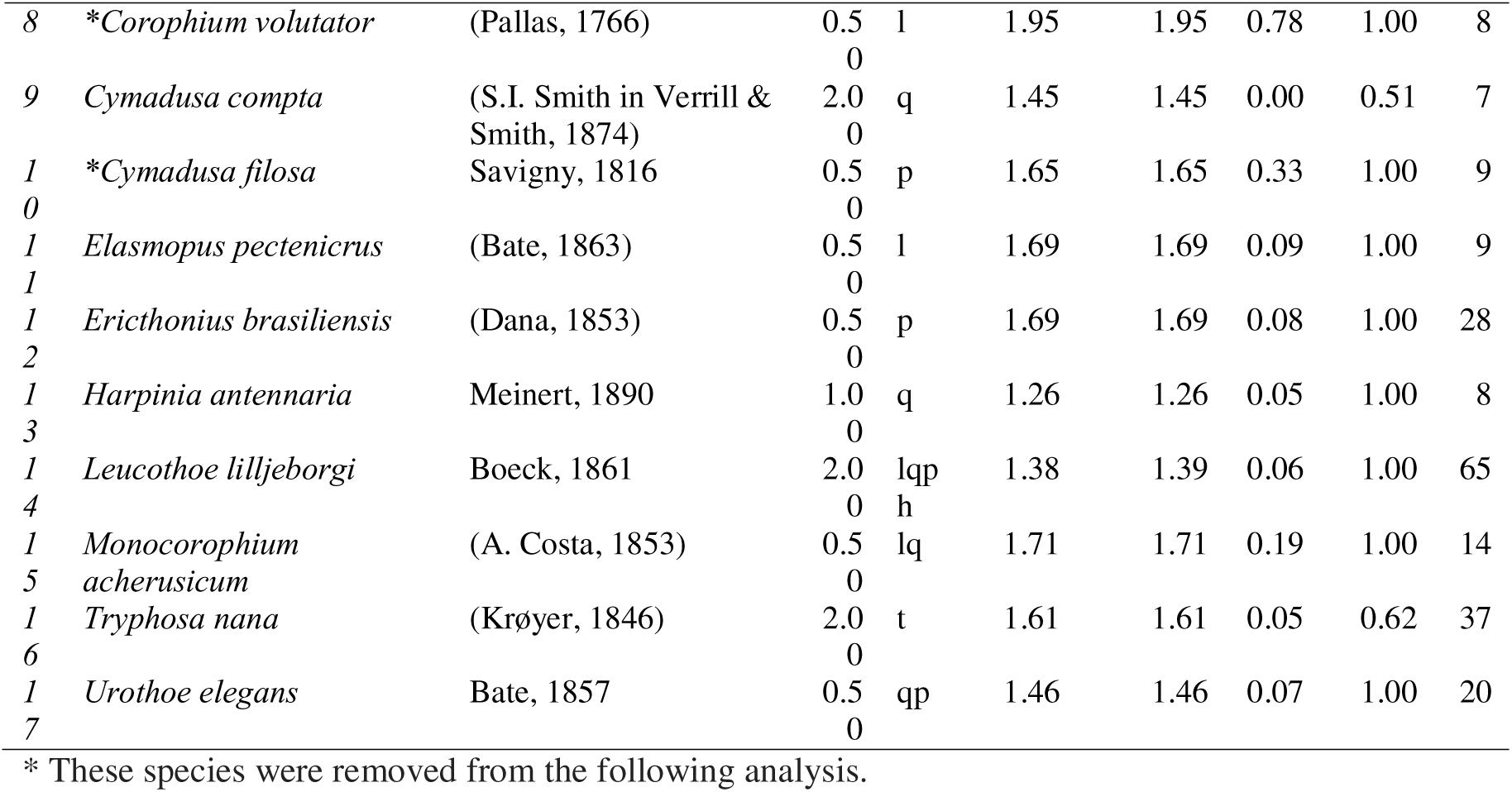
MaxEnt model evaluation for all 17 benthic amphipod species included in the study. M: regularisation strength, F: types of features used, Train AUC: Area under the Curve for training data, ROC: Receiver Operating Characteristic, OR: Omission rate, wAICc: Akaike weight for AICc, np: number of parameters in model. Species with an asterisk in front of their name were removed either when OR was >2 (i.e., >0.2), partial ROC was below 1.3, or training AUC was below 0.5.

Amphipod species showed distinct responses to climate-driven changes across different scenarios and time periods (Fig. 1). The magnitude of habitat changes increased under the stronger emissions scenario and longer projection period, but responses remained highly species-specific, with both substantial gains and losses occurring within the amphipod assemblage.

**Fig. 1.**
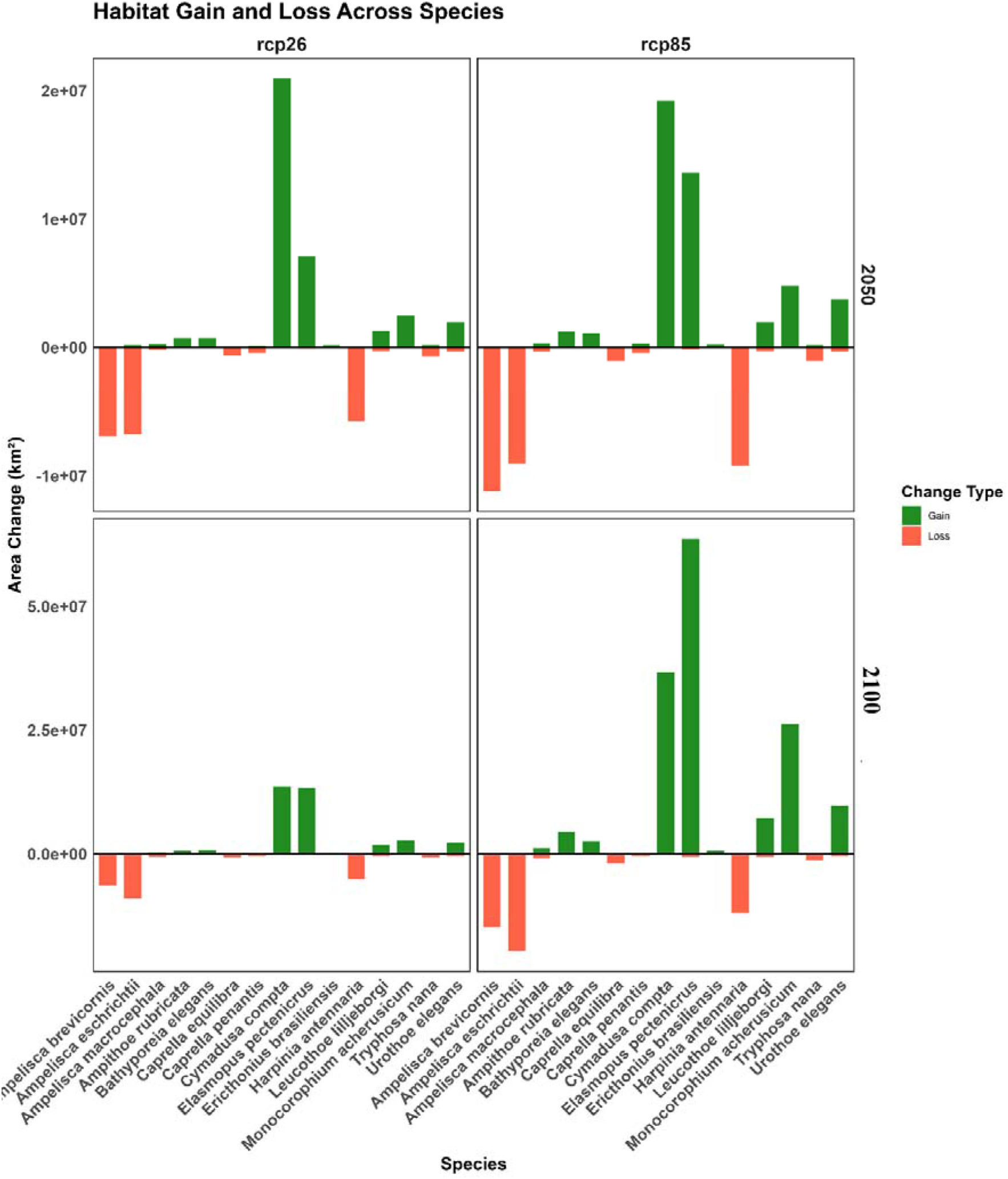
Suitable habitat area (km²) of 15 benthic amphipod species under current and future climate scenarios. Top panel: 2050 projections under RCP 2.6 and RCP 8.5. Bottom panel: 2100 projections under both scenarios. Bars represent the predicted area gained and lost for each species.

Under RCP 2.6, *Cymadusa compta* (Leach, 1814) and *Elasmopus pectenicrus* (Bate, 1862) showed the largest habitat gains, whereas *Ampelisca brevicornis* (Costa, 1853), *Harpinia antennaria* (Meinert, 1890), and *Ampelisca eschrichtii* Krøyer, 1842 experienced pronounced habitat losses. These differences became considerably stronger under RCP 8.5, particularly by 2100 *C. compta* and *E. pectenicrus* exhibited the largest projected gains, while *A. brevicornis* and *A. eschrichtii* showed the greatest suitable environmental losses (Figures S18-23).

### 3.2 Environmental Patterns of Climatic Suitability

The aggregate habitat suitability patterns revealed a predominantly temperate-to-high-latitude distribution under current climatic conditions (Fig. 2A), with higher suitability concentrated along the continental shelves of the North Atlantic and North Pacific. The overall geographic structure of suitable habitat remained broadly consistent across present and future scenarios, particularly under RCP 2.6 (Fig. 2B-C), except for a moderate loss of habitat suitability in comparison with the current status. Under RCP 8.5 (Fig. 2D-E), habitat suitability decreased by 2050 compared with the current conditions. However, by the end of the century, habitat suitability increased relative to the 2050 projection under the same scenario, although it remained lower than under current conditions. Figure 2 represents aggregate mean suitability, rather than the response of individual species. Therefore, areas with high values identified locations where, on average, the 15 modeled species have relatively high predicted suitability. The extracted habitat suitability for each species was presented under Figs 1-17S.

**Fig. 2.**
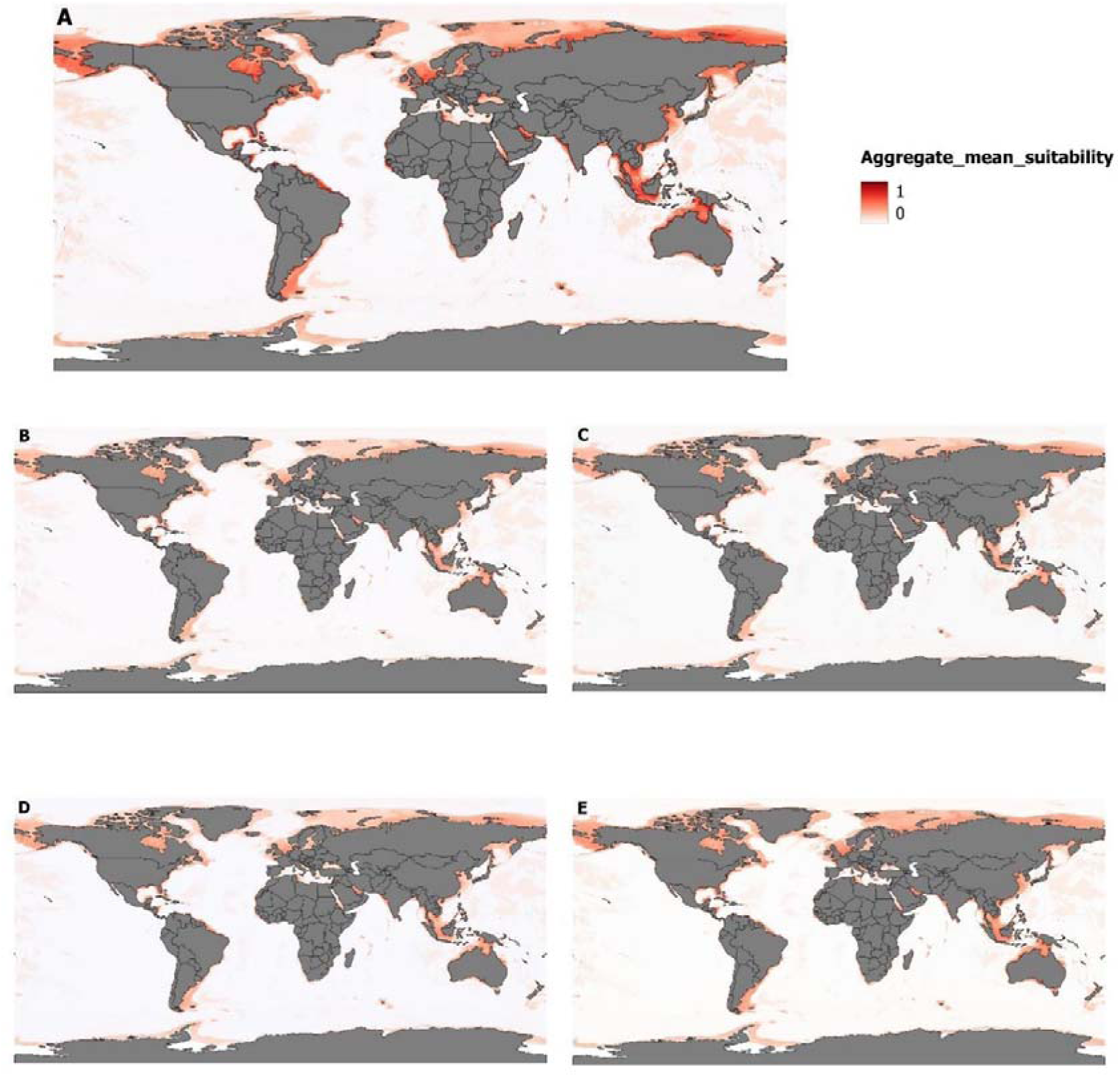
Global aggregate habitat suitability of 15 benthic amphipod species under present and future climate scenarios. Spatial projections represent the mean habitat suitability (0–1 scale) averaged across all modeled species. Warmer colors (red) indicate higher aggregate suitability, whereas cooler colors (white) indicate low suitability. Panels show: **(A)** Current conditions, **(B)** RCP 2.6 – 2050, **(C)** RCP 2.6 – 2100, **(D)** RCP 8.5 – 2050, and **(E)** RCP 8.5 – 2100.

Projected the number of studied species (species richness) of the studied species was strongly spatially distributed, with the highest richness concentrated along continental margins and shelf regions (Fig. 3). Under current conditions, areas of comparatively high species richness occurred mainly in the North Atlantic and North Pacific, northern European waters, and parts of the western Pacific and Southern Hemisphere. The broad spatial structure under RCP 2.6 (Fig. 3 B-C) will decrease in comparison with the current status; however, it will remain relatively consistent in time scale. Under RCP 8.5 (Fig. 3 D-E), changes in studied species richness will experience a stronger decrease until the mid of the century but at the end of the century in there will be an increase in species richness relative to the current status. Overall, no changes in pattern can be seen under different Scenarios and time scales.

**Fig. 3.**
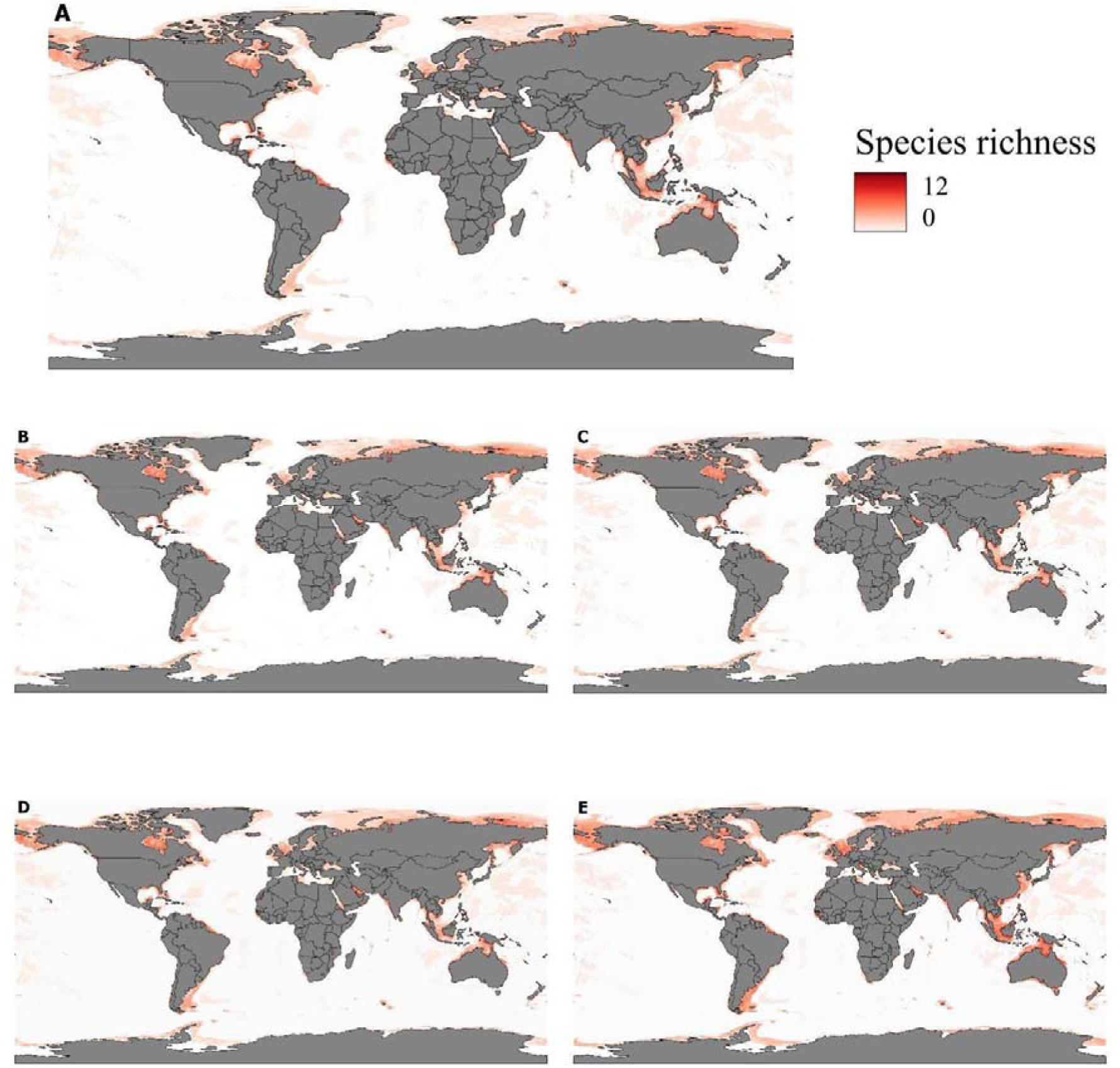
Global patterns of potential species richness for 15 benthic species under current and future climate scenarios. Studied species richness represents the number of species predicted as suitable per grid cell (range: 0–12). (A) Current conditions; (B) RCP 2.6 for 2050; (C) RCP 2.6 for 2100; (D) RCP 8.5 for 2050; and (E) RCP 8.5 for 2100. Warmer colors (orange–red) indicate higher species richness, while white indicates low richness. Future projections are based on Representative Concentration Pathways (RCPs) 2.6 and 8.5 for mid- and late-century periods.

Regarding the direction of potential distribution shifts, shallow-sea species were projected to exhibit northward shifts in their environmental suitability centroids (i.e., increases in centroid latitude) under all studied scenarios and time horizons. However, the centroid shifts never reached the Polar regions, even though the suitable habitat expansion does. The magnitude of these shifts was greater by 2100 than by 2050 and was more pronounced under RCP 8.5 compared to RCP 2.6. In contrast, deep-sea species showed projected southward shifts in their environmental suitability centroids across all scenarios (Fig. 4). *Elasmopus pectenicrus* will potentially experience the strongest shift under RCP 8.5 at the end of the century, and it will be replaced from the tropical region to temperate zones. The same habitat shifting with less strength can be seen for *Monocorophium acherusicum* and *Ampithoe rubricata*. The change in centroid shift of *Ampelisca brevicornis* is prominent in the temperate region world toward the polar zone.

**Fig. 4.**
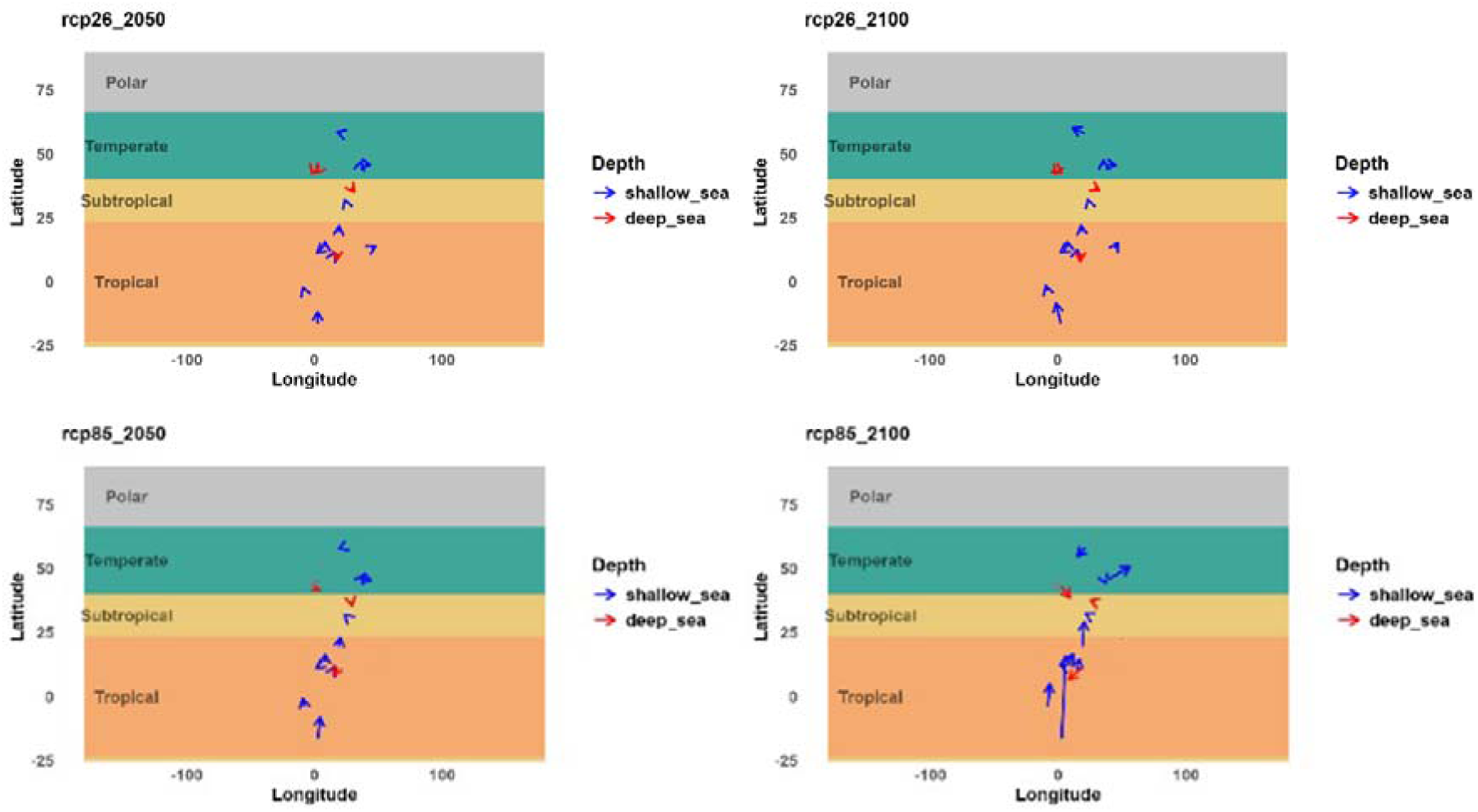
Maps show suitability centroid shifts between different climatic zones, projected for shallow-sea species for 2050 RCP 2.6 and RCP 8.5, 2100 RCP 2.6 and RCP 8.5, deep-sea species marked with red arrow and shallow-sea species marked with blue arrow.

### 3.3 Impact of environmental variables

The results of PCA analysis have shown the importance values of the extracted variables from the final MaxEnt model for amphipod species grouped by feeding groups. The first two principal components explained 64.3% of the total variance (PC1 = 41.5%, PC2 = 22.8%) (Fig. 5). Arrows represent the contribution and direction of environmental variables. PC1 is the main environmental contrast, and the strongest gradient is along PC1. PC1 primarily reflects a gradient from nutrient- and oxygen-associated conditions (positive scores) to more saline environments (negative scores, particularly in combination with PC2). PC1 appears to represent a broad contrast between species whose environmental importance profiles are more associated with nutrients, oxygen, and temperature versus those associated more with depth, chlorophyll, and seawater velocity (SWV). However, PC2 separates the species vertically. The upper part of the plot is strongly associated with salinity, pH, and SWV, while the lower part is more strongly associated with depth and Chlorophyll. PERMANOVA also could not detect a significant effect of the feeding group on environmental variable importance (F = 1.38, R² = 0.23, p = 0.23). Pairwise comparisons were not significant after correction for multiple testing (Table 3S3). The pairwise PERMANOVA results show no significant intragroup variance (Table S4).

**Fig. 5.**
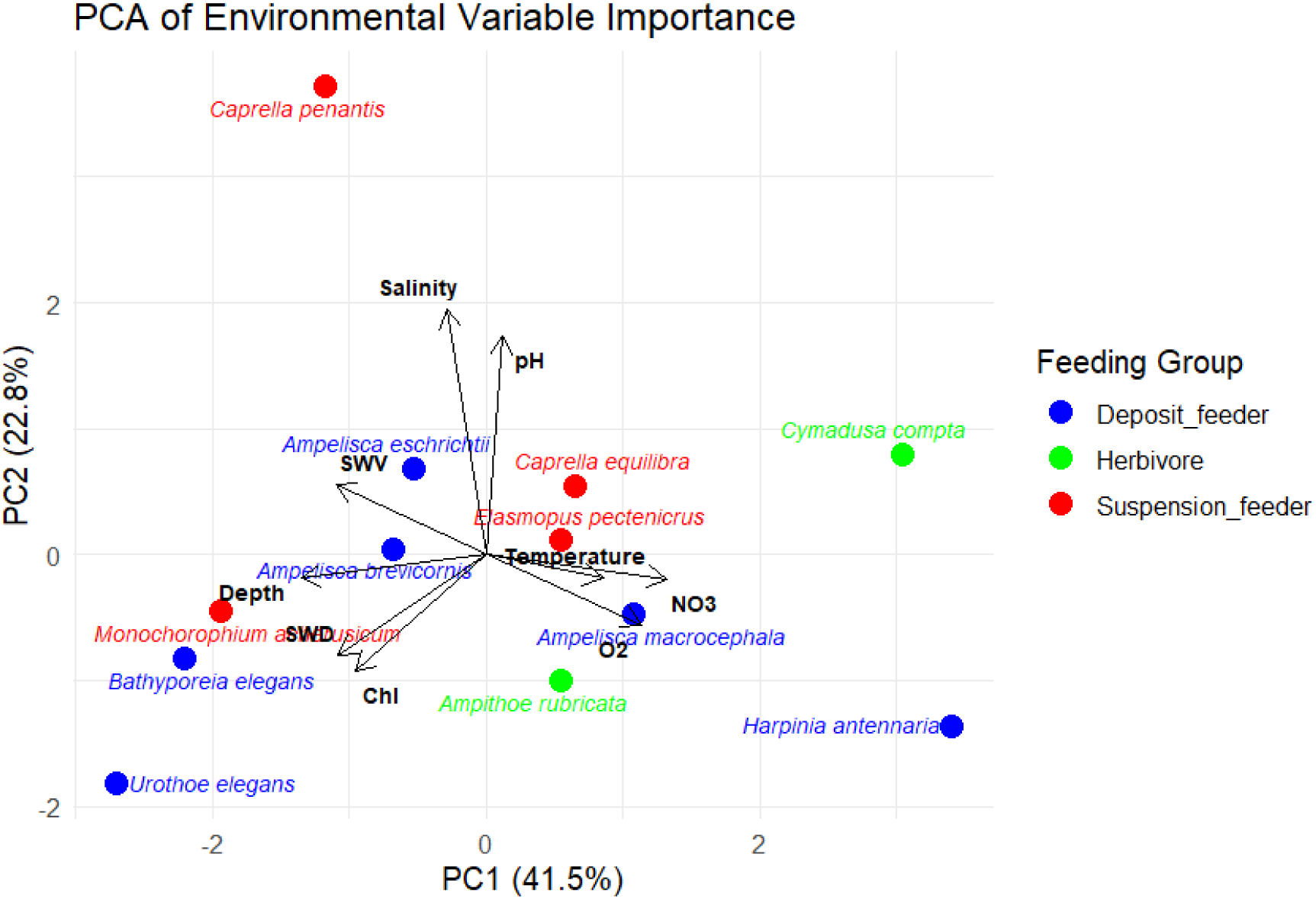
Principal Component Analysis (PCA) – biplot of variable importance values extracted from the final MaxEnt model for 15 amphipod species grouped based on feeding groups. The first two dimensions of PCA represent 64.3% of the variation. Arrows represent the contribution and direction of environmental variables. The abbreviations used for variables are SWV, Sea water velocity; SWD, Sea water direction; and Chl, chlorophyll-a concentration.

### 3.4 Impact of the feeding group

The percentage change in suitable habitat area did not differ significantly among feeding groups (LMM: FO,OO.O = 1.23, p = 0.322). Neither the interaction between feeding type and scenario–time (F□,□□.□ = 0.99, p = 0.445) nor the effect of scenario–time alone (F□,□□.□ = 2.30, p = 0.091) was statistically significant. Tukey-adjusted pairwise comparisons confirmed the absence of significant differences among feeding groups under any scenario–time combination (all p > 0.05; Table S5).

Centroid shift was significantly affected by scenario–time (F□,□□.□ = 5.51, p = 0.0029), and a significant interaction between feeding type and scenario–time was detected (F□,□□.□ = 4.55, p = 0.0013), indicating that the response of feeding groups depended on climate scenario and time period. In contrast, the main effect of feeding type on future distribution was not significant (F□,□□.□ = 0.85, p = 0.447). Tukey-adjusted post hoc comparisons revealed that an interaction exists due to a significant difference between deposit-feeders and suspension-feeders under the RCP8.5–2090 scenario (estimate = 1466 km, SE = 489 km, t = 3.00, p = 0.013). No other pairwise comparisons were statistically significant (Table S6). Niche overlap (Schoener’s D) was not significantly influenced by feeding type (F□,□□ = 0.56, p = 0.574), scenario–time (F□,□□ = 0.39, p = 0.762), or their interaction (F□,□□ = 0.57, p = 0.753). Consistent with these results, Tukey-adjusted pairwise comparisons detected no significant differences among feeding groups (all p > 0.05; Table S7)

## 4. Discussion

Considering amphipod’s low-dispersal biology (direct development), sensitivity to multiple climate-driven stressors (warming, deoxygenation, acidification), and central roles in benthic nutrient cycling and food webs, assessing amphipod responses to climate change is essential for forecasting benthic biodiversity change, maintaining ecosystem function, and designing effective conservation strategies (Habibi-Motlagh et al., 2026; Mandelli et al., 2025; Momtazi & Maghsoudlou, 2022; Shokri et al., 2024). In recent years, there has been an escalation of studies investigating how marine organisms respond to climate change scenarios using ecological niche models (ENMs) (Melo-Merino et al., 2020; Chaudhary et al., 2023; Kürzel et al., 2025; Kaiser et al., 2026). Although the theoretical and empirical limitations of ecological niche modeling (ENM) must be acknowledged when interpreting projections—such as niche truncation, assumption of niche conservatism, uncertainty from climate models and scenarios, and omission of biotic interactions and dispersal constraint (Kaiser et al., 2026; Sánchez-Fernández et al., 2016; Shin et al., 2025)s—ENM remains one of the most powerful and widely used tools for investigating species responses to climate change (Cuervo et al., 2023; Peterson et al., 2018; Wiens et al., 2009). Amphipods have rarely been examined in climate change impact studies. As an example, they were partially included in the global assessment by Simões et al. (2021) alongside other crustaceans, and have been studied at regional scales in the North Pacific (Kaiser et al., 2026) and the North Atlantic (Kürzel et al., 2025). The current study provides a broader comparison of climate-driven distributional responses across multiple amphipod species and feeding groups at a global scale. The present study attempts to reveal amphipod responses to future climate scenarios. It examined how trophic identity, considering bathymetric distribution, shapes species-specific vulnerability and redistribution patterns of amphipods globally. Our results suggest that amphipod responses were highly species-specific and changed across scenarios and time scales. However, centroid displacement revealed a different dimension of response, indicating that similar changes in habitat extent can be accompanied by substantial geographic redistribution.

The results, in agreement with previous studies on benthic species (Kürzel et al., 2025; Moraitis et al., 2019; Simões et al., 2021) indicate that the integrated habitat suitability of amphipods (Fig. 2), irrespective of depth distribution or feeding group, is projected to undergo spatial redistribution rather than simple contraction or expansion. These changes were characterized by substantial interspecific heterogeneity, with the magnitude of projected habitat redistribution generally increasing under the high-emission scenario and toward the end of the century. Although several species showed relatively limited changes in suitable habitat, a subset exhibited pronounced gains or losses, indicating that climate change is unlikely to affect the modeled amphipod assemblage uniformly. Notable increases in suitable habitat were projected for *A. rubricata*, *C. compta, M. acherusicum, U.elegans, L. lilljeborgi*, and *E. pectenicrus*. Such increases may partly reflect the expansion of warm-affinity species into higher-latitude environments as ocean temperatures rise, a process commonly referred to as marine tropicalization (Hastings et al., 2020; Poloczanska et al., 2016). However, it should be avoided to simplify the results, due to the role of each specieś characteristics, such as dispersal ability, biotic interactions, and local environmental conditions, which are species-specific (Åkesson et al., 2021; Comte et al., 2014; Román-Palacios & Wiens, 2020; Sasaki et al., 2026). As an example, based on the AMBI framework (Borja et al., 2000) *M. acherusicum* represents a more opportunistic group that has the ability to disperse more easily. In contrast, substantial habitat loss is projected for *A. brevicornis*, *T. nana*, *C.equilibra*, *H. antennaria*, and *A. eschrichtii*. The contrasting responses among these species highlight the absence of a uniform climate-driven trajectory across the amphipod assemblage.

The overall habitat suitability will decrease under the low-emission scenario, with a constant pattern over time. In contrast, under the high emission scenario (RCP 8.5) a non-monotonic response in species distribution model (SDM) projections could be detected by an initial decline in habitat suitability by mid-century (2050) relative to the present, followed by a partial recovery by 2100 relative to 2050, but still below current suitability. This “dip then partial recovery” is explicitly reported for several marine groups. For commercial squids, mean global habitat suitability declines from present to 2050 and 2100 under all RCPs, but in 2100 it increases relative to 2050 for most scenarios while staying below present values; RCP 8.5 shows particularly strong latitudinal restructuring with losses in the tropics and gains at higher latitudes (Guerreiro et al., 2023). Similar dynamics are described for the Octopus vulgaris species complex, where mean suitability rises between 2050 and 2100 yet the overall trend versus present remains negative under RCP 6.0 and 8.5 (Borges et al., 2022). This could be explained by midÓcentury habitat loss driven by local conditions exceeding thermal and biogeochemical tolerances (Gallagher & Albano, 2023), while lateÓcentury gains under severe scenarios reflect newly suitable highÓ latitude areas as isotherms and productivity regimes shift poleward (Mendoza-Segura et al., 2023; Simões et al., 2021). However, it should be mentioned in the real world, an increase in habitat suitability in new areas does not automatically offset local extinctions caused by habitat loss elsewhere, especially for benthic amphipods due to limited dispersal (Ingels et al., 2012), ignoring persisting barriers for free dispersal in modeling process (De Kort et al., 2020), and time lags and demographic process (Zurell et al., 2023).

At the assemblage level, species richness of the studied species will experience an overall decrease with different strength levels under scenarios and time scales. Under RCP 2.6 the pattern will decrease slightly compared with the current status but remain relatively consistent through time. It is reflected that the broad latitudinal and shelf–offshore structure remains similar to the present, with modest poleward shifts and limited expansion of richness pattern change (Jones & Cheung, 2015). However, the response under RCP 8.5 is more complicated and will start with decreasing species richness in the mid-century and increase at the end of the century, even in the polar zone, higher than current status. Similar dynamics are reported for mariculture species richness potential, which declines in the tropics but increases markedly at high latitudes under RCP 8.5 (Oyinlola et al., 2020). The same considerations about increasing species richness after decreasing should be aware specially for amphipod species with limited dispersal ability.

From the other aspect, however, species richness values change across scenarios and time steps, but the broad spatial pattern of richness remains largely constant. This suggests that climate change may alter the magnitude of local species richness without fundamentally changing the overall geographic configuration of richness patterns. Nevertheless, the redistribution of individual species, reflected in increasingly pronounced species-specific habitat gains and losses under stronger climate forcing, indicates substantial turnover beneath this apparently stable large-scale richness pattern (Kristiansen et al., 2025; Wiig et al., 2008). These projected declines may reflect the increasing exposure of marine species to environmental conditions outside their realized climatic niches, particularly where ocean warming exceeds their thermal limits and reduces the availability of suitable habitat. Temperature is a major determinant of marine species distributions, and continued warming can contribute to range contractions when conditions exceed species-specific thermal tolerances (IPCC, 2022). Similar species-specific losses of habitat suitability and distributional shifts have recently been reported for benthic amphipods under future climate scenarios in the North Atlantic (Kürzel et al., 2025). Although our SDMs did not directly estimate physiological thermal limits, this pattern is consistent with a temperature-driven contraction of suitable climatic space. In contrast, species such as *Cymadusa compta*, which have predominantly tropical distributions but currently occur at higher latitudes, may benefit from warming and the expansion of suitable environments toward higher latitudes. Similar poleward expansions have been reported for other peracarid crustaceans (Kaiser et al., 2026; Kürzel et al., 2025). In other aspects, species projected to experience habitat loss are predominantly suspension feeders inhabiting shallow marine environments. The decline in suitability for these functional groups may be linked to reductions in organic matter supply and altered trophic fluxes in coastal and shallow systems under climate change (Hyndes et al., 2022; Laroche et al., 2022).

The invasion of new habitats is one of the impacts of climate change (Stachowicz et al., 2002), and the predicted expansion of *C. compta* suggests the possibility of its invasion into higher latitudes. This species, which currently has a regional distribution in the Gulf of Mexico and adjacent areas, is herbivorous and potentially highly dispersing via rafting similar to other herbivorous amphipods (Poore, 2005), and could invade temperate or subpolar ecosystems. The western Atlantic Ocean coast and the eastern Australian coast could be invaded by this species at the end of the century. The other species of this genus, *C. filosa,* is recognized as a cryptogenic or non-indigenous species in various marine environments, particularly the Mediterranean and Red Sea (Bueno et al., 2020; Zakhama-Sraieb et al., 2018)

The centroid shifted poleward for shallow-sea species, in contrast to a lower-latitude shift for deep-sea species. However, the poleward shift was reported for marine crustacean shallow-water species; an opposite direction in deep-sea crustacean species was reported by Simönes et al. (2019), and for some amphipod species by Künzel et al. (2025) in the North Atlantic. The main reasons mentioned for the results by Künzel et al. (2025) and Simönes et al. (2019) are the presence of refugia and distribution to deeper areas.

The distribution of species in PCA space showed strong overlap among feeding groups, and the wide dispersion of species within each group. Additionally, PERMANOVA results demonstrated that feeding strategy does not impose a consistent environmental-variable importance profile. Instead, environmental sensitivity appears to be strongly species-specific; these findings are consistent with findings for amphipod species in the North Sea (Lörz et al., 2022). In contrast, depth and nitrate appeared more closely associated with the overall separation of species in PCA space, although nitrate showed a relatively minor contribution. The association of herbivorous species with nutrient-related variables may reflect their dependence on primary producers and the environmental conditions controlling their availability, whereas the distribution of suspension feeders may be more closely related to water-column properties and hydrodynamic conditions

The LMM results, together with the projected distribution maps, suggest that climate change may reshape the geographic distribution of benthic amphipod species rather than simply reduce their total suitable habitat. The LMMs showed that the percentage change in suitable habitat did not differ significantly among feeding groups, climate scenarios–time combinations, or their interaction. This indicates that no feeding group consistently experienced greater habitat loss or gain than the others. However, the distribution maps showed that suitable areas can shift geographically even when the overall amount of suitable habitat changes by a similar magnitude among groups. A similar pattern of geographic redistribution has been reported for some amphipod species in the North Atlantic (Kürzel et al., 2025). In contrast to habitat-area change, centroid displacement showed a clear response to future climate conditions. The scenario–time effect was significant, and the significant interaction between feeding type and scenario–time indicates that the magnitude of geographic displacement differed among feeding groups under different projected conditions.

This suggests that feeding strategy may be more important for determining where suitable habitats shift than for determining how much suitable habitat is lost or gained. The greater centroid displacement under the high-emission scenario, particularly toward the late-century period, is consistent with the expected increase in climate velocity and the need for species to track suitable temperature and oxygen conditions across geographic space (Bridge et al., 2014; Hoppit & Schmidt, 2022; Shimeta et al., 2004; Weinert et al., 2022).

Deposit-feeding amphipods showed the largest centroid displacement under the RCP8.5–2090 projection. Post-hoc comparisons indicated that their displacement was significantly greater than that of suspension feeders under this scenario–time combination. One possible explanation is that deposit-feeding amphipods may be indirectly affected by changes in food availability at the seafloor. Many benthic amphipods depend directly or indirectly on particulate organic matter (POM) originating from surface-water productivity. Changes in primary production, ocean stratification, and the export of organic matter to the seafloor could therefore influence the distribution of detritus-dependent species. However, this mechanism should be considered as a possible explanation rather than a direct causal relationship, because our models did not explicitly include food-web interactions or POM flux.

Similar geographic redistribution has been reported for mobile scavenging amphipods in the North Atlantic, where some lysianassid and eophilontid species showed pronounced poleward and coastal movements, potentially associated with their broad thermal tolerances and changes in productivity (Kürzel et al., 2025). Together, these findings suggest that climate change may alter the spatial organization of benthic amphipod communities even when the overall amount of suitable habitat does not differ significantly among feeding groups. This highlights the importance of considering both habitat extent and geographic displacement when assessing climate-driven changes in marine species distributions.

From a conservation perspective, these findings suggest that management strategies should consider the functional composition of benthic communities, as climate-driven shifts in species distributions may reorganize trophic structure and ecosystem functioning even when the overall extent of suitable habitat remains relatively stable. Although no significant differences in habitat change were detected among feeding groups, projected geographic redistribution of species may alter species co-occurrence patterns, community composition, and ecological interactions, with potential consequences for ecosystem functioning in marine environments.

### Considerations and limitations

A careful methodological approach should be considered when interpreting the results of this study. Although the selected species represent a broad range of feeding strategies and bathymetric distributions, they constitute only a small fraction of the amphipod diversity. Therefore, the observed patterns should be interpreted as representative of these widespread species rather than the entire amphipod fauna. Additionally, although we used permutation to account for sample size, feeding groups were still represented by different numbers of species, which may have reduced statistical power to detect group-level differences, particularly for herbivorous amphipods. On the other hand, feeding categories represent broad ecological classifications, and considerable ecological variation may still exist among species within the same trophic group.

Other common considerations for each modeling study are also considerable in this study, such as ignoring biotic interactions, threshold dependence, and uncertainty of future climate projections. In addition, despite compiling occurrence records from multiple global databases and applying spatial thinning to reduce spatial autocorrelation, sampling effort remains uneven across geographic regions, with greater representation from Europe and North America than from many tropical and Southern Hemisphere regions. This limitation should be considered when interpreting projected habitat suitability patterns.

## 5. Conclusion

Overall, our findings provide predictions about amphipod response to climate change in both shallow and deep waters, and at the species and ecological group levels. Climate change is expected to reshape benthic amphipod distributions primarily through geographic redistribution, with some species showing potential range contractions or expansions under future scenarios. Ecologically, such redistribution may have substantial consequences despite a stable total habitat area. Geographic displacement can modify species co-occurrence patterns, alter trophic interactions, and restructure functional assemblages, particularly across latitudinal gradients. Given the limited dispersal capacity of many benthic amphipods relative to projected climate velocities, future distributions may reflect transient disequilibrium states. These results underscore the importance of incorporating functional traits and climate velocity into global assessments of benthic biodiversity responses to warming oceans. Within the planetary boundaries framework, these changes highlight the close connection between the climate change boundary and biosphere integrity (Richardson et al., 2023). Continued climate change may therefore influence marine biodiversity not only through habitat loss, but also through the redistribution and reorganization of species and their ecological functions

Changes in geographic position may precede measurable changes in habitat extent, suggesting that distributional shifts can occur even when the total area of climatically suitable habitat remains relatively stable. These changes in areas of high projected habitat suitability overlap, shifting, contradicting, or expanding under climate change were mentioned in other studies and emphasized the importance of considering redistribution hotspots in MPA designation and conservation management strategies (Brodie et al., 2025; Chapman et al., 2019; García Molinos et al., 2016). From a conservation and management perspective, spatial redistribution without net habitat loss does not imply ecological stability. Shifts in functional composition across regions may alter benthic ecosystem processes such as organic matter recycling, sediment bioturbation, and energy transfer within food webs. Therefore, static conservation frameworks and geographically fixed marine protected areas (MPAs) may be insufficient to safeguard future benthic biodiversity. Adaptive management strategies that anticipate poleward shifts, incorporate functional vulnerability, and prioritize emerging high-latitude habitats will be essential. Integrating trait-based sensitivity and projected climate velocities into marine spatial planning can enhance resilience-oriented conservation efforts under accelerating ocean warming.

## Acknowledgement

We would like to especially thank the Humboldt Foundation, George Foster Fellowship for supporting the first author to stay and conduct research in Germany.

## Author contributions

FM: designed the study, extracted data, ran the analyses, created all the plots, and wrote the manuscript. HS: helped in developing the study design, supervised and contributed to the data analysis and manuscript writing.

## Data Availability Statement

Species occurrence records analyzed in this study are publicly available from the Ocean Biodiversity Information System (OBIS) and the Global Biodiversity Information Facility (GBIF). Environmental variables are publicly available through the Bio-ORACLE database. The R scripts used for data processing, species distribution modeling, and analyses have been deposited in Zenodo at (https://doi.org/10.5281/zenodo.19471064).

