## supplementary file for "Climate-driven shifts in distributions and habitat suitability of amphipod species under future ocean change"

**Table S1.** Species included in the species distribution modeling workflow, showing the original number of occurrence records, the number of records retained after spatial thinning, feeding type, and modeling status. Species with fewer than 30 occurrence records after spatial thinning were excluded before model fitting. For species meeting the minimum occurrence threshold, models were fitted and evaluated where possible. “Successful” indicates that the modeling workflow was completed and the resulting models met the predefined evaluation criteria. “Incomplete” indicates that the modeling workflow could not be fully completed and therefore no final model results were obtained. “Excluded after modeling” indicates that models were fitted, but the resulting predictions did not meet the predefined criteria for statistical validity and were therefore excluded from subsequent analyses. Feeding types were assigned based on the predominant feeding strategy reported for each species.

|  | **Species** | **Original records** | **Records after thinning** | **Feeding type** | **Modeling status** | **Reason/Outcome** |
| --- | --- | --- | --- | --- | --- | --- |
| 1 | *Ampelisca brevicornis* | 7686 | 255 | Deposit feeder | Successful | Model successfully fitted and evaluated |
| 2 | *Harpinia antennaria* | 1108 | 420 | Deposit feeder | Successful | Model successfully fitted and evaluated |
| 3 | *Perioculodes longimanus* | 8489 | 371 | Deposit feeder | Incomplete | Modeling could not be completed |
| 4 | *Bathyporeia elegans* | 8428 | 343 | Deposit feeder | Successful | Model successfully fitted and evaluated |
| 5 | *Urothoe elegans* | 8,667 | 338 | Deposit feeder | Successful | Model successfully fitted and evaluated |
| 6 | *Ericthonius brasiliensis* | 2,229 | 325 | Suspension feeder | Successful | Model successfully fitted and evaluated |
| 7 | *Hippomedon denticulatus* | 8262 | 317 | Scavenger | Incomplete | Modeling could not be completed |
| 8 | *Tryphosa nana* | 4,47 | 273 | Scavenger | Successful | Model successfully fitted and evaluated |
| 9 | *Leucothoe lilljeborgi* | 5491 | 272 | Commensal | Successful | Model successfully fitted and evaluated |
| 10 | *Photis longicaudata* | 1,023 | 266 | Deposit feeder | Incomplete | Modeling could not be completed |
| 11 | *Ampelisca macrocephala* | 6402 | 264 | Deposit feeder | Successful | Model successfully fitted and evaluated |
| 12 | *Monochorophium acherusicum* | 2,258 | 196 | Suspension feeder | Successful | Model successfully fitted and evaluated |
| 13 | *Corophium volutator* | 12135 | 196 | Suspension feeder | Invalid | Model fitted, but results did not meet validity criteria |
| 14 | *Cymadusa compta* | 647 | 107 | Herbivore | Successful | Model successfully fitted and evaluated |
| 15 | *Ampelisca eschrichtii* | 666 | 92 | Deposit feeder | Successful | Model successfully fitted and evaluated |
| 16 | *Ampithoe rubricata* | 917 | 79 | Herbivore | Successful | Model successfully fitted and evaluated |
| 17 | *Platorchestia platensis* | 735 | 69 | Deposit feeder | Incomplete | Modeling could not be completed |
| 18 | *Elasmopus pectenicrus* | 662 | 61 | Suspension feeder | Successful | Model successfully fitted and evaluated |
| 19 | *Caprella penantis* | 1,254 | 56 | Suspension feeder | Successful | Model successfully fitted and evaluated |
| 20 | *Cymadusa filosa* | 699 | 47 | Herbivore | Invalid | Model fitted, but results did not meet validity criteria |
| 21 | *Caprella equilibra* | 899 | 43 | Suspension feeder | Successful | Model successfully fitted and evaluated |
| 22 | *Gammarus oceanicus* | 3441 | 30 | Herbivore | Excluded | Excluded at the initial filtering stage |
| 23 | *Westwoodilla caecula* | 6532 | 30 | Deposit feeder | Excluded | Excluded at the initial filtering stage |
| 24 | *Tmetonyx cicada* | 3542 | 29 | Scavenger | Excluded | <30 records after thinning |
| 25 | *Ampelisca typica* | 3,819 | 28 | Deposit feeder | Excluded | <30 records after thinning |
| 26 | *Eriopisa elongata* | 6514 | 28 | Deposit feeder | Excluded | <30 records after thinning |
| 27 | *Ampelisca spinipes* | 7546 | 27 | Deposit feeder | Excluded | <30 records after thinning |
| 28 | *Phoxocephalus holbolli* | 3963 | 26 | Deposit feeder | Excluded | <30 records after thinning |
| 29 | *Stenothoe monoculoides* | 3632 | 25 | Scavenger | Excluded | <30 records after thinning |
| 30 | *Tryphosites longipes* | 1867 | 24 | Scavenger | Excluded | <30 records after thinning |
| 31 | *Monocorophium insidiosum* | 3478 | 24 | Deposit feeder | Excluded | <30 records after thinning |
| 32 | *Argissa hamatipes* | 5795 | 24 | Deposit feeder | Excluded | <30 records after thinning |
| 33 | *Ampelisca tenuicornis* | 9289 | 22 | Suspension feeder | Excluded | <30 records after thinning |
| 34 | *Abludomelita obtusata* | 4361 | 21 | Scavenger | Excluded | <30 records after thinning |
| 35 | *Crassicorophium crassicorne* | 4208 | 20 | Suspension feeder | Excluded | <30 records after thinning |

**Table S2.** Relative importance of environmental variables in species distribution models for benthic amphipods. Values represent the **percent contribution** and **permutation importance** of each predictor variable, including average measures of depth, nitrate (NO₃), dissolved oxygen (O₂), pH, chlorophyll-a (Chl), salinity (S), seawater direction (SWD), seawater velocity (SWS), and temperature (T). Percent contribution reflects the relative influence of each variable during model training, whereas permutation importance indicates the decrease in model performance when the variable is randomly permuted. Species are grouped by habitat (shallow vs. deep sea) and feeding type. Higher values indicate greater importance of the corresponding environmental variable in shaping species distributions.

| **species** | **depth** | **feeding_type** | **Depth** | **NO3** | **O2** | **pH** | **Chl** | **S** | **SWD** | **SWV** | **T** |
| --- | --- | --- | --- | --- | --- | --- | --- | --- | --- | --- | --- |
| *Ampelisca brevicornis* | shallow sea | Deposit_feeder | 74.3 | 7.1 | 3.6 | 3.1 | 2.4 | 7.0 | 0.5 | 1.1 | 1.0 |
| *Ampelisca eschrichtii* | deep sea | Deposit_feeder | 52.6 | 1.2 | 8.5 | 5.1 | 9.3 | 15.4 | 1.1 | 0.5 | 6.3 |
| *Ampelisca macrocephala* | deep sea | Deposit_feeder | 36.1 | 11.7 | 2.1 | 1.8 | 0.9 | 3.0 | 0.4 | 1.4 | 42.7 |
| *Bathyporeia elegans* | shallow sea | Deposit_feeder | 72.8 | 7.6 | 0.8 | 0.0 | 4.2 | 6.2 | 4.1 | 2.9 | 1.4 |
| *Harpinia antennaria* | deep sea | Deposit_feeder | 12.3 | 36.5 | 36 | 3.5 | 2.3 | 0.0 | 0.3 | 0.2 | 9.0 |
| *Urothoe elegans* | deep sea | Deposit_feeder | 55.4 | 4.9 | 2.1 | 0.9 | 27.7 | 2.3 | 3.4 | 3.3 | 0 |
| *Ampithoe rubricata* | shallow sea | Herbivore | 37.9 | 23.2 | 4.4 | 0.8 | 7.2 | 4.0 | 1.7 | 1.1 | 19.7 |
| *Cymadusa compta* | shallow sea | Herbivore | 0.0 | 39.7 | 12.6 | 12.2 | 1.3 | 8.5 | 0.8 | 0.1 | 24.8 |
| *Elasmopus pectenicrus* | shallow sea | Suspension_feeder | 35.6 | 6.4 | 6.7 | 6.1 | 2.9 | 8.6 | 1.9 | 0.6 | 31.2 |
| *Caprella equilibra* | shallow sea | Suspension_feeder | 31.9 | 22.6 | 4.0 | 7.2 | 1.4 | 10.7 | 1.9 | 0.0 | 0.0 |
| *Caprella penantis* | shallow sea | Suspension_feeder | 44.4 | 8.2 | 0.0 | 17.8 | 1.9 | 21.4 | 0.7 | 4.4 | 1.3 |
| *Monochorophium acherusicum* | shallow sea | Suspension_feeder | 55.3 | 4.3 | 2.4 | 11.3 | 16.7 | 4.4 | 4.1 | 1.5 | 0.0 |

**Table S3.** Pairwise PERMANOVA results comparing environmental variables among feeding types (PERMANOVA; 999 permutations, Euclidean distance)

|  | **Group1** | **Group2** | **F_value** | **R2** | **p_value** | **p_adjusted** |
| --- | --- | --- | --- | --- | --- | --- |
| 1 | Deposit_feeder | Herbivore | 1.89 | 0.23 | 0.14 | 0.44 |
| 2 | Deposit_feeder | Suspension_feeder | 0.51 | 0.06 | 0.73 | 1.00 |
| 3 | Herbivore | Suspension_feeder | 2.46 | 0.39 | 0.2 | 0.6 |

**Table S4:** Pairwise PERMDISP results comparing environmental variables among feeding types (999 permutations, Euclidean distance)

| **Comparison** | **Difference** | **Lower CI** | **Upper CI** | **Adjusted p-value** |
| --- | --- | --- | --- | --- |
| Herbivore – Deposit feeder | -7.06 | -41.47 | 27.36 | 0.83 |
| Suspension feeder – Deposit feeder | -9.30 | -36.50 | 17.91 | 0.62 |
| Suspension feeder – Herbivore | -2.23 | -38.74 | 34.27 | 0.98 |

**Table S5.** Tukey post hoc comparisons among feeding groups for changes in suitable habitat area (%) under different climate scenarios and time periods

| **Scenario–Time** | **Comparison** | **Estimate** | **SE** | **df** | **t-ratio** | **p-value** |
| --- | --- | --- | --- | --- | --- | --- |
| RCP2.6–2050 | Deposit-feeder vs Herbivore | -82.98 | 157 | 48.7 | -0.530 | 0.8572 |
|  | Deposit-feeder vs Suspension-feeder | -32.52 | 116 | 49.8 | -0.281 | 0.9575 |
|  | Herbivore vs Suspension-feeder | 50.45 | 161 | 47.4 | 0.313 | 0.9475 |
| RCP2.6–2090 | Deposit-feeder vs Herbivore | -61.75 | 155 | 47.4 | -0.400 | 0.9159 |
|  | Deposit-feeder vs Suspension-feeder | -58.60 | 113 | 47.4 | -0.519 | 0.8623 |
|  | Herbivore vs Suspension-feeder | 3.15 | 161 | 47.4 | 0.020 | 0.9998 |
| RCP8.5–2050 | Deposit-feeder vs Herbivore | -95.09 | 155 | 47.4 | -0.615 | 0.8124 |
|  | Deposit-feeder vs Suspension-feeder | -63.28 | 113 | 47.4 | -0.561 | 0.8415 |
|  | Herbivore vs Suspension-feeder | 31.81 | 161 | 47.4 | 0.197 | 0.9788 |
| RCP8.5–2090 | Deposit-feeder vs Herbivore | -203.69 | 155 | 47.4 | -1.318 | 0.3922 |
|  | Deposit-feeder vs Suspension-feeder | -265.19 | 113 | 47.4 | -2.350 | 0.0586 |
|  | Herbivore vs Suspension-feeder | -61.51 | 161 | 47.4 | -0.381 | 0.9231 |

**Table S6.** Tukey post hoc comparisons among feeding groups for centroid shift distance.

| **Scenario–Time** | **Comparison** | **Estimate** | **SE** | **df** | **t-ratio** | **p-value** |
| --- | --- | --- | --- | --- | --- | --- |
| RCP2.6–2050 | Deposit-feeder vs Herbivore | -1121 | 663 | 37.7 | -1.690 | 0.2221 |
|  | Deposit-feeder vs Suspension-feeder | -225 | 489 | 38.6 | -0.460 | 0.8904 |
|  | Herbivore vs Suspension-feeder | 896 | 685 | 36.6 | 1.309 | 0.3994 |
| RCP2.6–2090 | Deposit-feeder vs Herbivore | -661 | 656 | 36.6 | -1.007 | 0.5770 |
|  | Deposit-feeder vs Suspension-feeder | -371 | 479 | 36.6 | -0.775 | 0.7205 |
|  | Herbivore vs Suspension-feeder | 290 | 685 | 36.6 | 0.423 | 0.9064 |
| RCP8.5–2050 | Deposit-feeder vs Herbivore | -236 | 656 | 36.6 | -0.360 | 0.9310 |
|  | Deposit-feeder vs Suspension-feeder | 124 | 479 | 36.6 | 0.258 | 0.9639 |
|  | Herbivore vs Suspension-feeder | 360 | 685 | 36.6 | 0.526 | 0.8591 |
| RCP8.5–2090 | Deposit-feeder vs Herbivore | 465 | 663 | 37.7 | 0.701 | 0.7644 |
|  | ***Deposit-feeder vs Suspension-feeder*** | ***1466*** | ***489*** | ***38.6*** | ***3.000*** | ***0.0128**** |
|  | Herbivore vs Suspension-feeder | 1001 | 685 | 36.6 | 1.463 | 0.3204 |

* Significant difference

**Table S7.** Tukey post-hoc comparisons among feeding groups for niche overlap.

| **Scenario–Time** | **Comparison** | **Estimate** | **SE** | **df** | **t-ratio** | **p-value** |
| --- | --- | --- | --- | --- | --- | --- |
| RCP2.6–2050 | Deposit-feeder vs Herbivore | 0.083 | 14200 | 70.3 | 0.000 | 1.0000 |
|  | Deposit-feeder vs Suspension-feeder | 0.018 | 10500 | 70.3 | 0.000 | 1.0000 |
|  | Herbivore vs Suspension-feeder | -0.066 | 14500 | 70.3 | 0.000 | 1.0000 |
| RCP2.6–2090 | Deposit-feeder vs Herbivore | 0.048 | 13900 | 70.3 | 0.000 | 1.0000 |
|  | Deposit-feeder vs Suspension-feeder | 0.012 | 10200 | 70.3 | 0.000 | 1.0000 |
|  | Herbivore vs Suspension-feeder | -0.036 | 14500 | 70.3 | 0.000 | 1.0000 |
| RCP8.5–2050 | Deposit-feeder vs Herbivore | 0.064 | 13900 | 70.3 | 0.000 | 1.0000 |
|  | Deposit-feeder vs Suspension-feeder | 0.029 | 10200 | 70.3 | 0.000 | 1.0000 |
|  | Herbivore vs Suspension-feeder | -0.035 | 14500 | 70.3 | 0.000 | 1.0000 |
| RCP8.5–2090 | Deposit-feeder vs Herbivore | -17600 | 13900 | 70.3 | -1.262 | 0.4213 |
|  | Deposit-feeder vs Suspension-feeder | -17600 | 10200 | 70.3 | -1.728 | 0.2020 |
|  | Herbivore vs Suspension-feeder | -0.105 | 14500 | 70.3 | 0.000 | 1.0000 |

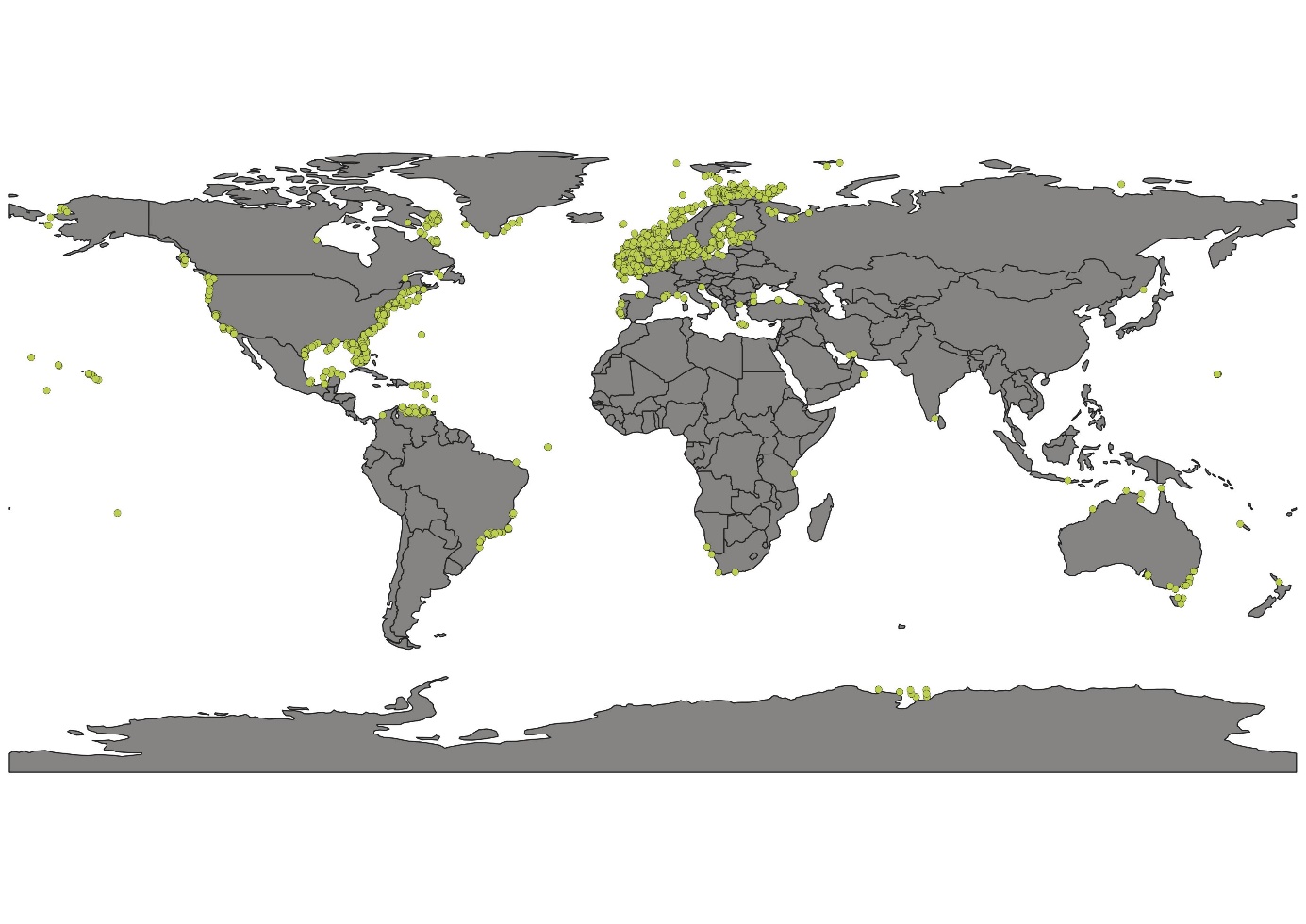

**Fig. 1S**. The geographical position of amphipod records (after cleaning the data) was applied for species distribution modeling, containing 3351 records.

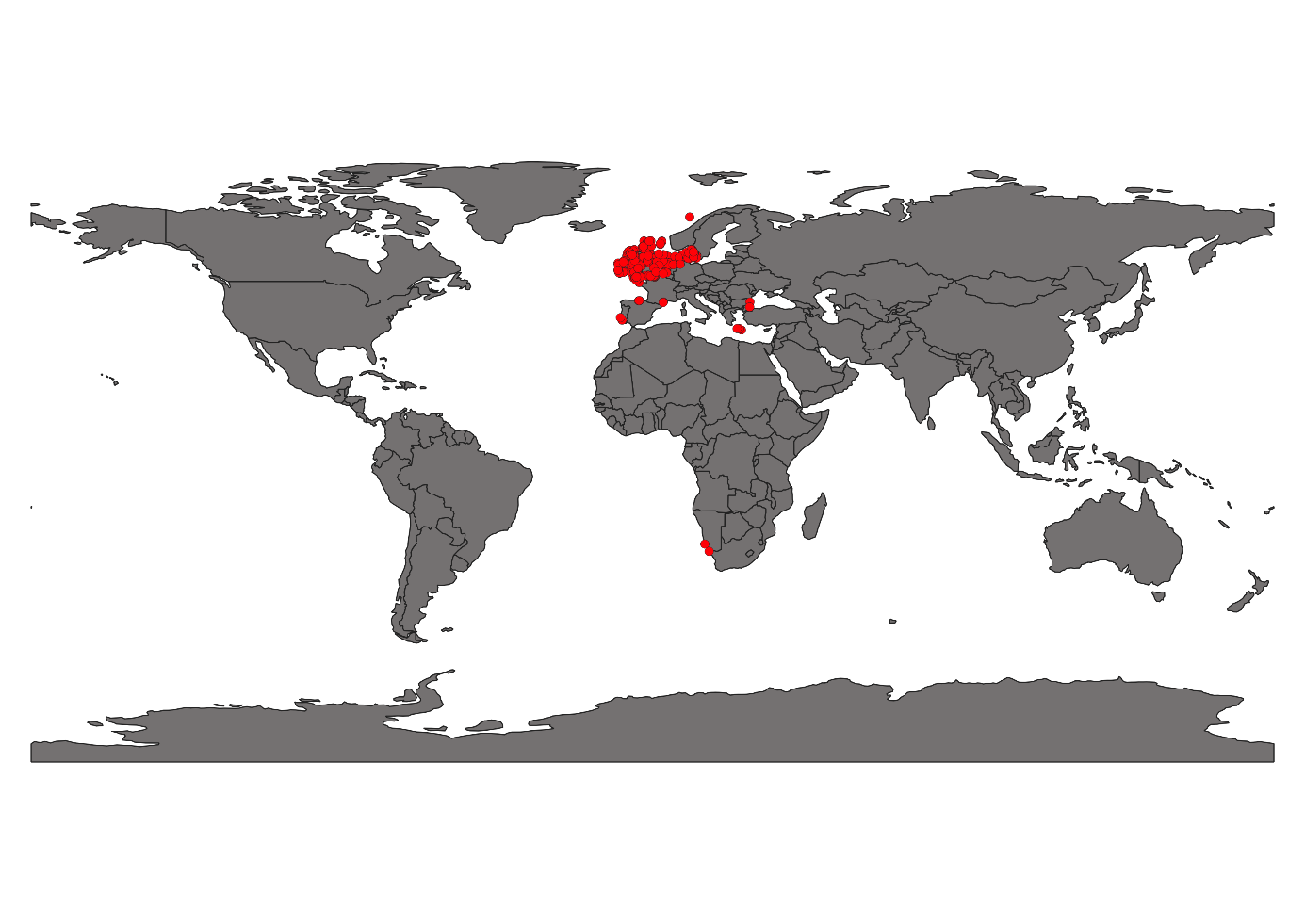

**Fig S2.** The geographical position of Ampelisca brevicornis (after thinning the data) was applied for species distribution modeling.
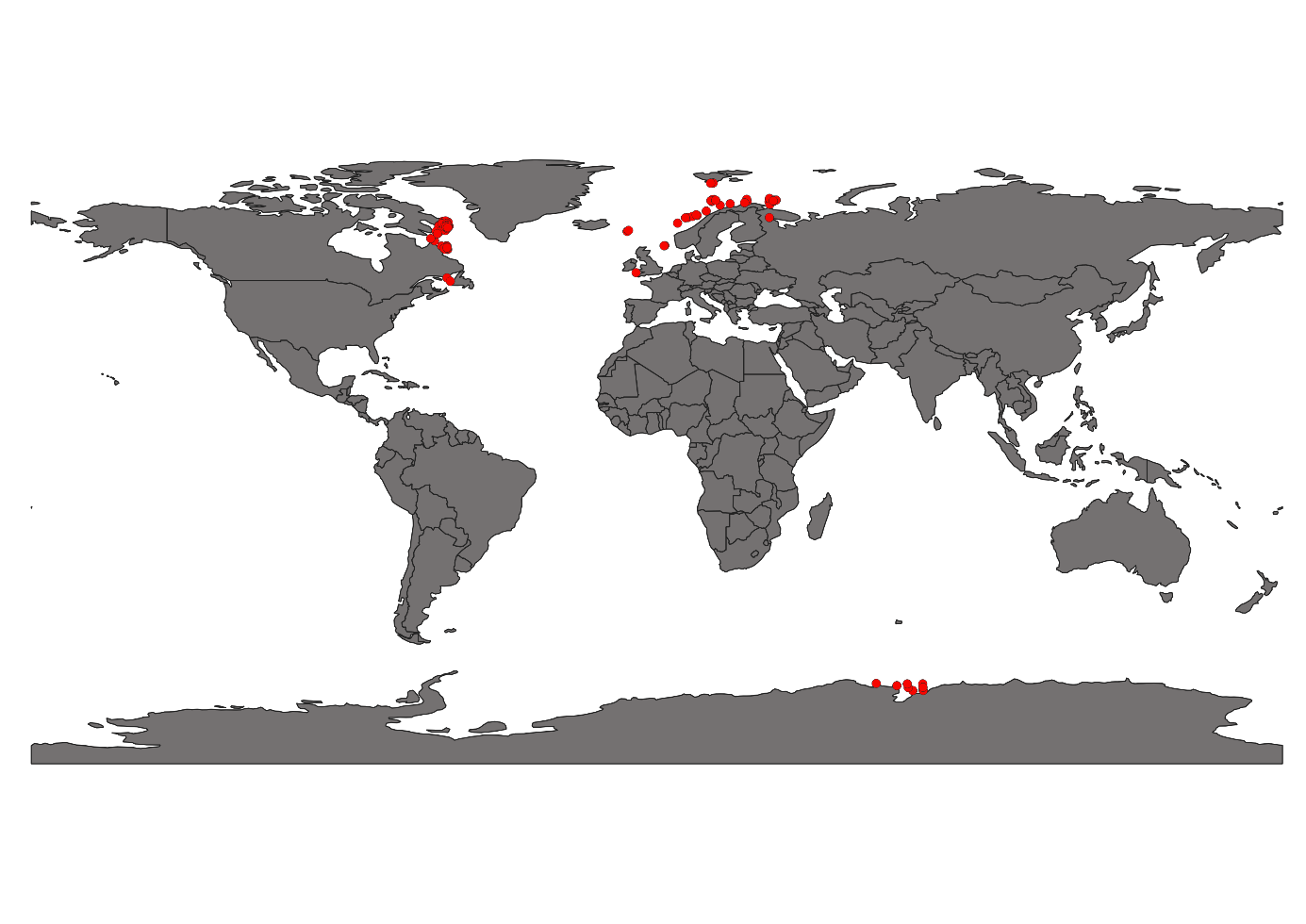

**Fig S3.** The geographical position of *Ampelisca eschrichtii* (after thinning the data) was applied for species distribution modeling.

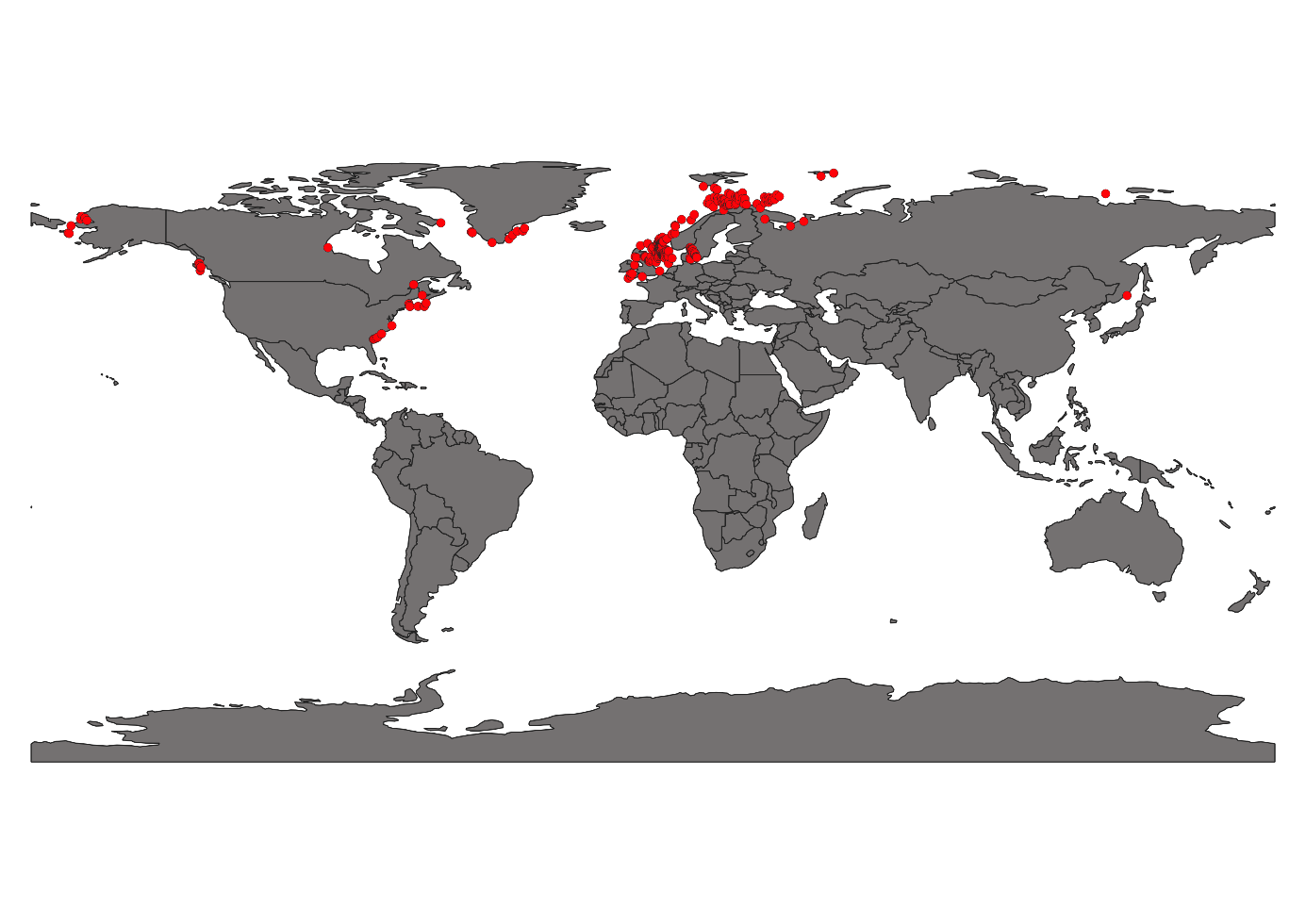

**Fig S3.** The geographical position of *Ampelisca macrocephala* (after thinning the data) was applied for species distribution modeling.

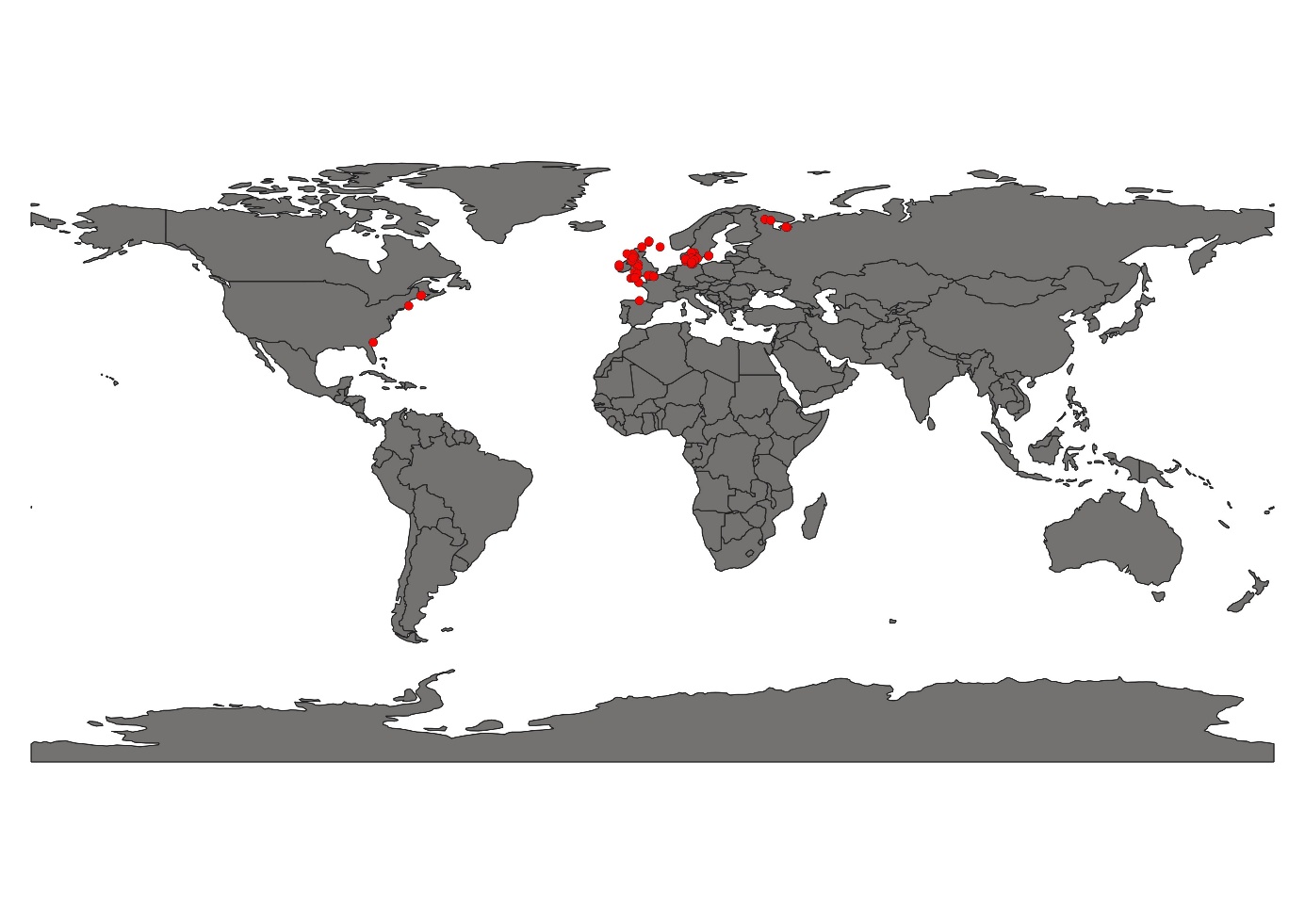

**Fig S4.** The geographical position of *Ampithoe rubricata* (after thinning the data) was applied for species distribution modeling.

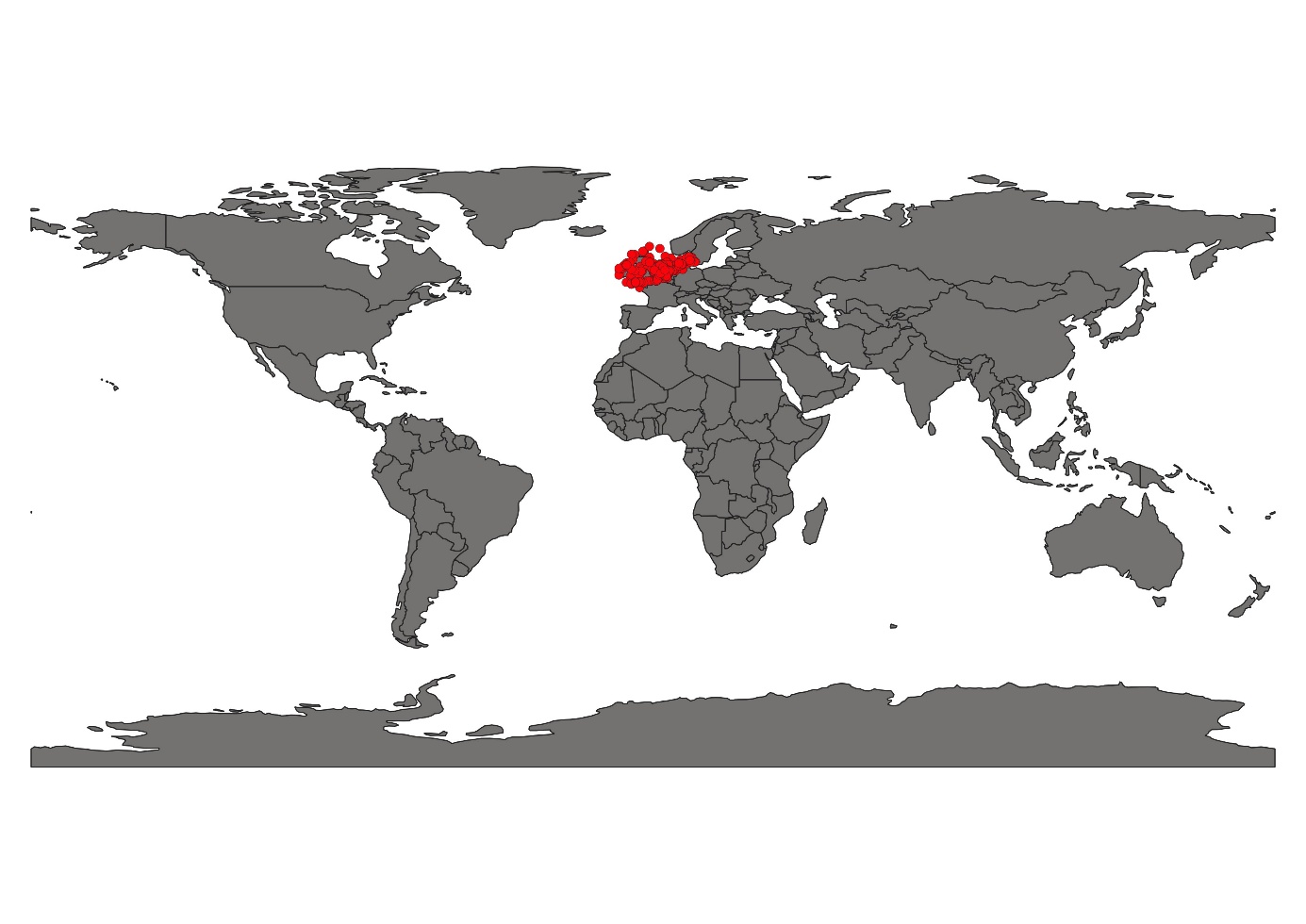

**Fig S5.** The geographical position of *Bathyporeia elegans*(after thinning the data) was applied for species distribution modeling.

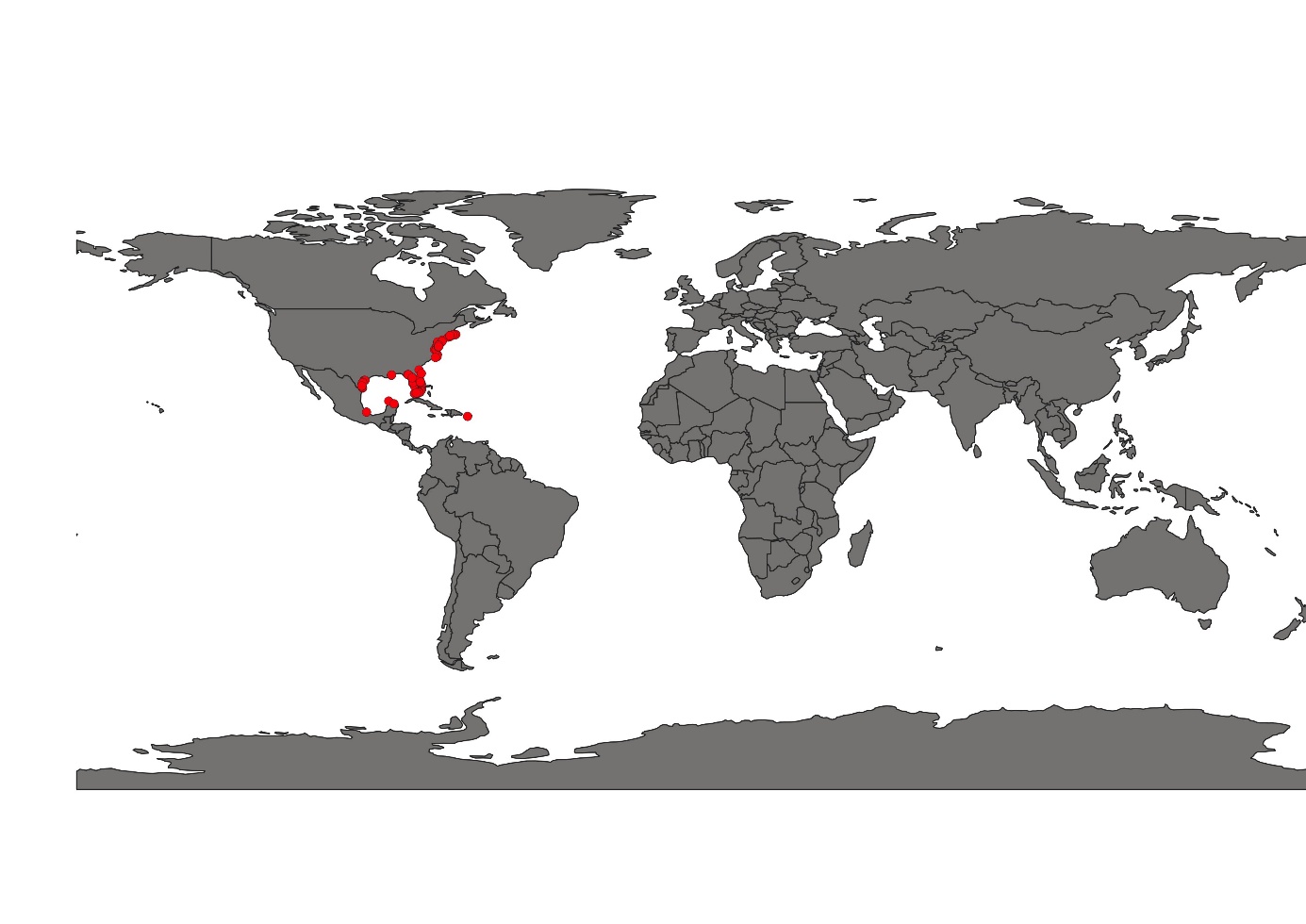

**Fig S6.** The geographical position of *Cymadusa compta* (after thinning the data) was applied for species distribution modeling.

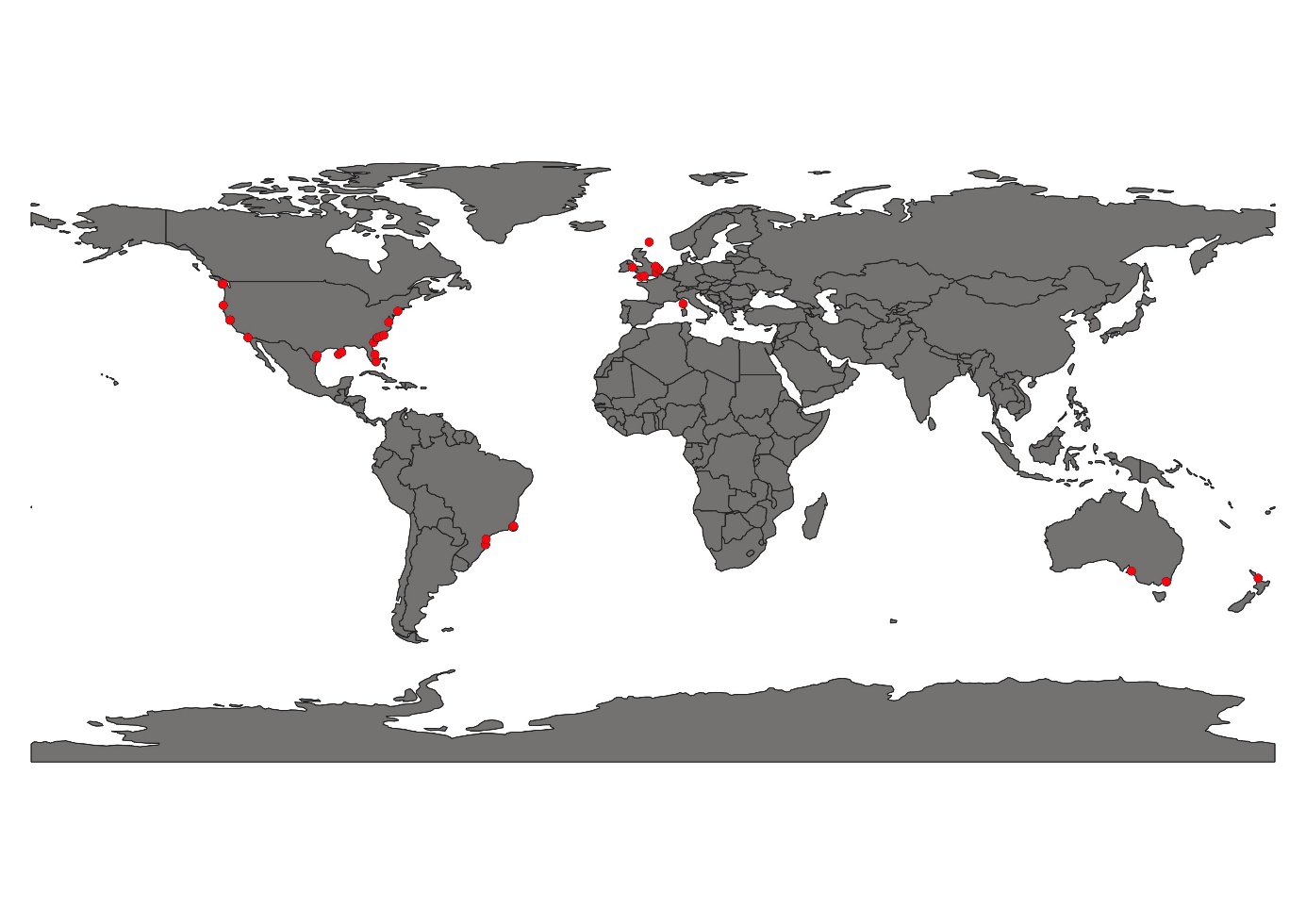

**Fig S7.** The geographical position of *Caprella equilibra* (after thinning the data) was applied for species distribution modeling.

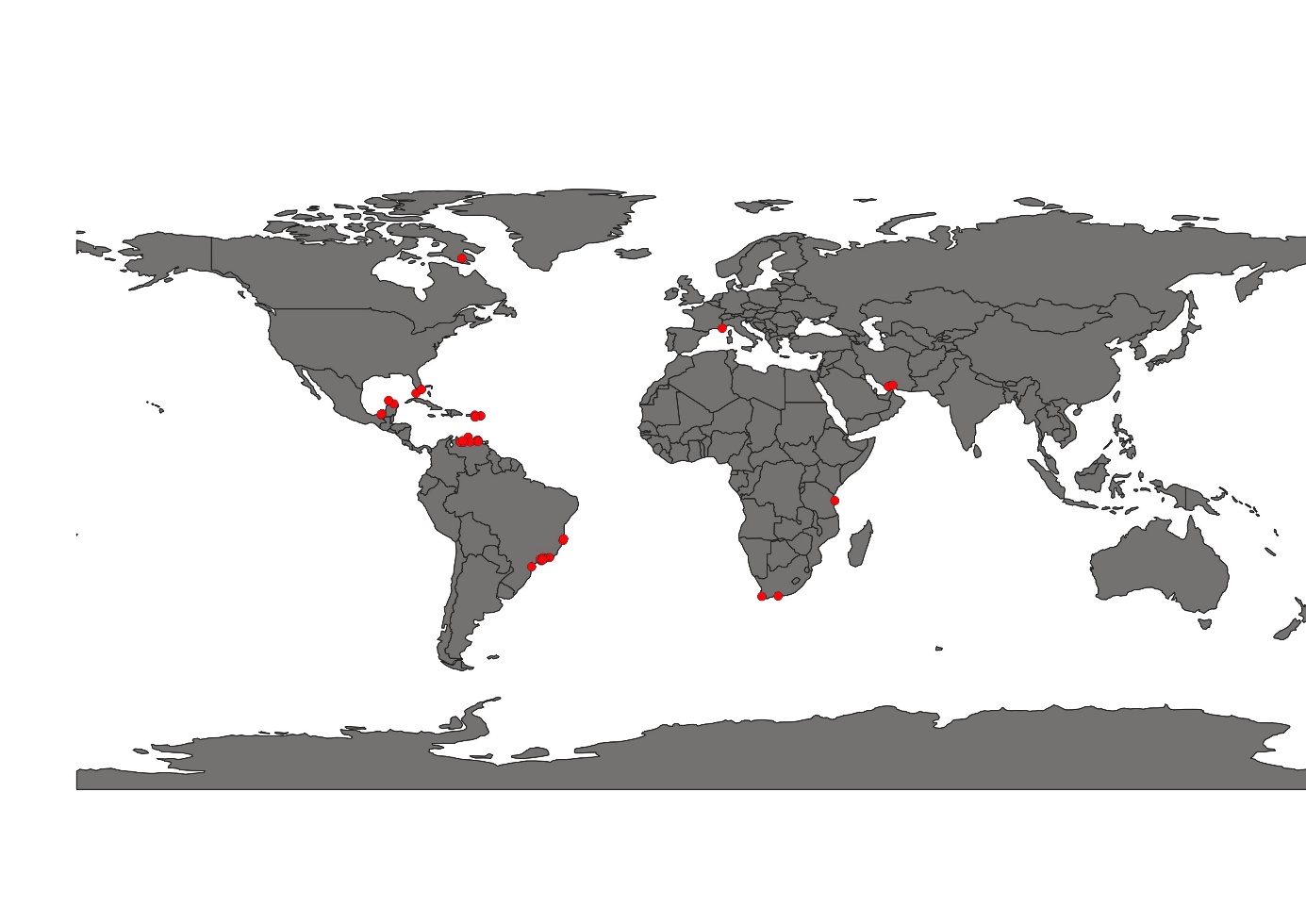

**Fig S8.** The geographical position of *Cymadusa filosa* (after thinning the data) was applied for species distribution modeling.

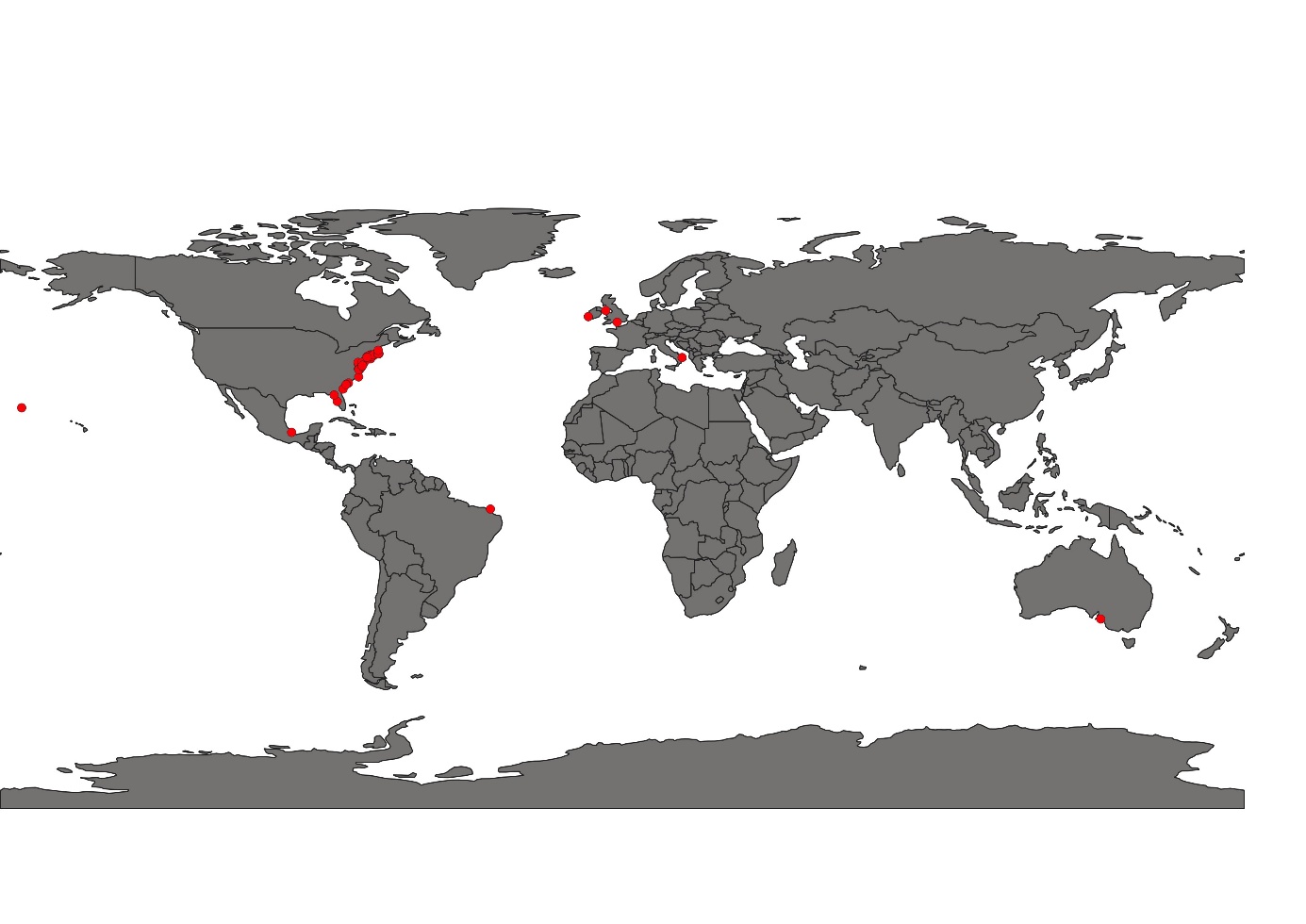

**Fig S9.** The geographical position of *Caprella penantis* (after thinning the data) was applied for species distribution modeling.

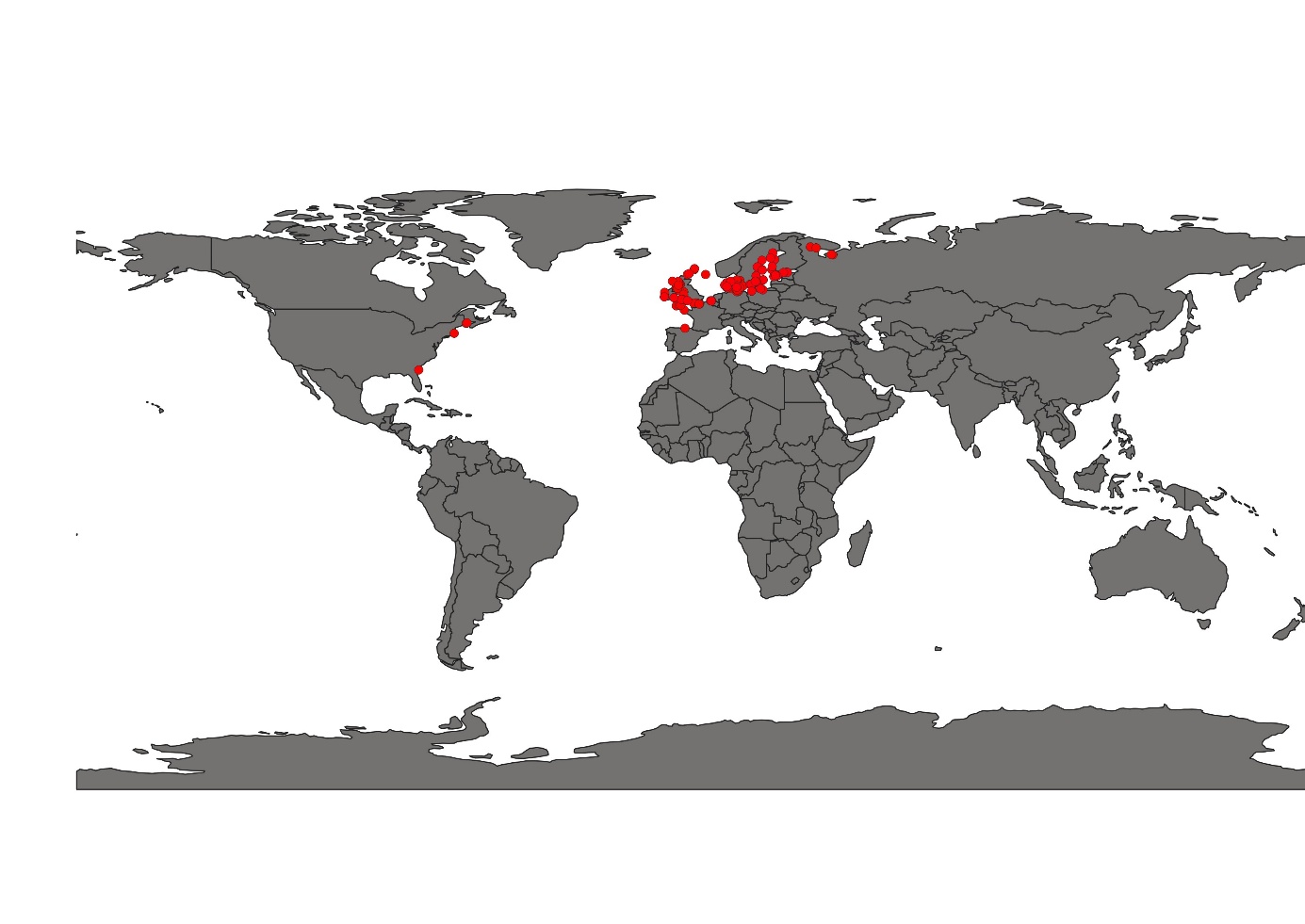

**Fig S10.** The geographical position of *Corophium volutator* (after thinning the data) was applied for species distribution modeling.

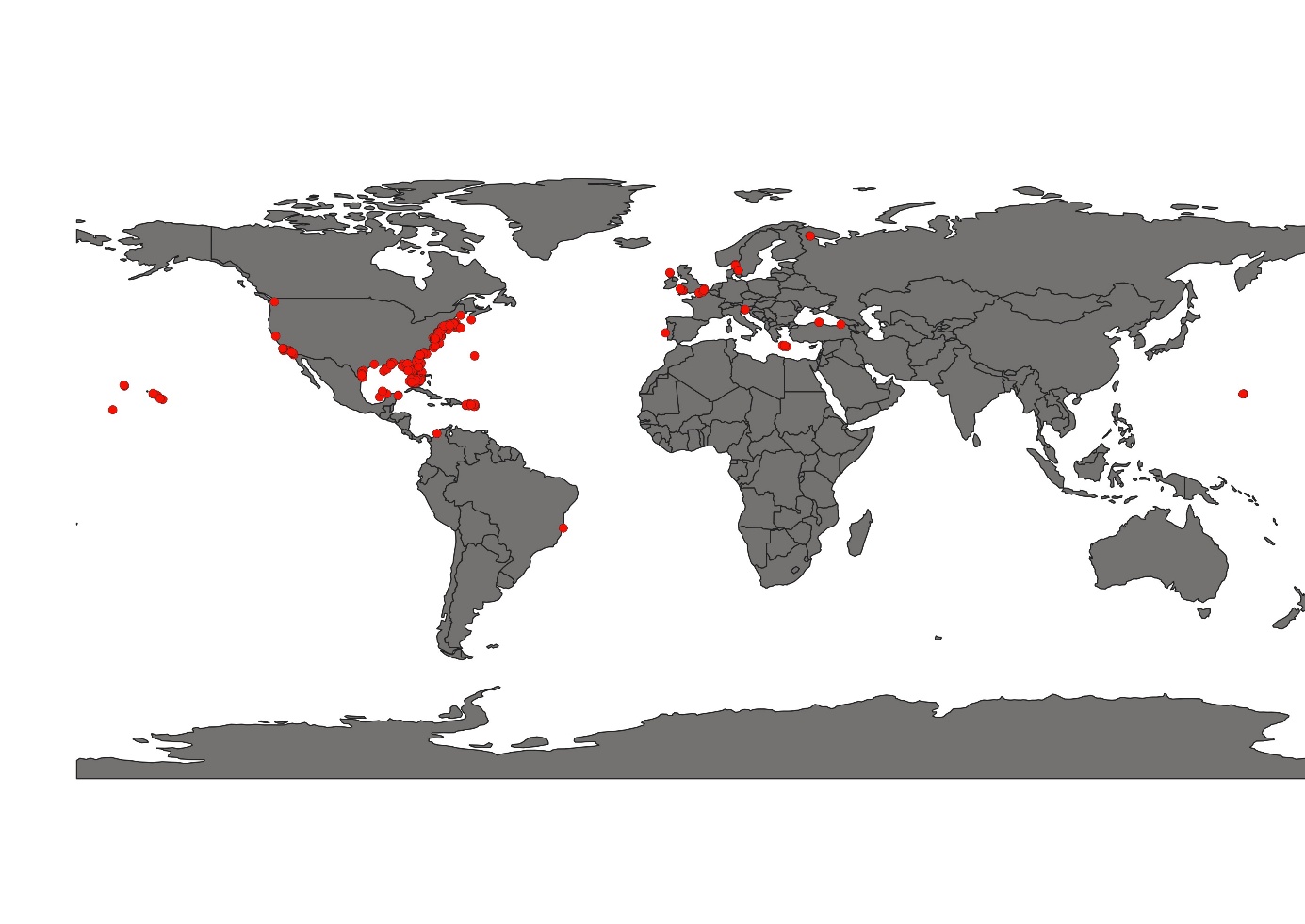

**Fig S11.** The geographical position of *Ericthonius brasiliensis* (after thinning the data) was applied for species distribution modeling.

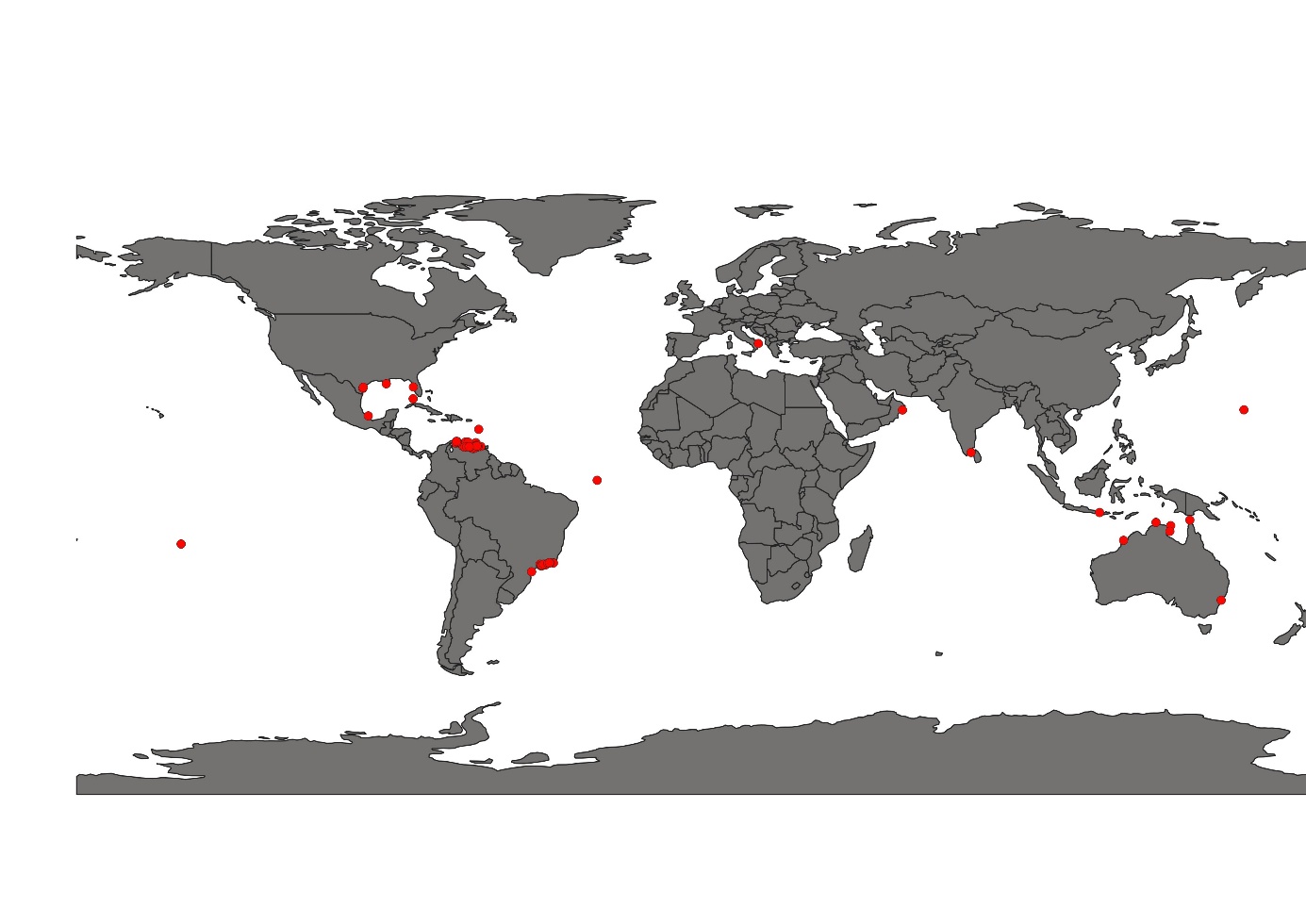

**Fig S12.** The geographical position of *Elasmopus pectenicrus* (after thinning the data) was applied for species distribution modeling.

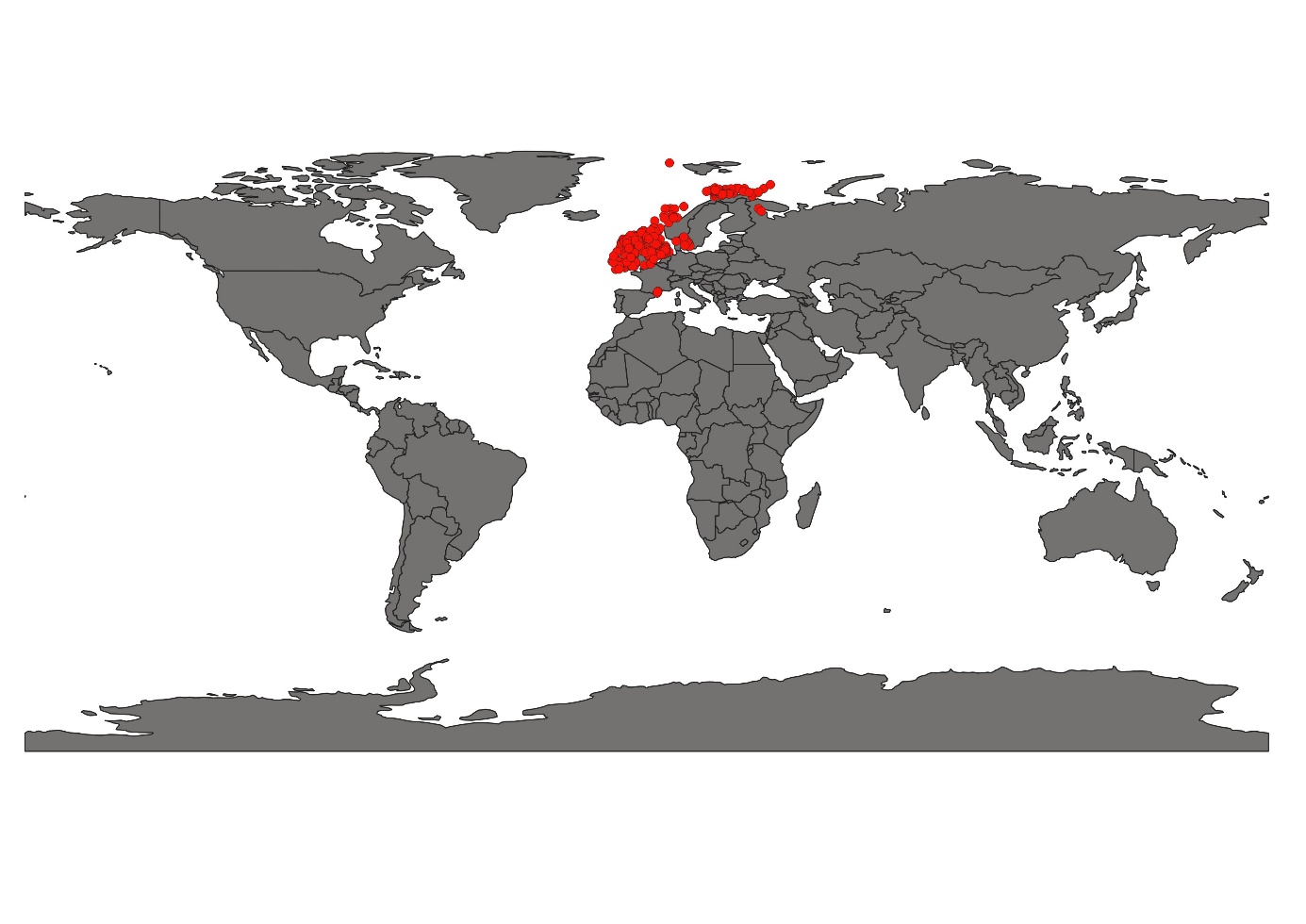

**Fig S12.** The geographical position of *Harpinia antennaria* (after thinning the data) was applied for species distribution modeling.

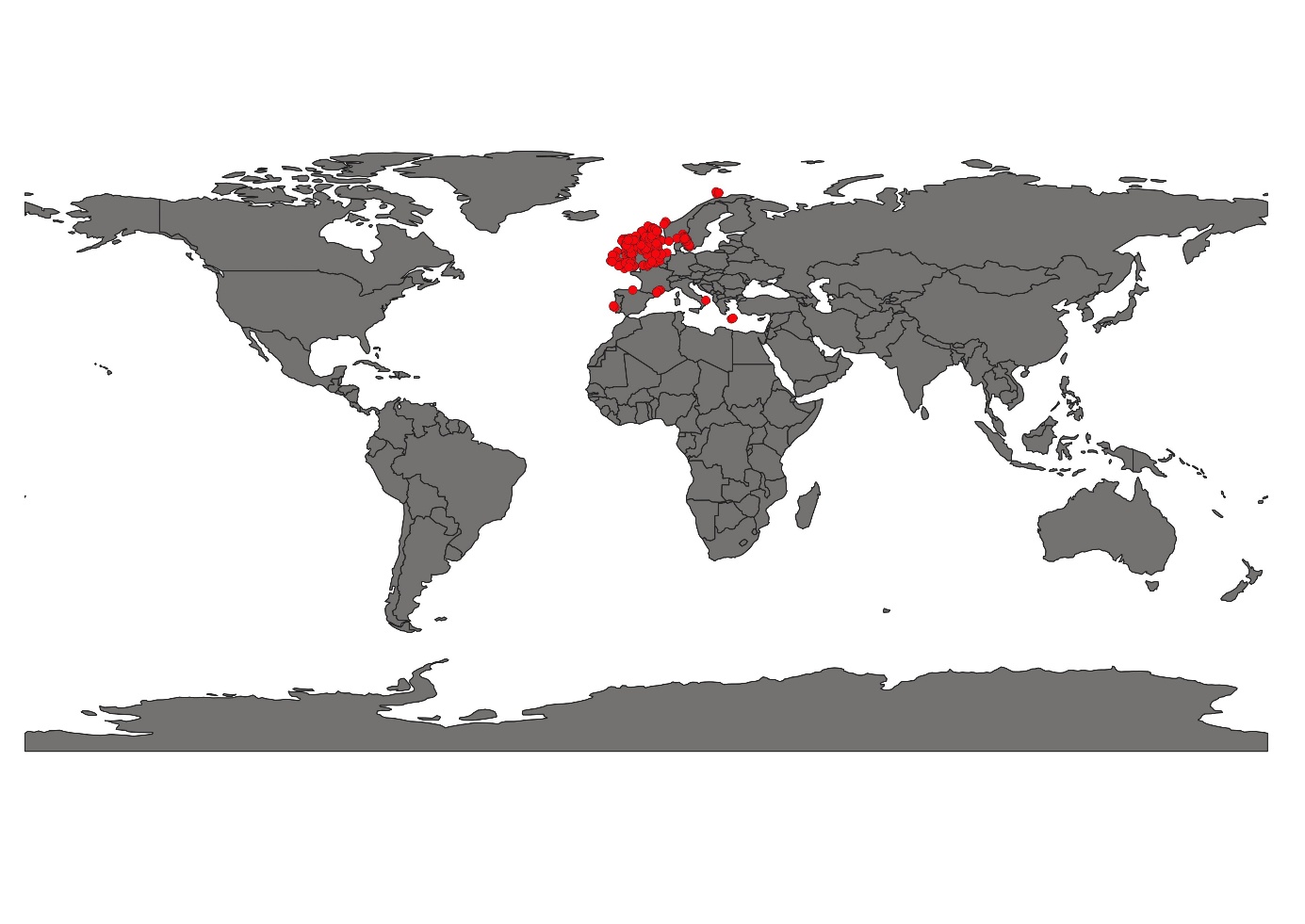

**Fig S13.** The geographical position of *Leucothoe lilljeborgi* (after thinning the data) was applied for species distribution modeling.

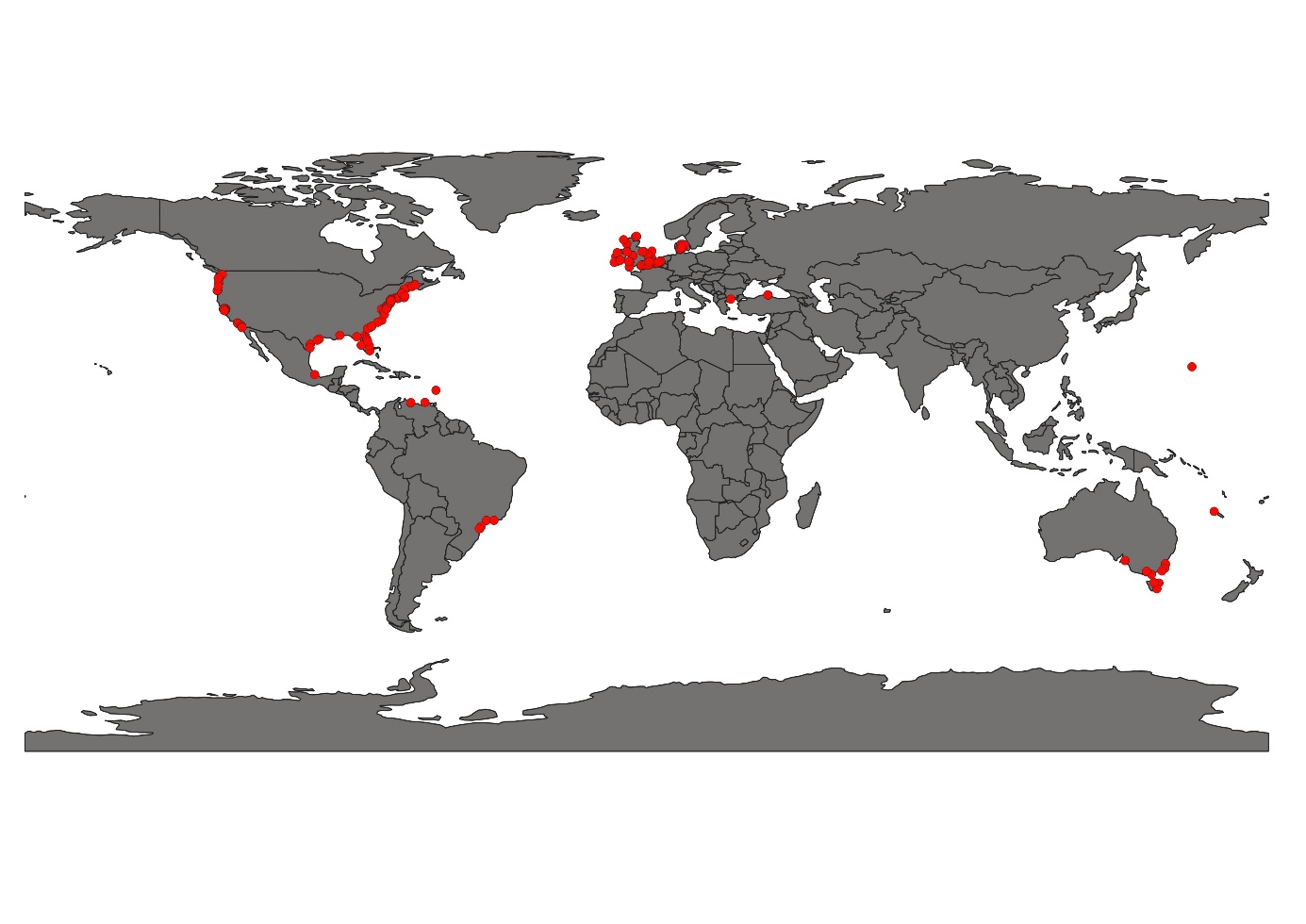

**Fig S14.** The geographical position of *Monocorophium* *acherusicum* (after thinning the data) was applied for species distribution modeling.

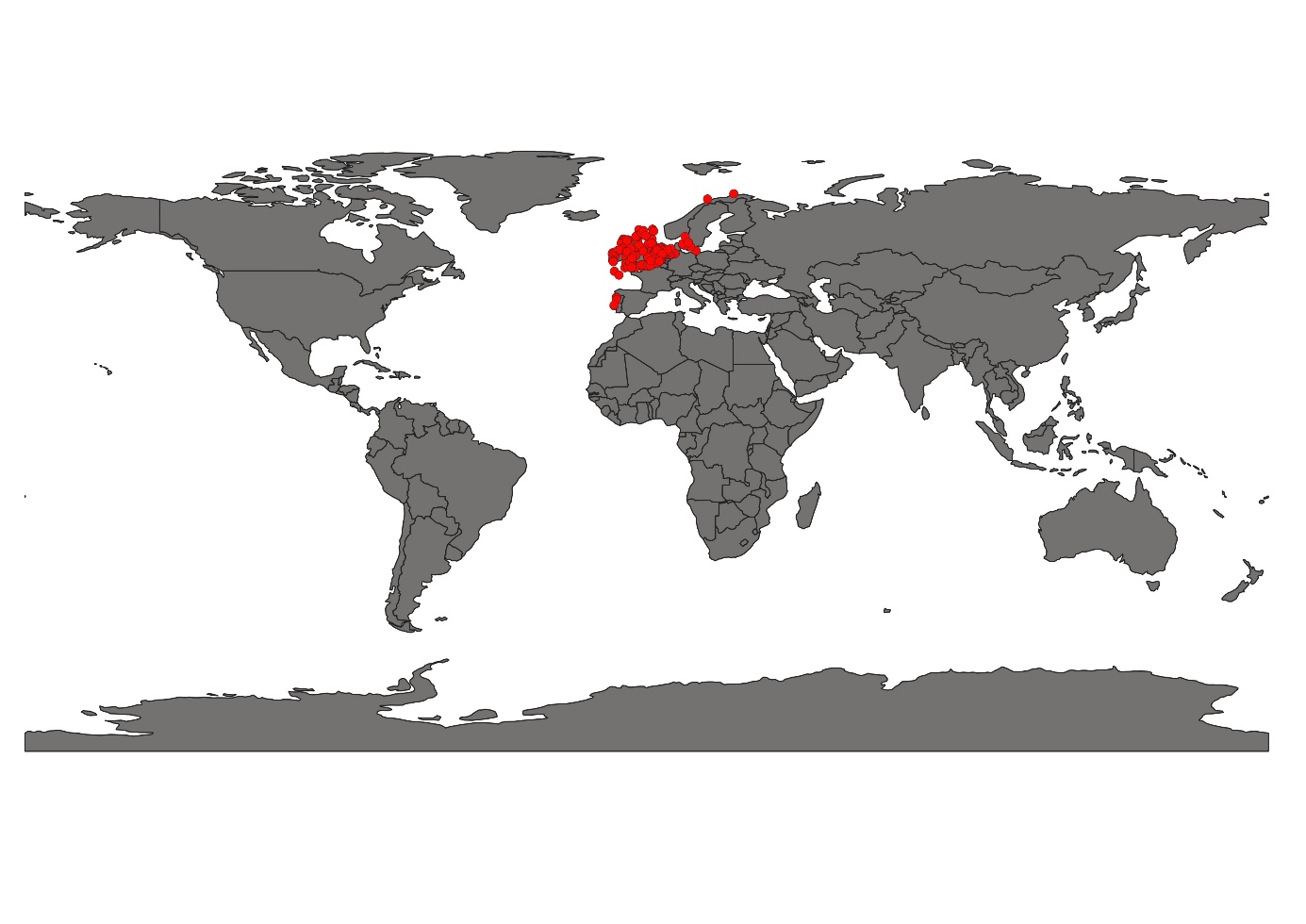

**Fig S15.** The geographical position of *Tryphosa nana* (after thinning the data) was applied for species distribution modeling.

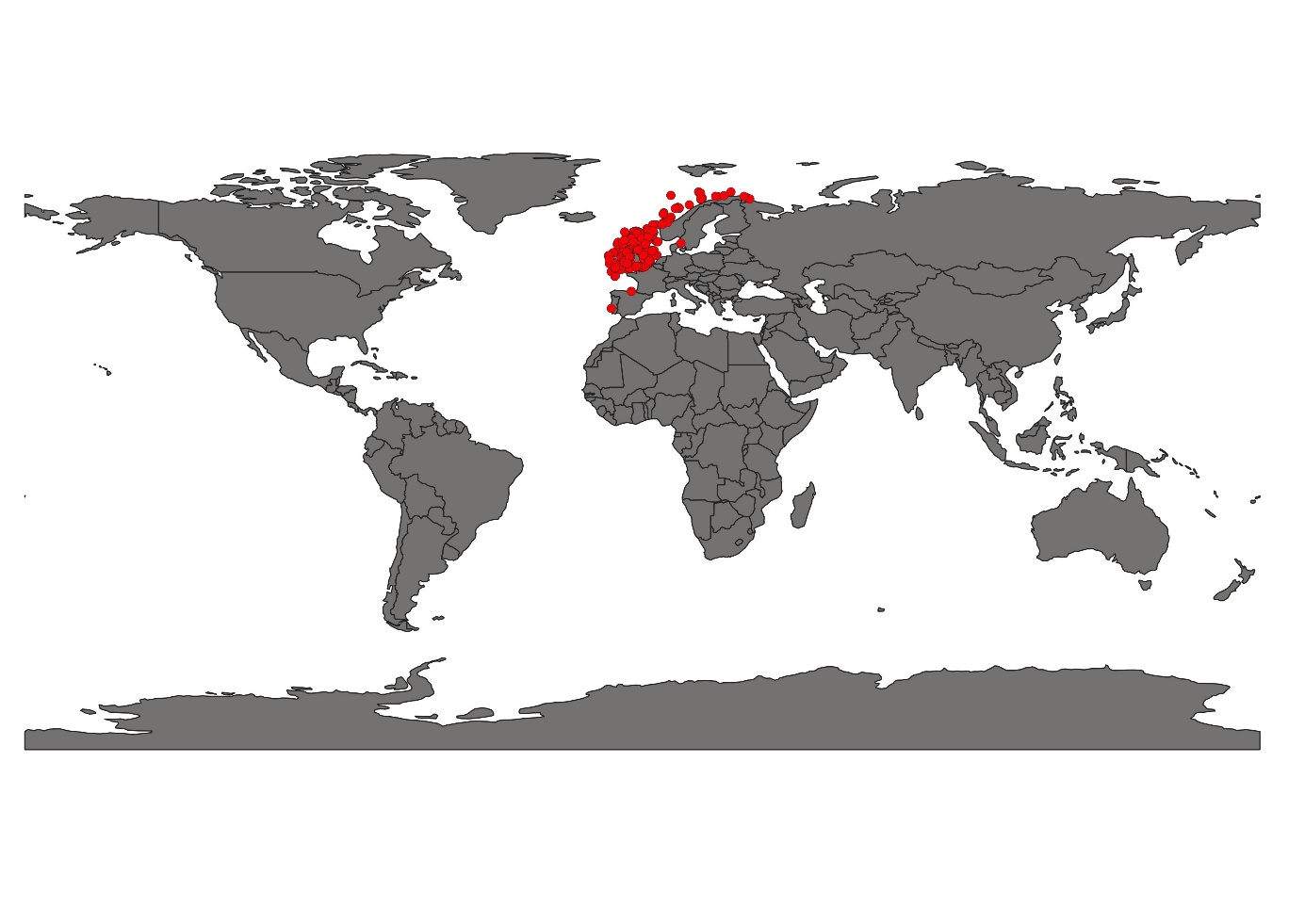

**Fig S16.** The geographical position of *Urothoe elegans* (after thinning the data) was applied for species distribution modeling.

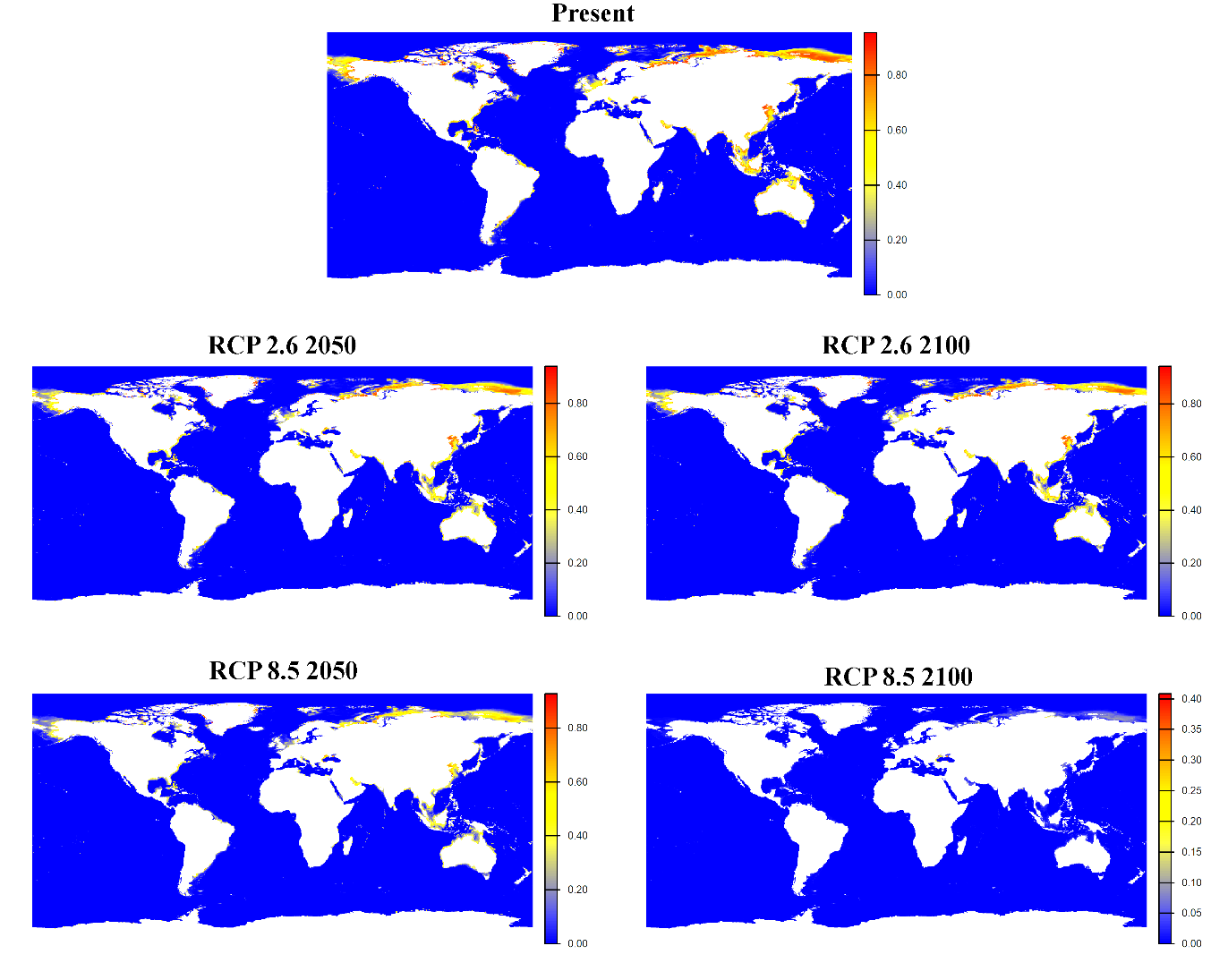

**Fig S17.** Global habitat suitability projections for Ampelisca brevicornis under present and future climate scenarios. The top panel shows the current predicted distribution. Middle panels illustrate mid-century (2040–2050) and end-century (2090–2100) projections under the low-emissions scenario SSP1-2.6. Bottom panels present corresponding projections under the high-emissions scenario SSP5-8.5. Color gradients represent modeled habitat suitability (0–1), with warmer colors (yellow–red) indicating higher suitability and blue indicating low suitability.

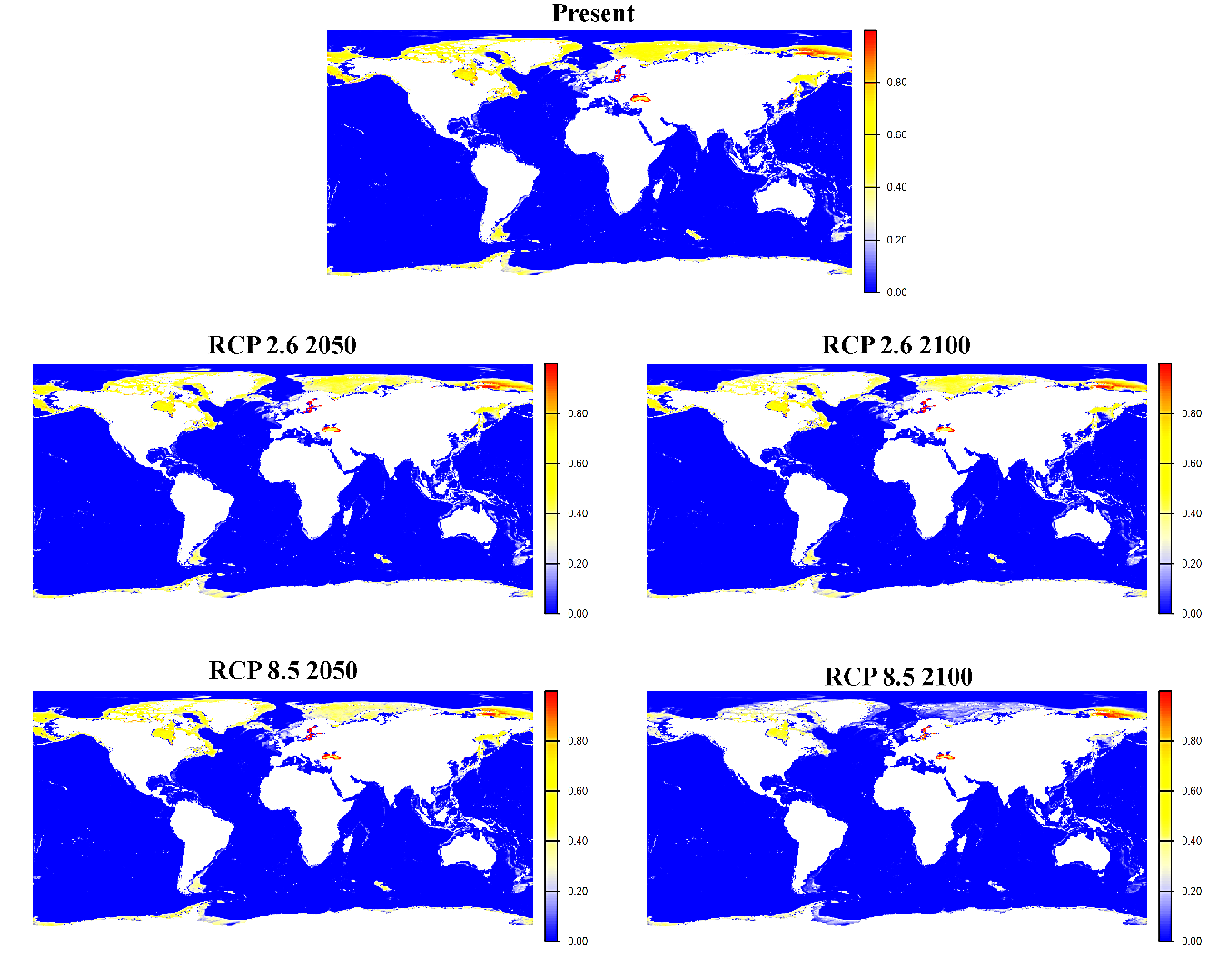

**Fig S18.** Global habitat suitability projections for *Ampelisca eschrichtii* under present and future climate scenarios. The top panel shows the current predicted distribution. Middle panels illustrate mid-century (2040–2050) and end-century (2090–2100) projections under the low-emissions scenario SSP1-2.6. Bottom panels present corresponding projections under the high-emissions scenario SSP5-8.5. Color gradients represent modeled habitat suitability (0–1), with warmer colors (yellow–red) indicating higher suitability and blue indicating low suitability.

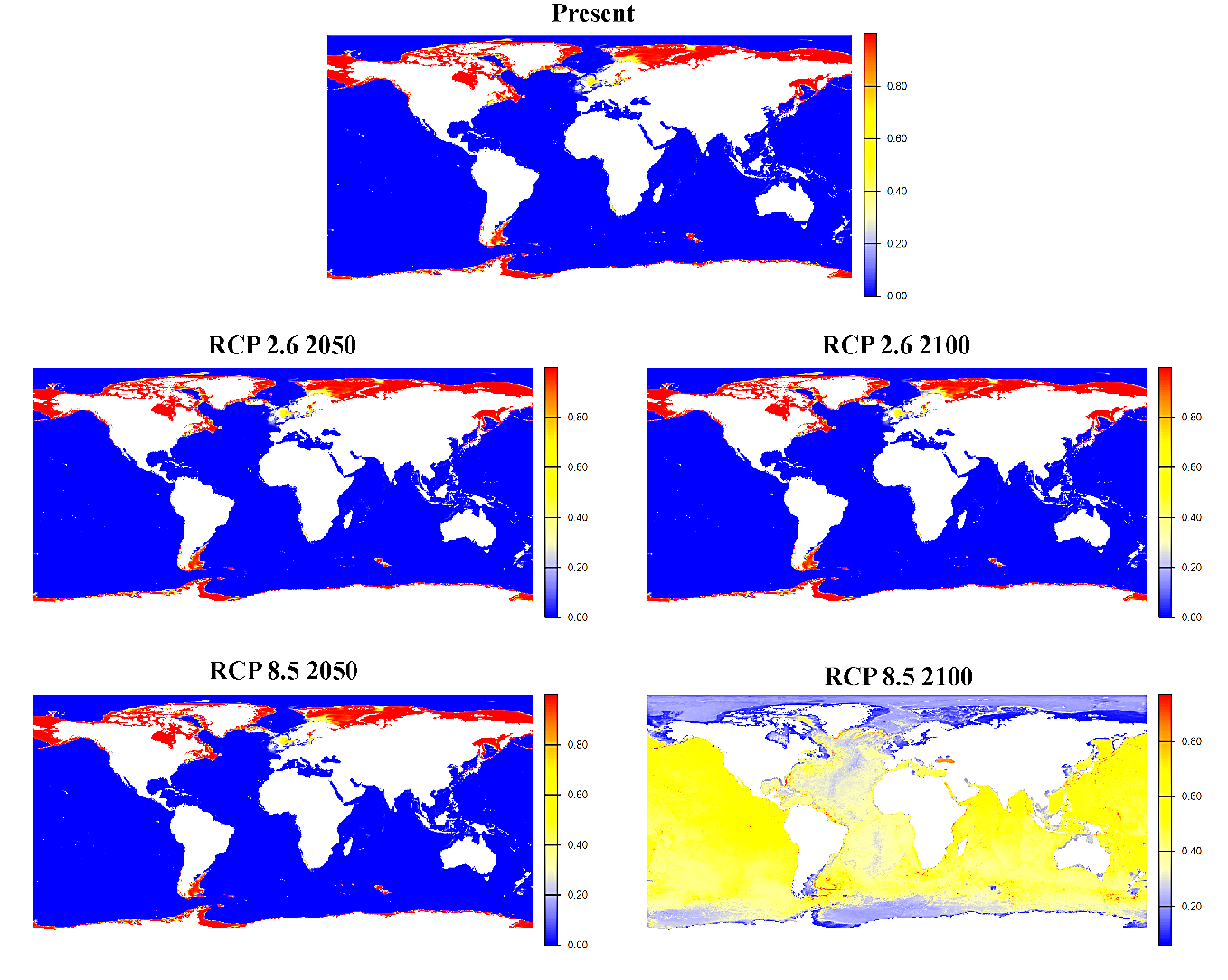

**Fig S19.** Global habitat suitability projections for *Ampelisca macrocephala* under present and future climate scenarios. The top panel shows the current predicted distribution. Middle panels illustrate mid-century (2040–2050) and end-century (2090–2100) projections under the low-emissions scenario SSP1-2.6. Bottom panels present corresponding projections under the high-emissions scenario SSP5-8.5. Color gradients represent modeled habitat suitability (0–1), with warmer colors (yellow–red) indicating higher suitability and blue indicating low suitability.

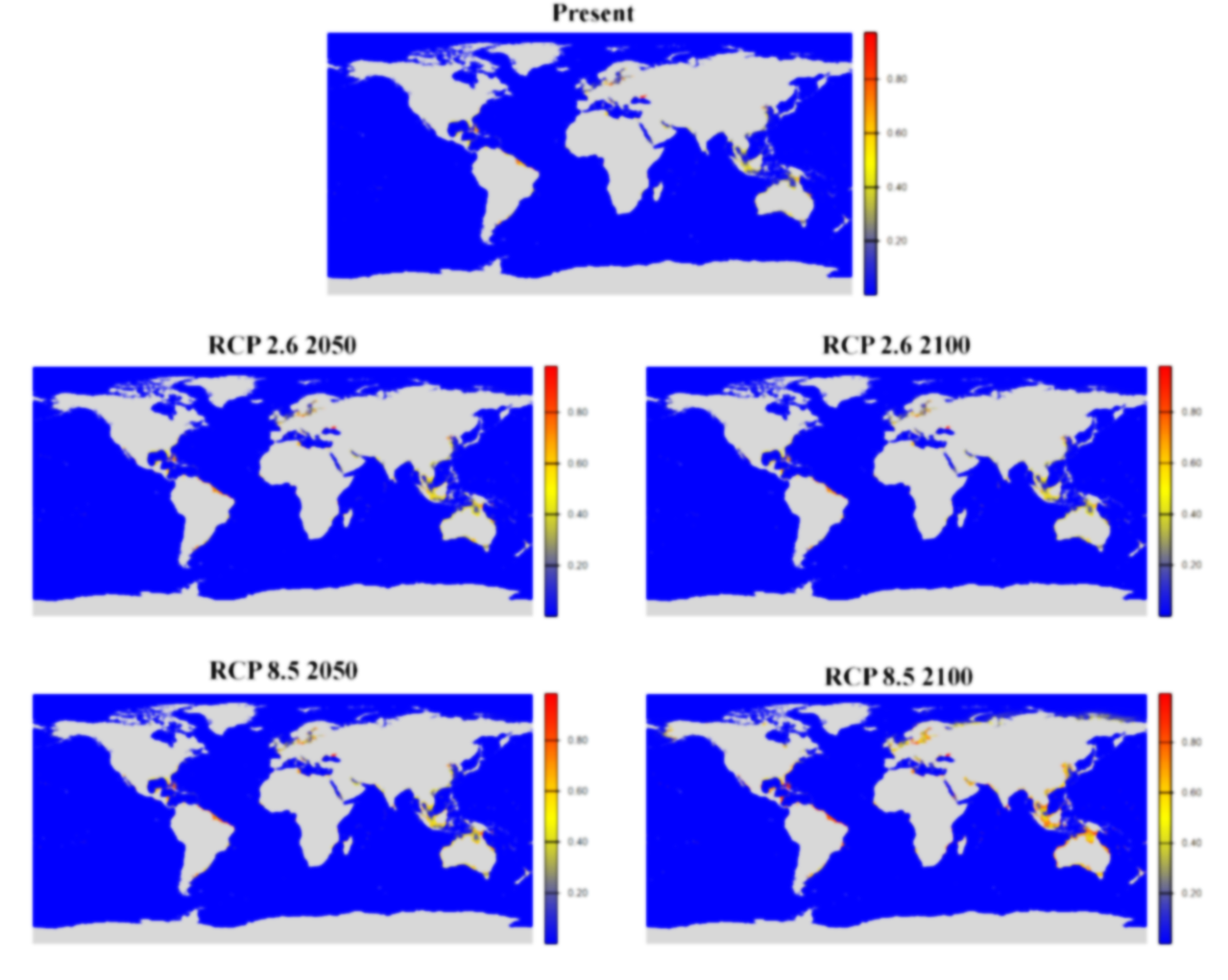

**Fig S20.** Global habitat suitability projections for *Ampithoe rubricata* under present and future climate scenarios. The top panel shows the current predicted distribution. Middle panels illustrate mid-century (2040–2050) and end-century (2090–2100) projections under the low-emissions scenario SSP1-2.6. Bottom panels present corresponding projections under the high-emissions scenario SSP5-8.5. Color gradients represent modeled habitat suitability (0–1), with warmer colors (yellow–red) indicating higher suitability and blue indicating low suitability.

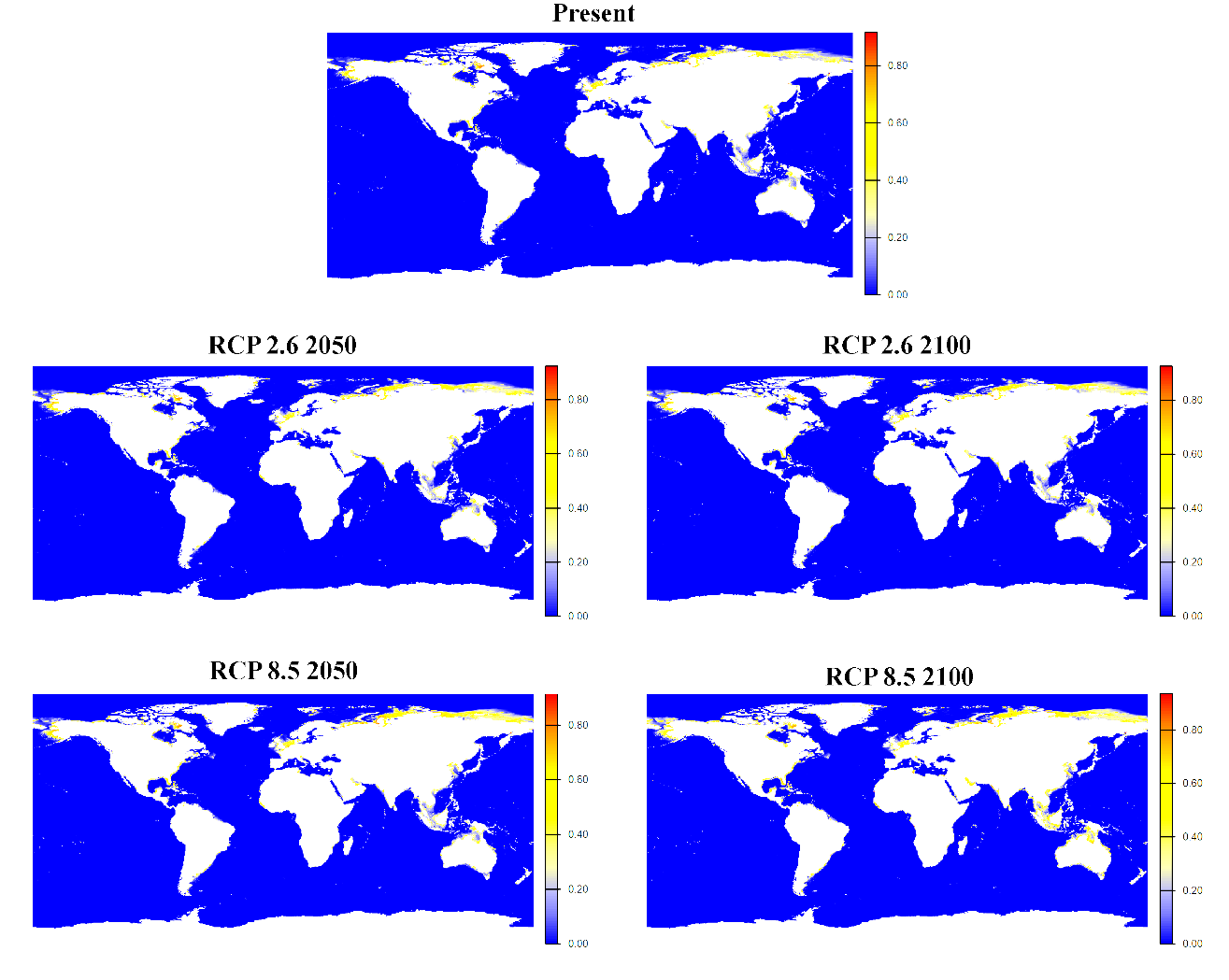

**Fig S21.** Global habitat suitability projections for *Bathyporeia elegans* under present and future climate scenarios. The top panel shows the current predicted distribution. Middle panels illustrate mid-century (2040–2050) and end-century (2090–2100) projections under the low-emissions scenario SSP1-2.6. Bottom panels present corresponding projections under the high-emissions scenario SSP5-8.5. Color gradients represent modeled habitat suitability (0–1), with warmer colors (yellow–red) indicating higher suitability and blue indicating low suitability.

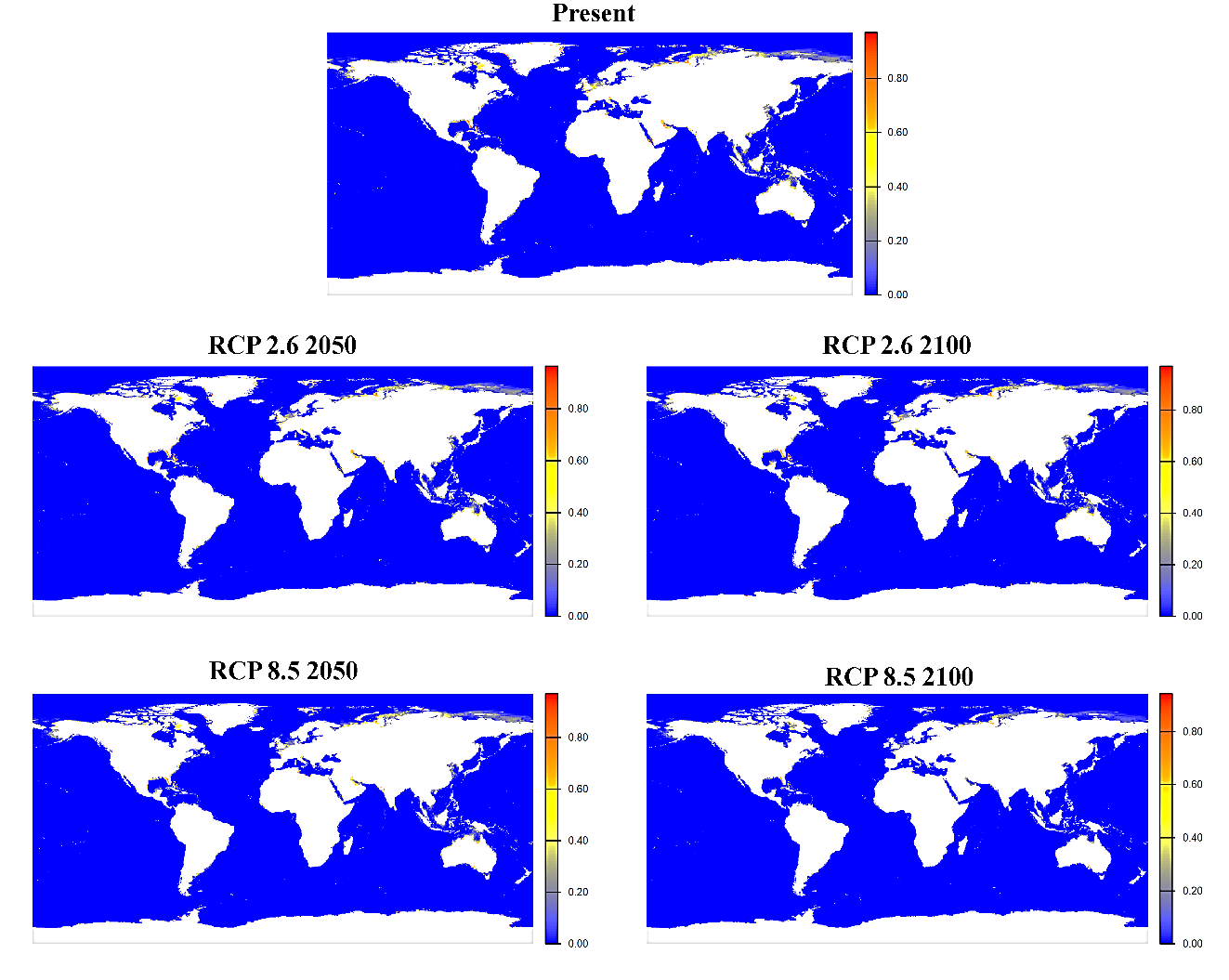

**Fig S22.** Global habitat suitability projections for *Caprella equilibra* under present and future climate scenarios. The top panel shows the current predicted distribution. Middle panels illustrate mid-century (2040–2050) and end-century (2090–2100) projections under the low-emissions scenario SSP1-2.6. Bottom panels present corresponding projections under the high-emissions scenario SSP5-8.5. Color gradients represent modeled habitat suitability (0–1), with warmer colors (yellow–red) indicating higher suitability and blue indicating low suitability.

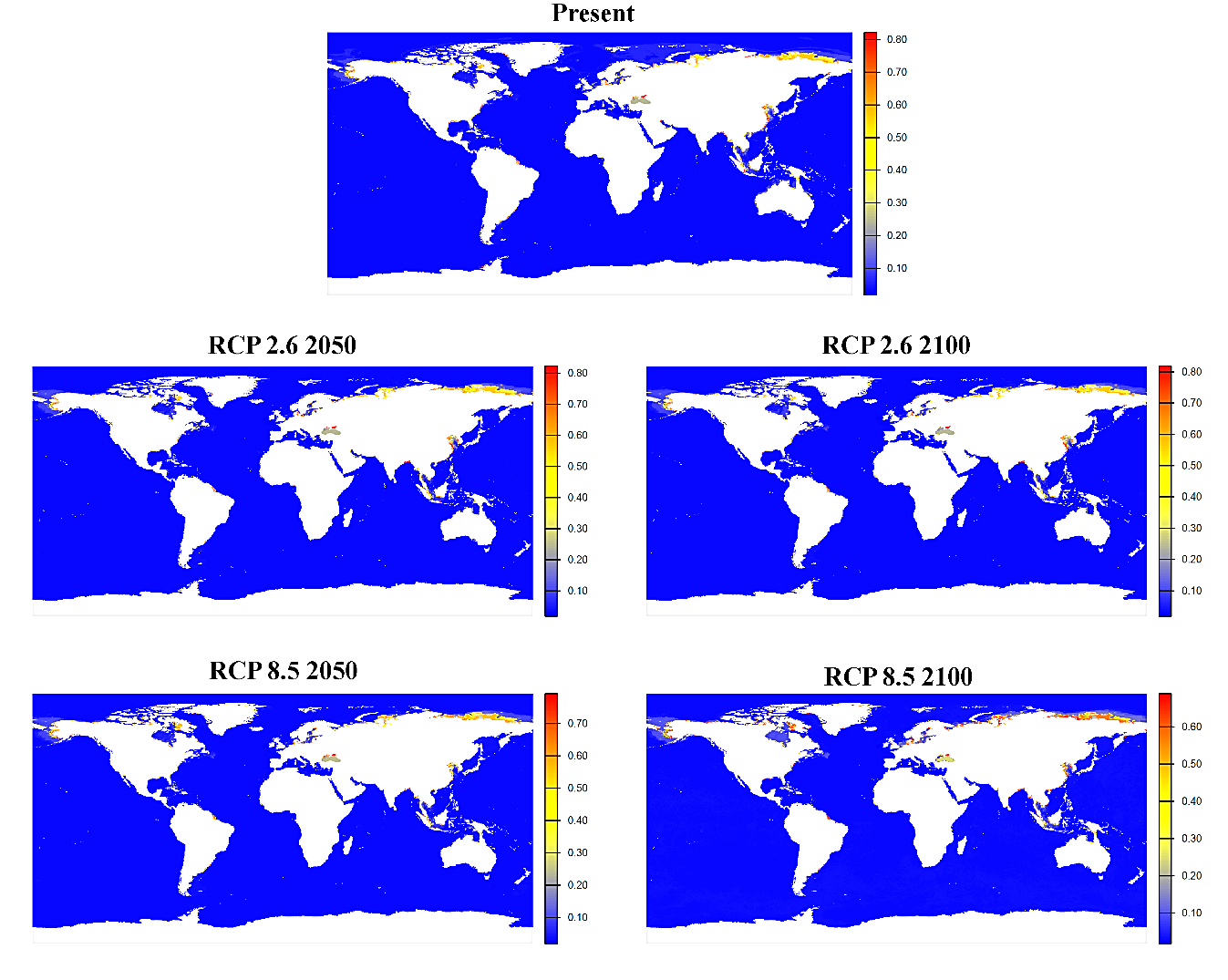

**Fig S23.** Global habitat suitability projections for *Caprella penantis* under present and future climate scenarios. The top panel shows the current predicted distribution. Middle panels illustrate mid-century (2040–2050) and end-century (2090–2100) projections under the low-emissions scenario SSP1-2.6. Bottom panels present corresponding projections under the high-emissions scenario SSP5-8.5. Color gradients represent modeled habitat suitability (0–1), with warmer colors (yellow–red) indicating higher suitability and blue indicating low suitability.

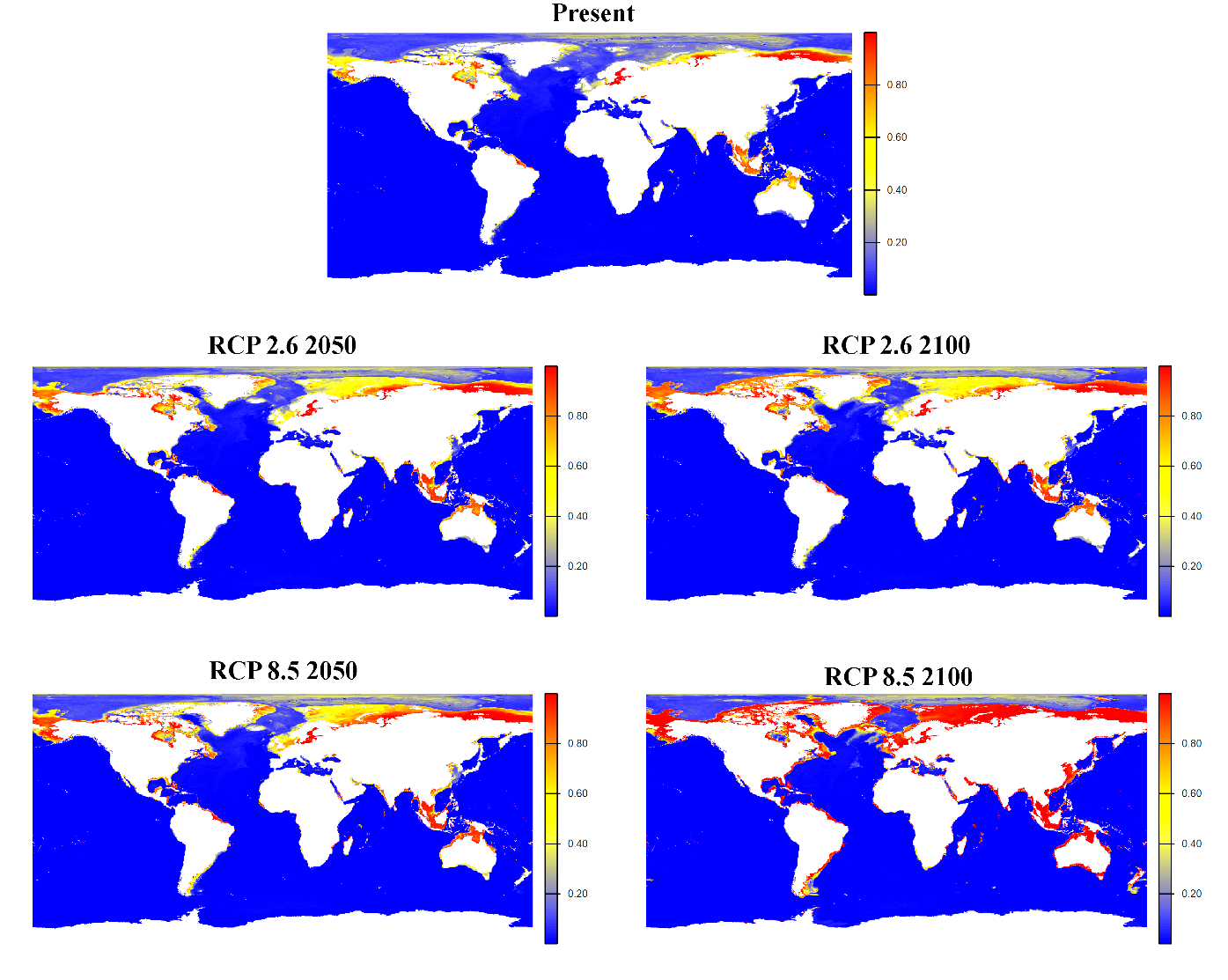

**Fig S24.** Global habitat suitability projections for *Corophium volutator* under present and future climate scenarios. The top panel shows the current predicted distribution. Middle panels illustrate mid-century (2040–2050) and end-century (2090–2100) projections under the low-emissions scenario SSP1-2.6. Bottom panels present corresponding projections under the high-emissions scenario SSP5-8.5. Color gradients represent modeled habitat suitability (0–1), with warmer colors (yellow–red) indicating higher suitability and blue indicating low suitability.

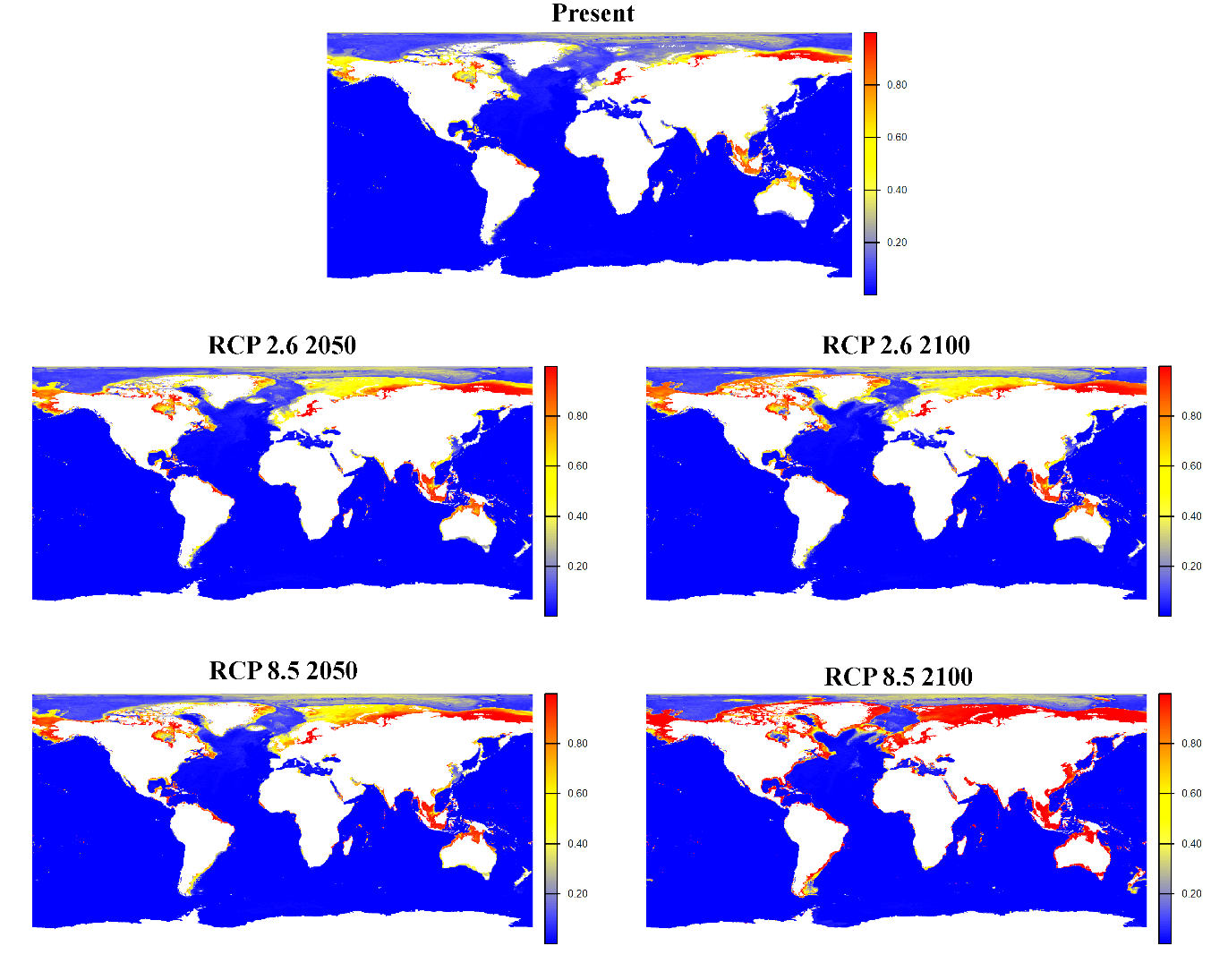

**Fig S25.** Global habitat suitability projections for *Cymadusa compta* under present and future climate scenarios. The top panel shows the current predicted distribution. Middle panels illustrate mid-century (2040–2050) and end-century (2090–2100) projections under the low-emissions scenario SSP1-2.6. Bottom panels present corresponding projections under the high-emissions scenario SSP5-8.5. Color gradients represent modeled habitat suitability (0–1), with warmer colors (yellow–red) indicating higher suitability and blue indicating low suitability.

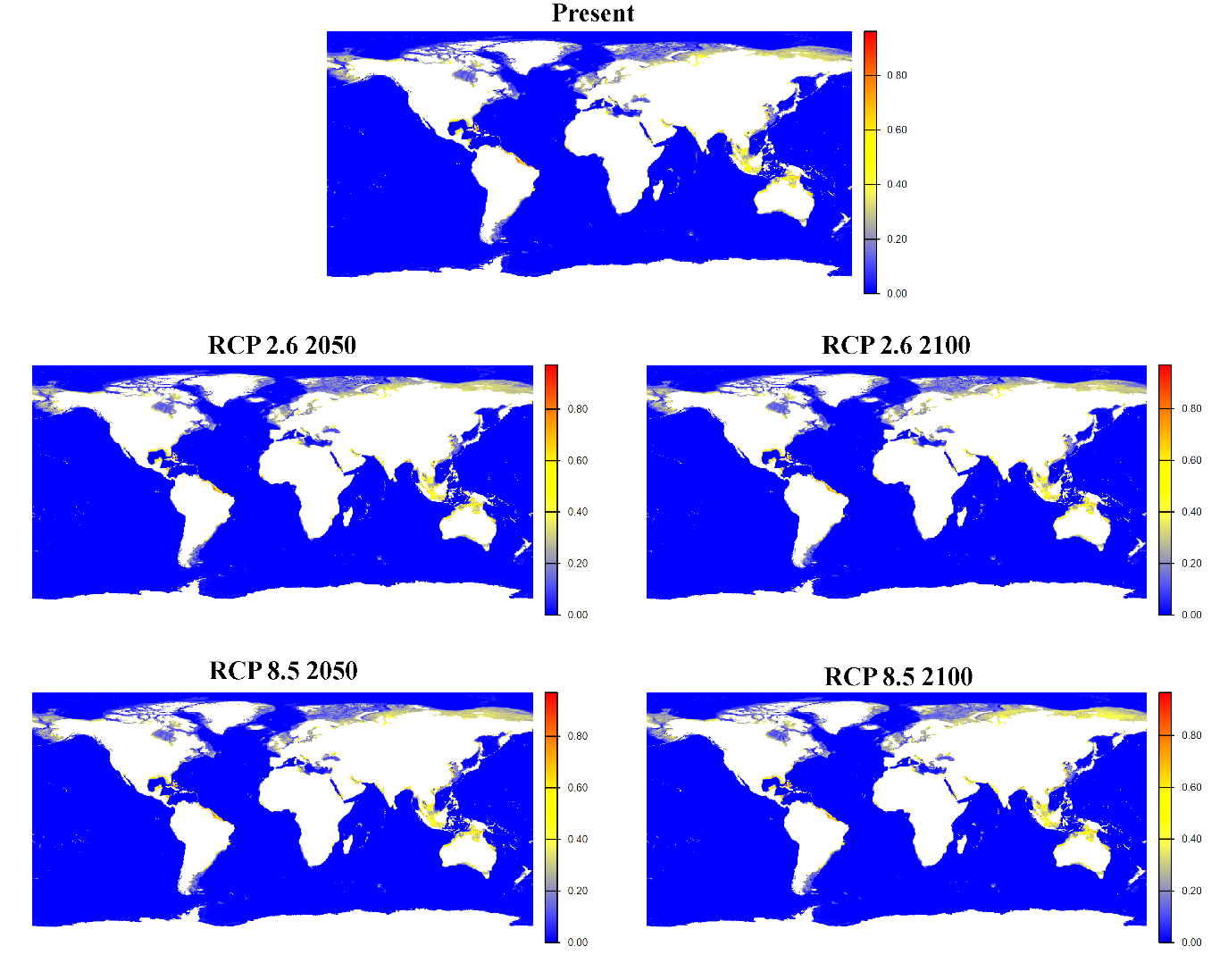

**Fig S26.** Global habitat suitability projections for *Cymadusa filosa* under present and future climate scenarios. The top panel shows the current predicted distribution. Middle panels illustrate mid-century (2040–2050) and end-century (2090–2100) projections under the low-emissions scenario SSP1-2.6. Bottom panels present corresponding projections under the high-emissions scenario SSP5-8.5. Color gradients represent modeled habitat suitability (0–1), with warmer colors (yellow–red) indicating higher suitability and blue indicating low suitability.

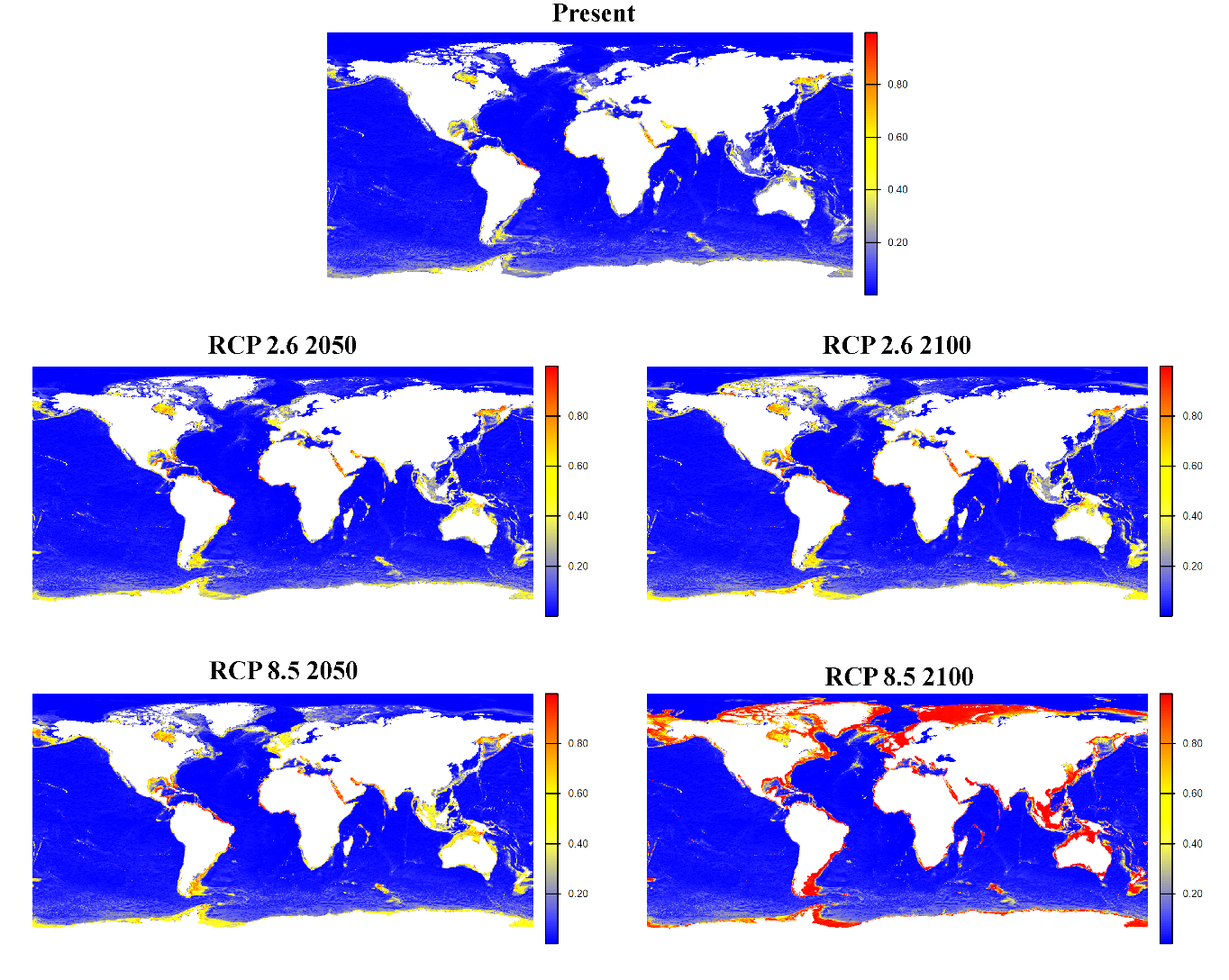

**Fig S27.** Global habitat suitability projections for *Elasmopus pectenicrus* under present and future climate scenarios. The top panel shows the current predicted distribution. Middle panels illustrate mid-century (2040–2050) and end-century (2090–2100) projections under the low-emissions scenario SSP1-2.6. Bottom panels present corresponding projections under the high-emissions scenario SSP5-8.5. Color gradients represent modeled habitat suitability (0–1), with warmer colors (yellow–red) indicating higher suitability and blue indicating low suitability.

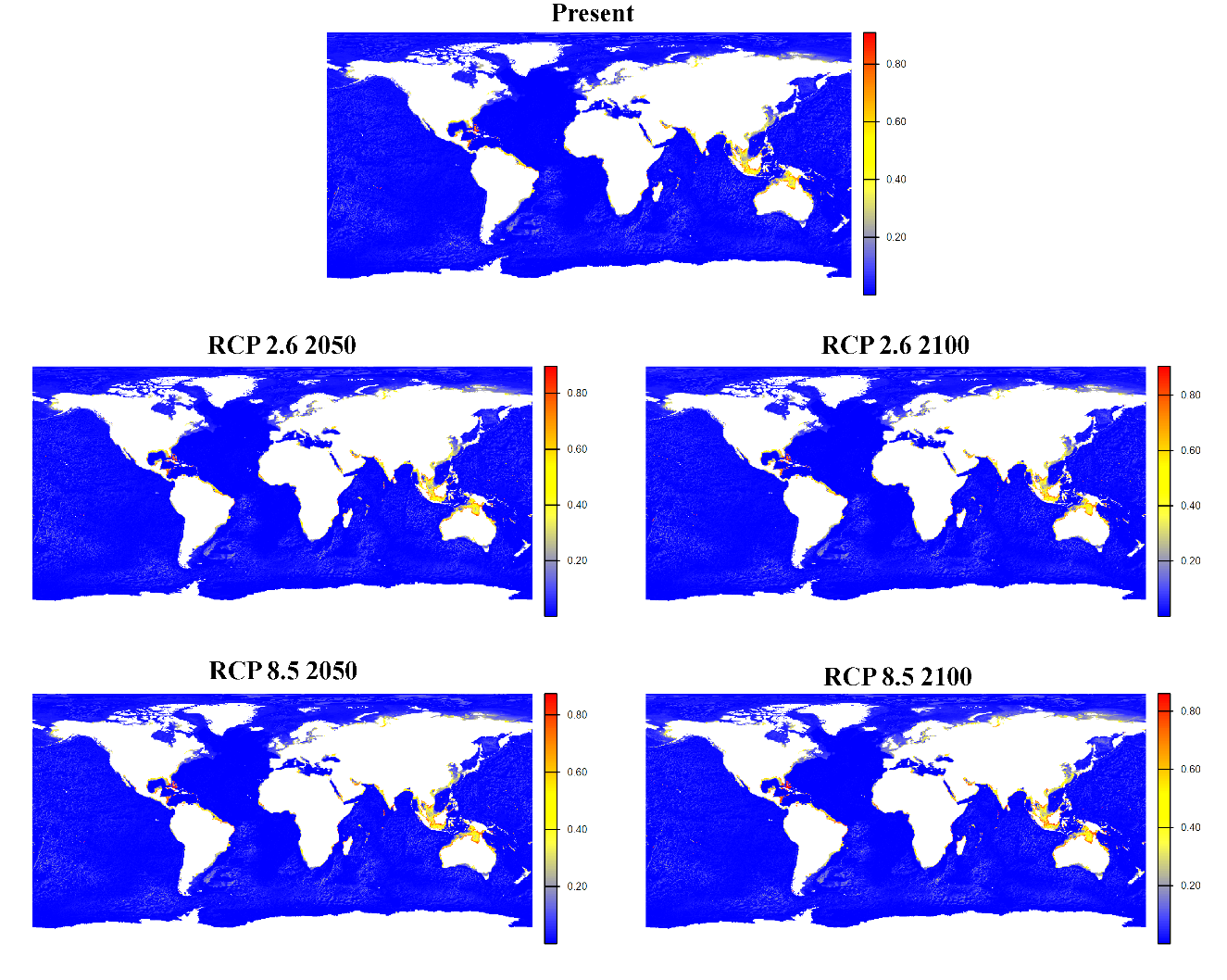

**Fig S28.** Global habitat suitability projections for *Ericthonius brasiliensis* under present and future climate scenarios. The top panel shows the current predicted distribution. Middle panels illustrate mid-century (2040–2050) and end-century (2090–2100) projections under the low-emissions scenario SSP1-2.6. Bottom panels present corresponding projections under the high-emissions scenario SSP5-8.5. Color gradients represent modeled habitat suitability (0–1), with warmer colors (yellow–red) indicating higher suitability and blue indicating low suitability.

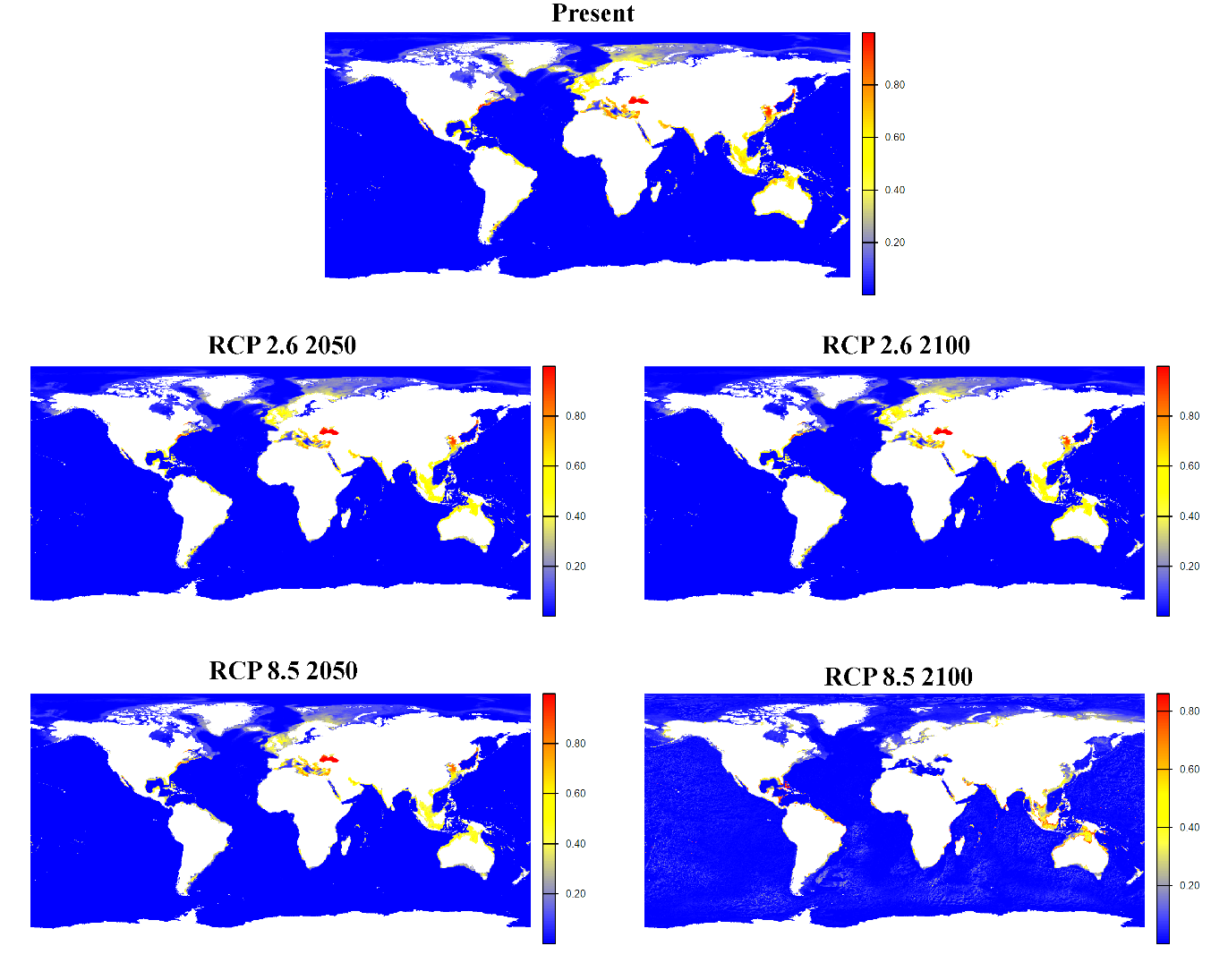

**Fig S29.** Global habitat suitability projections for *Harpinia antennaria* under present and future climate scenarios. The top panel shows the current predicted distribution. Middle panels illustrate mid-century (2040–2050) and end-century (2090–2100) projections under the low-emissions scenario SSP1-2.6. Bottom panels present corresponding projections under the high-emissions scenario SSP5-8.5. Color gradients represent modeled habitat suitability (0–1), with warmer colors (yellow–red) indicating higher suitability and blue indicating low suitability.

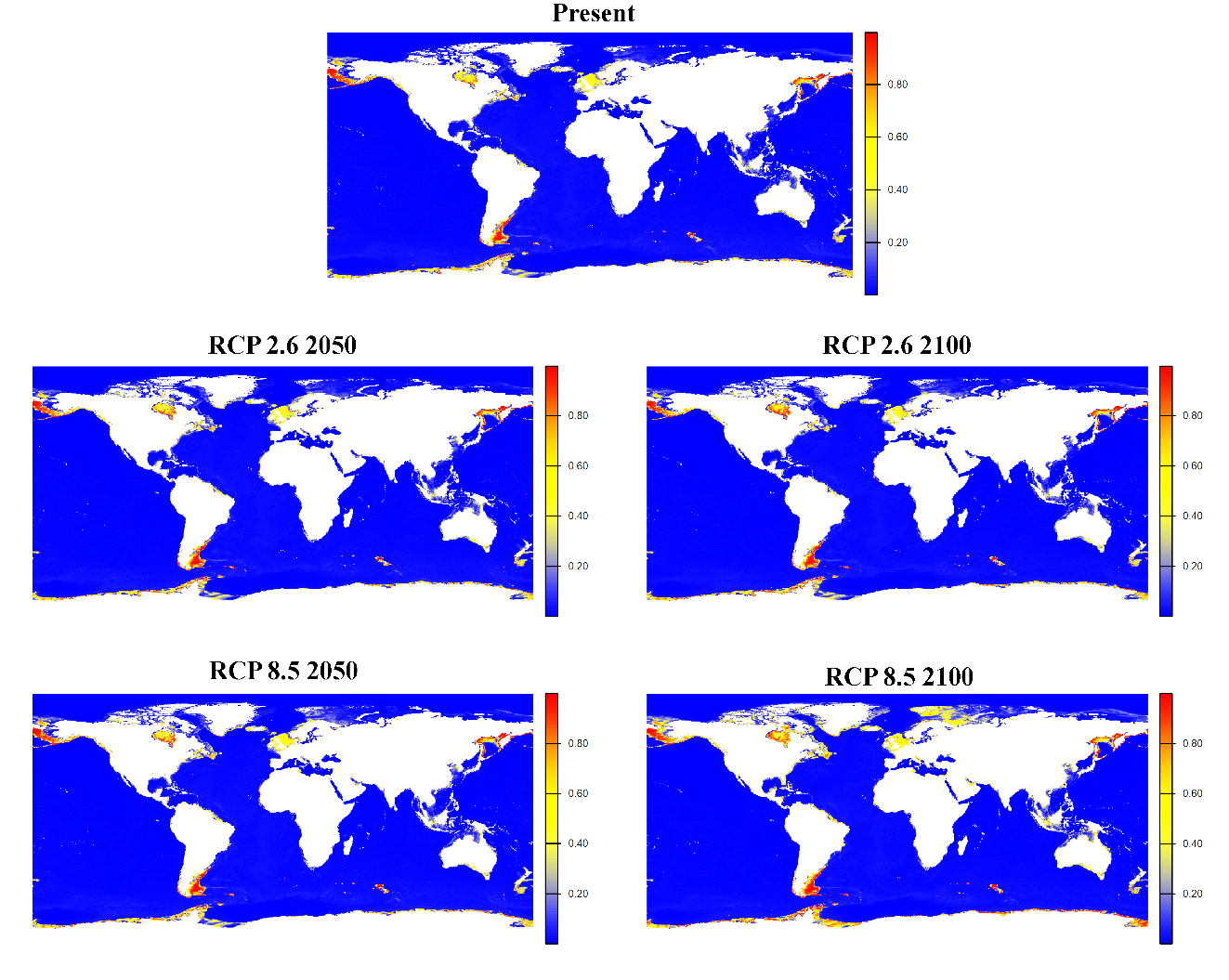

**Fig S30.** Global habitat suitability projections for *Leucothoe lilljeborgi* under present and future climate scenarios. The top panel shows the current predicted distribution. Middle panels illustrate mid-century (2040–2050) and end-century (2090–2100) projections under the low-emissions scenario SSP1-2.6. Bottom panels present corresponding projections under the high-emissions scenario SSP5-8.5. Color gradients represent modeled habitat suitability (0–1), with warmer colors (yellow–red) indicating higher suitability and blue indicating low suitability.

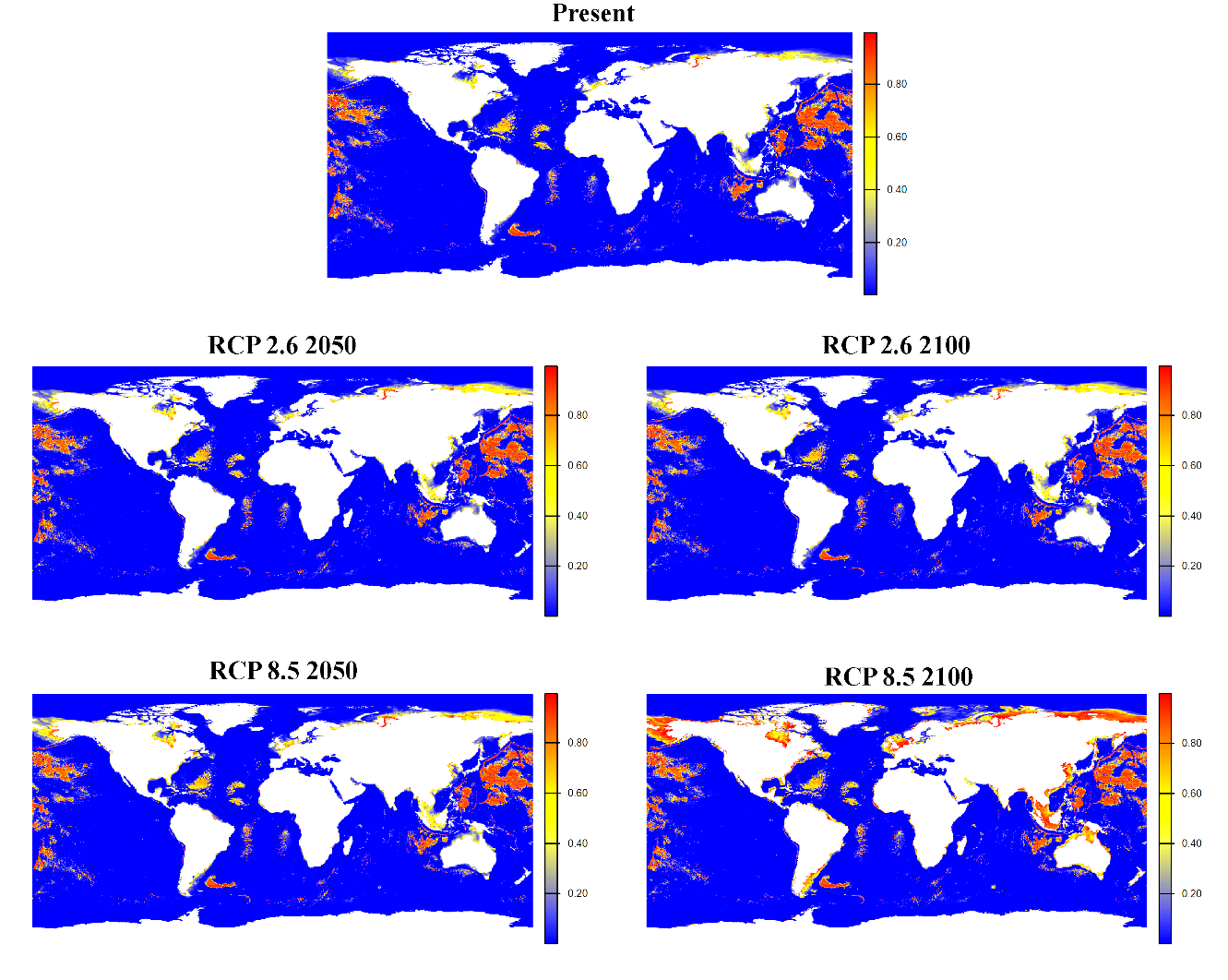

**Fig S31.** Global habitat suitability projections for *Monocorophium* *acherusicum* under present and future climate scenarios. The top panel shows the current predicted distribution. Middle panels illustrate mid-century (2040–2050) and end-century (2090–2100) projections under the low-emissions scenario SSP1-2.6. Bottom panels present corresponding projections under the high-emissions scenario SSP5-8.5. Color gradients represent modeled habitat suitability (0–1), with warmer colors (yellow–red) indicating higher suitability and blue indicating low suitability.

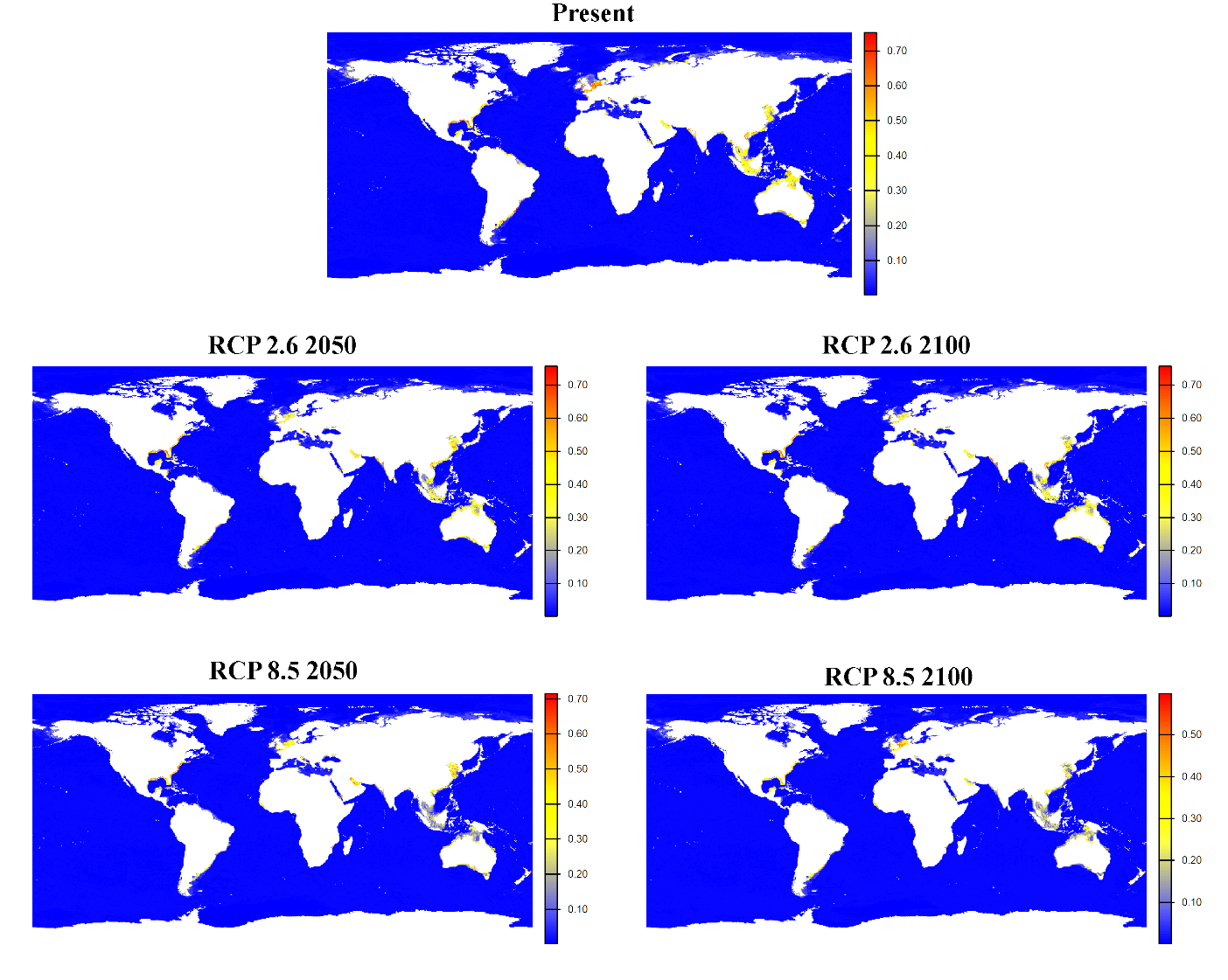

**Fig S32.** Global habitat suitability projections for *Tryphosa nana* under present and future climate scenarios. The top panel shows the current predicted distribution. Middle panels illustrate mid-century (2040–2050) and end-century (2090–2100) projections under the low-emissions scenario SSP1-2.6. Bottom panels present corresponding projections under the high-emissions scenario SSP5-8.5. Color gradients represent modeled habitat suitability (0–1), with warmer colors (yellow–red) indicating higher suitability and blue indicating low suitability.

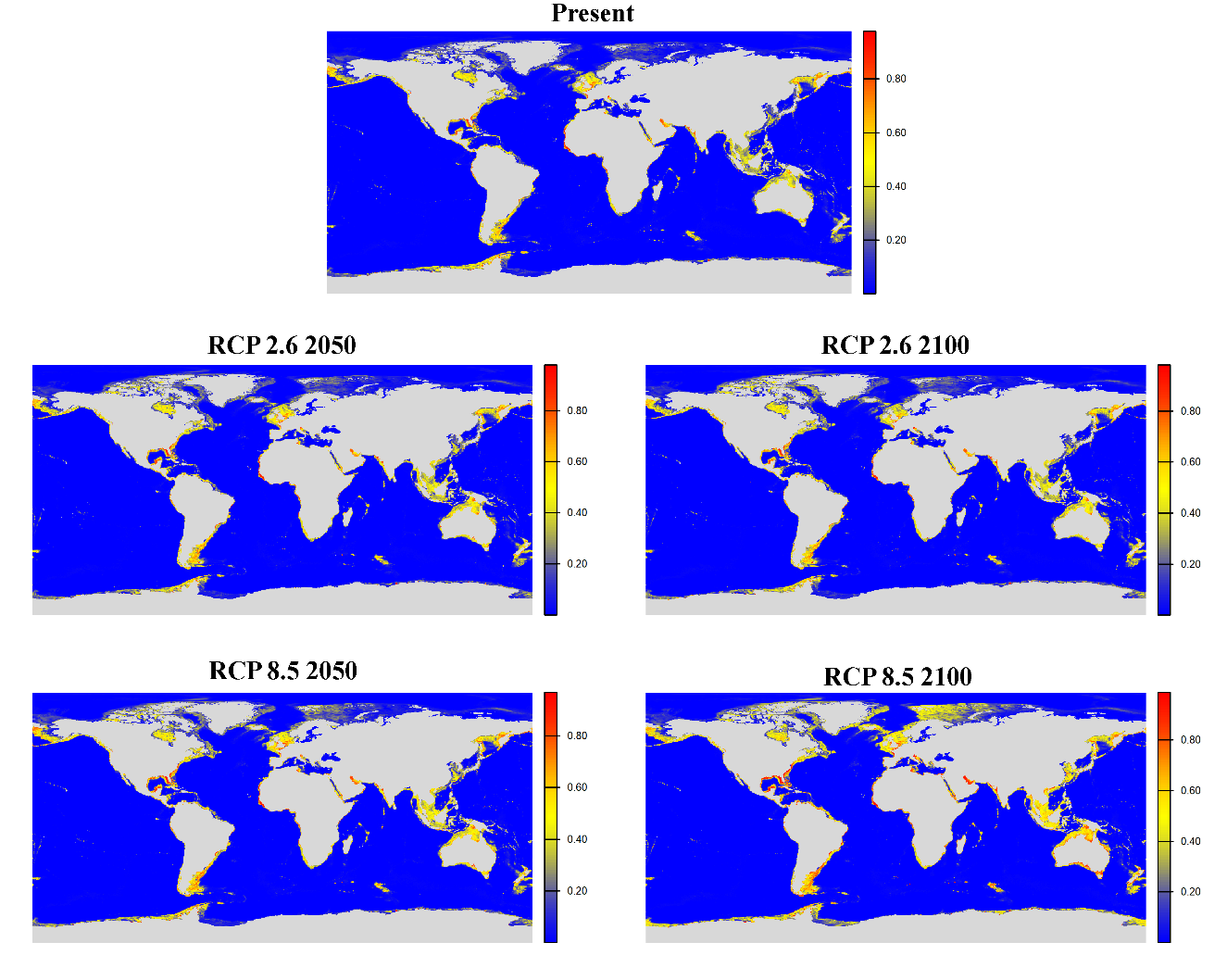

**Fig S33.** Global habitat suitability projections for *Urothoe elegans* under present and future climate scenarios. The top panel shows the current predicted distribution. Middle panels illustrate mid-century (2040–2050) and end-century (2090–2100) projections under the low-emissions scenario SSP1-2.6. Bottom panels present corresponding projections under the high-emissions scenario SSP5-8.5. Color gradients represent modeled habitat suitability (0–1), with warmer colors (yellow–red) indicating higher suitability and blue indicating low suitability.
